# Evolution of drug resistance drives progressive destabilizations in functionally conserved molecular dynamics of the flap region of the HIV-1 protease

**DOI:** 10.1101/2022.11.22.517502

**Authors:** Madhusudan Rajendran, Maureen C. Ferran, Leora Mouli, Eric Everingham, Gregory A. Babbitt, Miranda L. Lynch

## Abstract

The HIV-1 protease is one of several common key targets of combination drug therapies for human immunodeficiency virus infection and acquired immunodeficiency syndrome (HIV/AIDS). During the progression of the disease, some individual patients acquire -drug resistance due to mutational hotspots on the viral proteins targeted by combination drug therapies. It has recently been discovered that drug-resistant mutations accumulate on the ‘flap region’ of the HIV-1 protease, which is a critical dynamic region involved in non-specific polypeptide binding during invasion and infection of the host cell. In this study, we utilize machine learning assisted comparative molecular dynamics, conducted at single amino acid site resolution, to investigate the dynamic changes that occur during functional dimerization and polypeptide binding of the main protease. We use a multi-agent machine learning model to identify conserved dynamics of the HIV-1 main protease that are preserved across simian and feline protease orthologs (SIV and FIV). We also investigate changes in dynamics due to common drug-resistant mutations in many patients. We find that a key functional site in the flap region, a solvent-exposed isoleucine (ILE50) and surrounding sites that control flap dynamics is often targeted by drug-resistance mutations, likely leading to malfunctional molecular dynamics affecting the overall flexibility of the flap region. We conclude that better long term patient outcomes may be achieved by designing drugs that target protease regions which are less dependent upon single sites with large functional binding effects.

## Introduction

In the early stages of the global spread of Acquired Immune Deficiency Syndrome (AIDS), infection with Human Immunodeficiency Virus (HIV), the causative agent of AIDS, was essentially a fatal diagnosis, as there were no treatment options to combat progression to AIDS. Since then, however, an arsenal of therapeutics that target the virus have been developed. Drugs are available that target every stage of the HIV replication cycle. Recent information from the US Department of Health and Human Services HIV/AIDS medical practice guidelines list 23 individual FDA-approved HIV drugs, with an additional 23 unique combination regimens for treating HIV infection (1). These drugs target individual events in viral replication, from viral fusion and entry, to reverse transcription of the HIV RNA genome and integration into the host genome, to viral assembly, budding, and maturation into new infectious particles. For many infected patients now, therapeutic intervention has rendered the disease a chronic, managed condition with reasonable expectation of near-normal life span.

The number of people living with HIV continues to grow, as does the number of deaths due to AIDS. 38.4 million (33.9 million - 43.8 million) people were living with HIV in 2021, with a total of 650,000 (510,000 – 860,000) deaths from AIDS in 2021 (2). The widespread use of highly active antiretroviral therapy (HAART) has dramatically reduced HIV-related morbidity and mortality. For example, AIDs related death has reduced by 68% since its peak in 2004 and by 52% since 2010 (2). Yet despite the resounding success of pharmaceutical development for HIV, there remains the widely recognized problem of drug resistance. Drug resistance refers to the failure of a previously successful therapeutic intervention to maintain viral suppression within a patient. Development of resistance mutations in the viral genome undermines the efficacy of modern HIV combination therapy, and resistance monitoring is an important component of a treatment regimen (3,4). As HIV therapeutics span a range of targets in the HIV replication cycle, so too is there a wide range of mechanisms by which therapeutic resistance emerges in a treated patient. There has been tremendous effort to map the viral genomic changes that underlie viral escape from therapeutic pressure, and to comprehend the role of viral transmission in propagating emerged resistant variants. Development of resistance is an evolutionary process, as the virus in a treated host is under intense selective pressure and gains mutations that enable escape. These genomic mutations have been extensively charted, and are made available to the research community in a dedicated curated database (5). The result of mutations, though, are often altered versions of viral entities that imply a structural component to developed therapeutic escape. Given the evolutionary nature of resistance development, a comparative perspective on the mechanistic, structural and dynamic changes is potentially informative. In this work, we employ a novel molecular dynamic based comparative analysis method to probe structural dynamic aspects of the resistant drug targets.

Many structural studies of HIV drug resistance have focused on comparative analyses of the drug target structures to locate how drug binding pockets are affected (6–8). An additional consideration for probing HIV drug resistance mechanisms involves looking at not just how protein shape is altered, but also how protein dynamics are affected. Viral protein actions are key to how the virus exploits the host cell machinery, and also to how drugs bind to and change protein targets (7). Like specifics of protein shape in relation to function, protein dynamics can also be subject to evolutionary conservation (9). Our work uses machine learning-based comparative analysis tools to uncover how different forms of drug resistance impact the overall dynamics of HIV proteins in consistent ways. We use the DROIDS analysis tool, developed in previous work by our research group, to carry out proper statistical comparisons of protein molecular motions modeled via molecular dynamics (MD) simulations, focused on the class of HIV-1 protease inhibitors (PIs) (10,11). PIs were one of the first classes of HIV antivirals to benefit from concerted structure-based drug design (12), however analyses of protein dynamics that shed light on a structural aspect of viral proteins that contributes to drug resistance but have received less attention. Here we are able to demonstrate the role of adaptive evolution to alter the molecular dynamics of the flap region of the proteolytic binding pocket of HIV-1 protease in response to the selective pressure of competitive inhibitor drug therapies. These analyses provide an important contribution to our understanding of the mechanisms of HIV drug resistance, and could lead to design of future HIV therapeutics less prone to rapid emergence of viral resistance.

## Methods

### PDB structure and model preparation

Protein structures of wildtype (WT) and mutant HIV-1 protease bound and unbound to drug inhibitors were obtained from the Protein Data Bank (PDB). Summary of the different PDB structures used for the MD simulation runs are listed in table 1. After downloading the structures from the PDB database, any crystallographic reflections, ions, and other solvents used in the crystallization process were removed. Any missing loop structures in the protein structures were inferred using the SWISS-MODEL homology modelling server (13,14). Using pdb4amber (AmberTools20) hydrogen atoms were added, and crystallographic water molecules were removed (15).

**Table 1:** List of query and reference protein structures used in this molecular dynamic simulations.

| Effect of ... on MD | Query protein (PDB ID) | Reference protein (PDB ID) | References |
| --- | --- | --- | --- |
| Dimerization | HIV-1 protease dimer (6DGX) | HIV-1 protease monomer (6DGX) | (33) |
| Mutation | MDR-769 (1RPI) | WT HIV-1 protease (2PC0) | (34,35) |
| Mutation | 8mut (6OPV) | WT HIV-1 protease (6DGX) | (33,36) |
| Mutation | 6x mutant (2HB2) | WT HIV-1 protease (2HB4) | (37) |
| Drug binding | WT HIV-1 protease with Darunavir (6DGX) | Unbound WT HIV-1 protease (6DGX) | (33) |
| Drug binding | MDR-769 with Lopinavir (4L1A) | Unbound MDR-769 (4L1A) | (38) |
| Conserved dynamics | Simian SIV protease (1YTG) | HIV-1 protease (6DGX) | (31,33) |
| Conserved dynamics | Feline FIV protease (1B11) | HIV-1 protease (6DGX) | (32,33) |

### Molecular dynamic simulation protocols

For each molecular dynamic comparison (monomer vs. dimer, wildtype vs. mutant protease; protease bound and unbound to drug), large replicate sets of accelerated molecular dynamic (MD) simulations were performed. MD simulation protocol was followed as previously described, with slight modifications (16–20). In brief, for each MD comparison, large replicate sets of accelerated MD simulation were prepared and then conducted using the particle mesh Ewald method implemented on A100 and V100 NVIDIA graphical processor units by pmemd.cuda running Amber20 (15,21–24). The MD simulations were on a high performance computing cluster hosted by Rochester Institute of Technology (RIT, Rochester, NY) (25). All comparative MD analysis via our Detecting Relative Outlier Impacts in Dynamic Simulation 4.0 (DROIDS 4.0) was based upon 100 replicated sets of 1ns accelerated MD runs (i.e 100 x 1 ns MD run in each comparative state, e.g., monomer vs. dimer, wildtype vs. mutant, protease bound to drug vs. protease unbound to drug). Explicitly solvated protein systems were first prepared using tLeap (AmberTools 20), using ff14SSB protein force field, in conjunction with modified GAFF2 small molecule force field (26,27). Solvation was generated using the Tip3p water model in a 12nm octahedral water box. Charge neutralization was performed using Na+ and Cl- ions using the Ambertools20 tLeap program. Force field modifications for the small molecule ligands were generated using scaled quantum mechanical optimization via the sqm version 17 program in antechamber/Amber20 (28). For each MD comparison, an energy minimization was first performed, then heated to 300K for 300 pico seconds, followed by 10 ns of equilibration, and then finally a replicate set of 100 MD production runs was created for each comparative state. Each MD production run was simulated for 1 ns of time. All simulations were regulated using the Anderson thermostat at 300k and 1atm (29). Root mean square atom fluctuations and atom correlations were conducted in CPPTRAJ using atomicfluct and atomicorr commands (30).

### Comparative protein dynamic and statistical analyses with DROIDS 4.0

An overview of the DROIDS 4.0 pipeline using the HIV-1 protease dimer as the query protein, and the HIV-1 protease monomer as the refence protein is shown in supplemental figure 1. In brief, comparative signatures between the query and reference protein were presented as site-wise divergence in atom fluctuation. Site-wise divergences were calculated using signed symmetric Kullback-Leibler (KL) divergence calculation in DROIDS 4.0. Significance tests and p-values for the site wise differences were calculated using a two sample Kolmogorov-Smirnov (KS) test with the less conservative Benjamin-Hochberg multiple test correction. The mathematical and statistical details of DROIDS 4.0 site wise comparative protein dynamic analysis were published previously by our group (16,17,19,20). Furthermore, the code for our DROIDS 4.0 pipeline is available at our GitHub web landing: https://gbabbitt.github.io/DROIDS-4.0-comparativeprotein-dynamics/, which is also available at our GitHub repository https://github.com/gbabbitt/DROIDS-4.0-comparative-protein-dynamics.

### Identifying regions of conserved dynamics with maxDemon 2.0

An overview of the maxDemon 2.0 pipeline using the HIV-1 protease dimer as the query protein, and the HIV-1 protease monomer as the reference protein is shown in supplemental figure 2. In summary, this method trains a multi-agent learning model on the monomer and dimer dynamic states in human HIV-1 protease. The learner is then deployed upon simulations of evolutionary orthologs in unclassified functional states of dimerization (i.e. the learner attempts to classify the site-wise ortholog dynamics as either human monomer or dimer). The feature vector for the multi-agent learner is comprised of the atom fluctuations at a given site as well as atom correlations taken at 1,3,5 and 9 sites downstream on the structure. The multi-agent learner is a stacked model comprised of K-nearest neighbors, naïve Bayes, linear discriminant analysis, quadratic discriminant analysis, support vector machine, random forest, and adaptive gradient boosting algorithms employed in R language packages (base, MASS, kernlab, adaboost and randomForest). This classification is attempted at every 50 frame time slice of a 10ns simulation and the frequency of correct classification at each site is calculated. A learning profile curve is generated across all sites of the protein whereas values of 0.5 indicate learning is random and the dynamics of a particular site are indistinguishable as to whether the protease is in a monomerized or dimerized functional state. Values of 0 or 1 on the learning profile indicate 100% successful machine learning classification of the functional states (dimer = 0 monomer = 1). A canonical correlation analysis of the 7 learning profiles for each method generated on the human and ortholog dynamics and hypothesis test with Wilk’s lambda (alpha < 0.01) was used to identify regions with significantly conserved dynamics. The orthologs we used for human protease (PDB 6DGX) included several simian SIV proteases and a single feline FIV protease (PDB 1YTG, 1YTH, 1YTI, 1YTJ, and 1B11) (31,32). For more detailed information see our previous software publications and protocols (16,17,20).

## Results

### HIV-1 Protease and its inhibitors

The HIV-1 protease is a homodimeric aspartyl protease enzyme, with each monomeric subunit comprising 99 amino acid residues, and with the catalytic aspartic acid (D) residue at position 25 as part of the common triad Asp-Thr-Gly. Figure 1 shows a ribbon diagram of HIV-1 protease, with main domains indicated. The single catalytic site occurs across the two-fold symmetry axis between the two monomers, so only the dimeric form of the protein is active. An important flap domain near the top of the dimer (subunit residues 45-54) undergoes pronounced translocation to create substrate access to the active site. The protease is essential for the HIV-1 replication cycle, as it processes the GAG and GAG-POL polyproteins into the functional components needed for maturation of the intact HIV virion. Structural characterizations of the protein have highlighted key motions of the protease in its catalytic cycle (39,40). The importance of dynamics in HIV-1 protease function led us to examine further using comparative dynamics tools to assess the role of protein motion in the overall evolution of protease inhibitor drug resistance.

**Figure 1.**
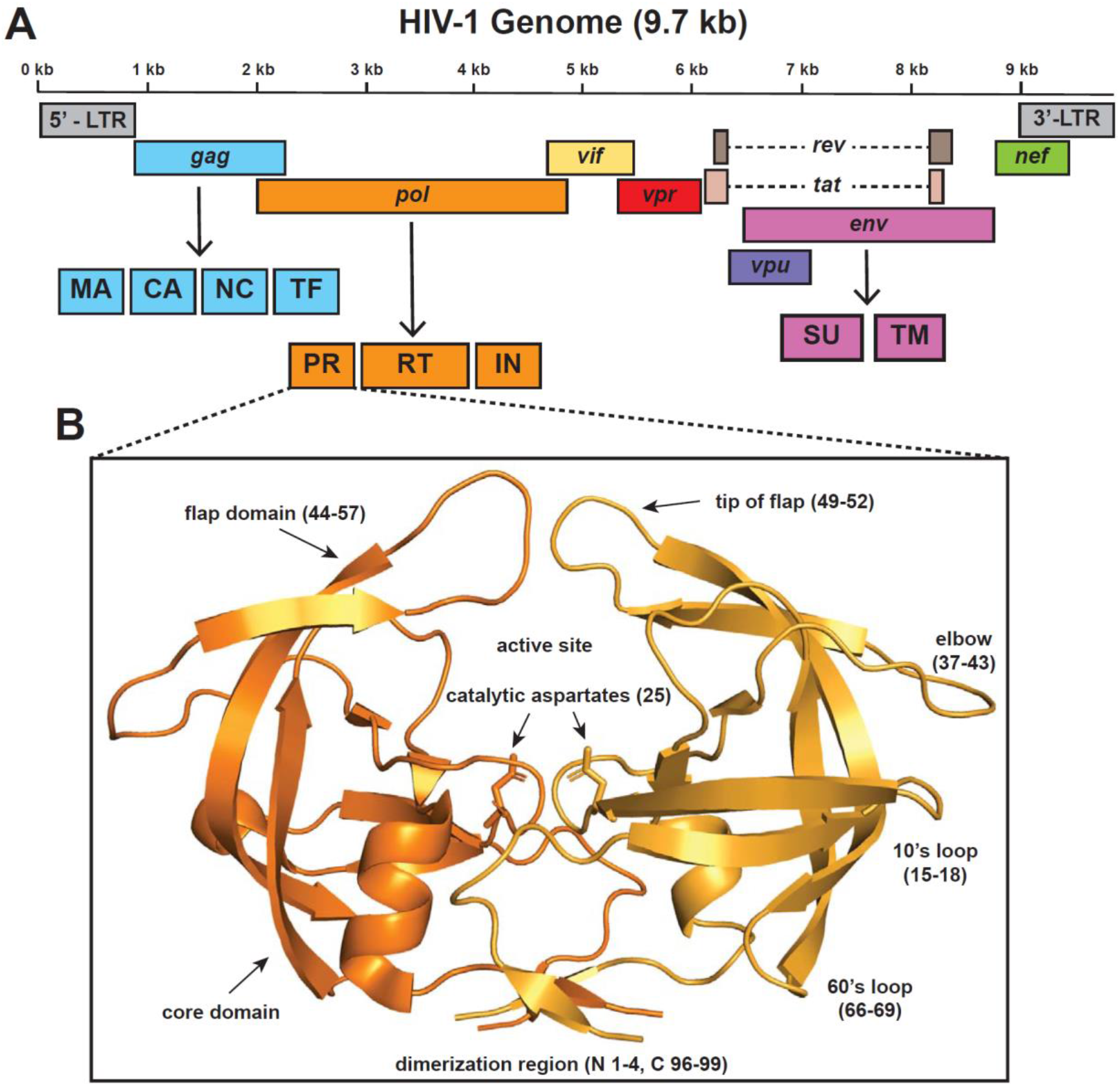
Schematic diagram of (A) HIV-1 gnome and the (B) protease dimer. **(A)** The total size of the HIV-1 genome is approximately 9.7 kb. The HIV-1 viral genes are drawn based on the RNA genome’s relative orientation. Arrow points to the cleaved protein products. The dashed line represents RNA splicing. LTR long-term repeat, Gag group-specific antigen, MA matrix protein, CA capsid domain, NC nucleocapsid, TF trans-frame protein, Pol polymerases, PR protease, RT reverse transcriptase, IN integrase, Env envelope protein, SU surface membrane protein, TM trans-membrane protein, Vif viral infectivity factor, Vpr viral protein R, Vpu viral protein U, Nef negative regulatory factor, Rev regulator of expression of viral proteins, Tat trans-activator of transcription. **(B)** The cartoon diagram of HIV-1 protease (PDB 6DGX) shows the monomers in light orange and dark orange. Key regions of the protease are labeled, and the relevant residue numbers are denoted in parenthesis.

### HIV-1 protease dimerization causes dampening of atom fluctuation in the flap region

HIV-1 protease exists as a homodimer. To investigate dimerization’s importance, we conducted comparative dynamics between the HIV-1 protease monomer and the HIV-1 protease dimer bound to a peptide ligand. In this particular MD simulation, the monomer was the reference state, and the dimer was the query state (Figure 2A). We first looked at the average flux of the dimer and monomer as a function of the amino acid position (Figure 2C). The average flux profiles for the monomer and the dimer are comparatively different, as expected.

**Figure 2.**
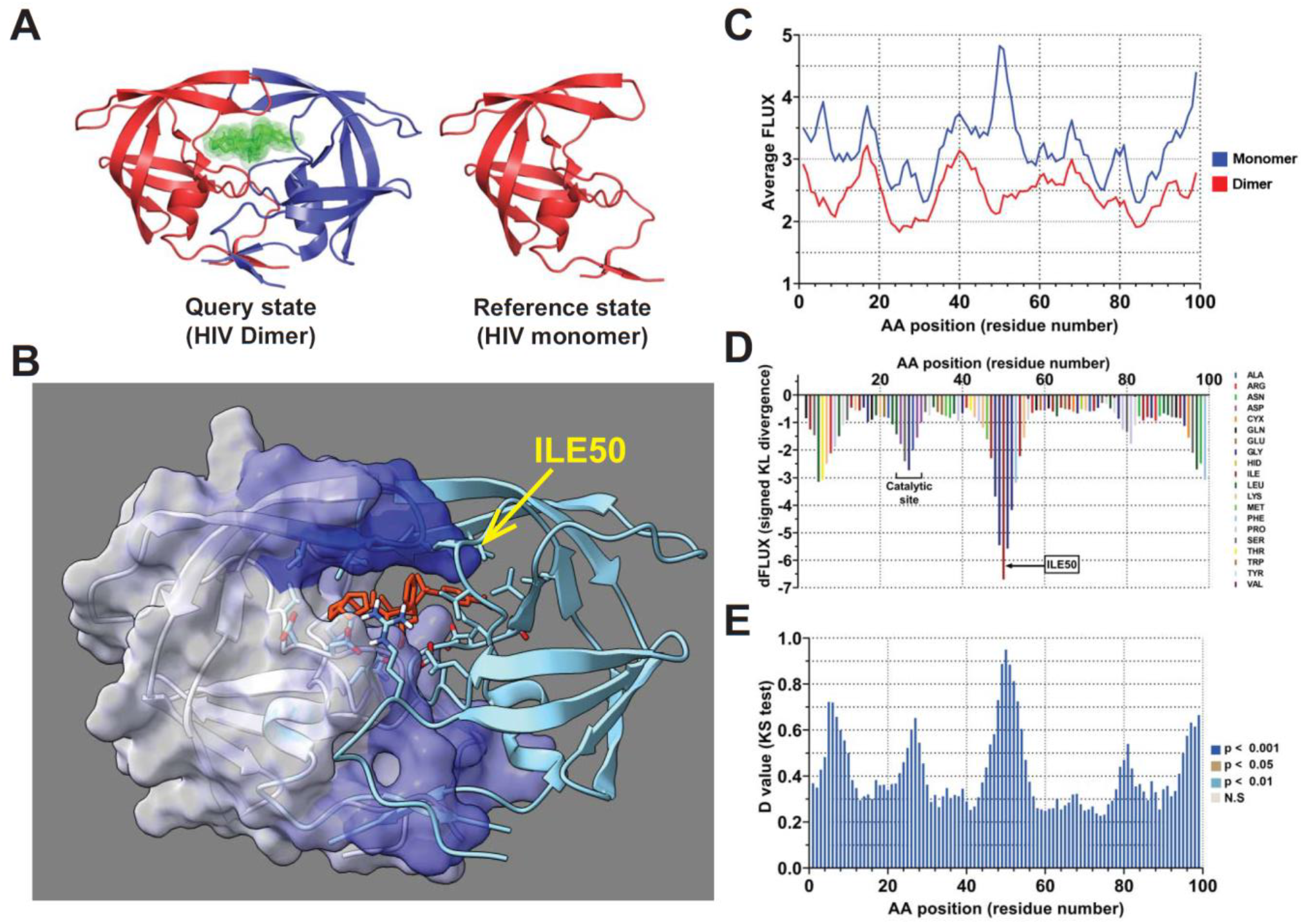
Site-wise effects of functional dimerization in HIV-1 protease on molecular dynamics. **(A)** The molecular dynamics of the HIV-1 protease dimerization function were compared by querying the peptide-bound dimer against the unbound monomer (PDB 6dgx). **(B)** A key site driving functional dimerization is isoleucine at position 50, which is double flanked by glycine (i.e., GGIGG) at the tip of the main protease “flap’. This highly solvent-exposed nonpolar hydrophobic site is highly dampened during dimerization (indicated in blue). This is also represented in the site-wise plots of **(C)** atom fluctuations in monomer and dimer, **(D)** signed KL divergence metric, and **(E)** the 2 sample KS hypothesis test as a function of amino acid position. ILE50 and catalytic site are shown in the **(D)** KL divergence plot.

Furthermore, the average flux value for the dimer is lower than the monomer, mainly due to the structure being stabilized by dimerization and existing with its native ligand. To investigate further and tease out any amino acid-specific importance, we looked at the site-wise divergence between the monomer and dimer, which was calculated using signed symmetric Kullback-Leibler (KL) divergence (Figure 2D). We observed negative signed KL divergences at all amino acid positions, indicating a universal dampening of atomic fluctuation at all sites in the dimer state. Furthermore, we observe a more substantial dampening of atomic fluctuation at the catalytic site (Asp25, Thr26, and Gly27) and Ile50 (Figure 2B, 2D). As expected, forming a proper homodimer is essential for an active catalytic site in HIV-1 protease. Other studies have also shown the importance of Ile50, with the position only tolerating a handful of amino acid substitutions (41). Ile50 residue is a part of the flap region and plays a crucial role in stabilizing the protease’s open confirmation. Lastly, we also observe a dampening of atomic fluctuation in the N-terminal residues 1-5 (figure 2D). The N-terminal residues 1-4 contribute to dimer stability in the protease (42). When the dampening of the atomic fluctuations was color-mapped to the HIV-1 protease, we observed that the strongest dampening happened across the flap regions of the dimer (Figure 2B). To further confirm that the dampening of atomic fluctuations is significant, we calculated p-values across each amino acid residue using a two-sample Kolmogorov-Smirnov (KS) test with the less conservative Benjamin-Hochberg multiple test correction. We see that the atomic fluctuation differences are significant from the N-terminus to the C-terminus of the protease (Figure 2E).

### Importance of functionally conserved dynamics in the HIV-1 protease dimerization

Functionally conserved dynamics are defined as repeated, sequence-dependent dynamics discovered after training machine learners on the functional state ensembles derived from our DROIDS pipeline. To detect functionally conserved dynamics after training and validation, an additional new MD run matching the functional reference state is simulated (i.e., matching the validation new). The learning performance of this run is compared to the MD validation run using a canonical correlation analysis conducted using all selected learners across both space and time (17). Any sequence-dependent or ‘functionally conserved’ dynamics can be recognized through a significant canonical correlation in the profile of the overall learning performance along the amino acid positions for the two similar state runs (16,17).

To identify sites of functionally conserved dynamics, a multiagent classifier comprised of up to seven learning methods (Adaboost, random forest, decision tree, linear support vector machine, radial basis function support vector machine, linear discriminant analysis, quadratic discriminant analysis, Naïve Bayes, and K nearest neighbors) was individually deployed on each site. From this, a learning performance profile across all sites on the protein is generated. We first compared the seven machine learning methods across HIV-1 protease dimer, HIV-1 protease monomer, feline immunodeficiency virus (FIV) protease monomer, and simian immunodeficiency virus (SIV) protease monomer (supplemental figure 3). In figure 3A-3D, an average classification of 0 or 1 indicated perfect learning classification at a given site, while an average learning classification of 0.5 indicates no learning was achieved at that respective site. Robust learning is observed at ILE50, a key residue of the flap region, across all dimers and monomers. As one might anticipate, we also observed strong learning in the catalytic site of the proteases (Asp25, Thr26, and Gly27). We also see near-perfect learning at the N-terminus (1–5), residues that are known to contribute to dimer stability.

**Figure 3.**
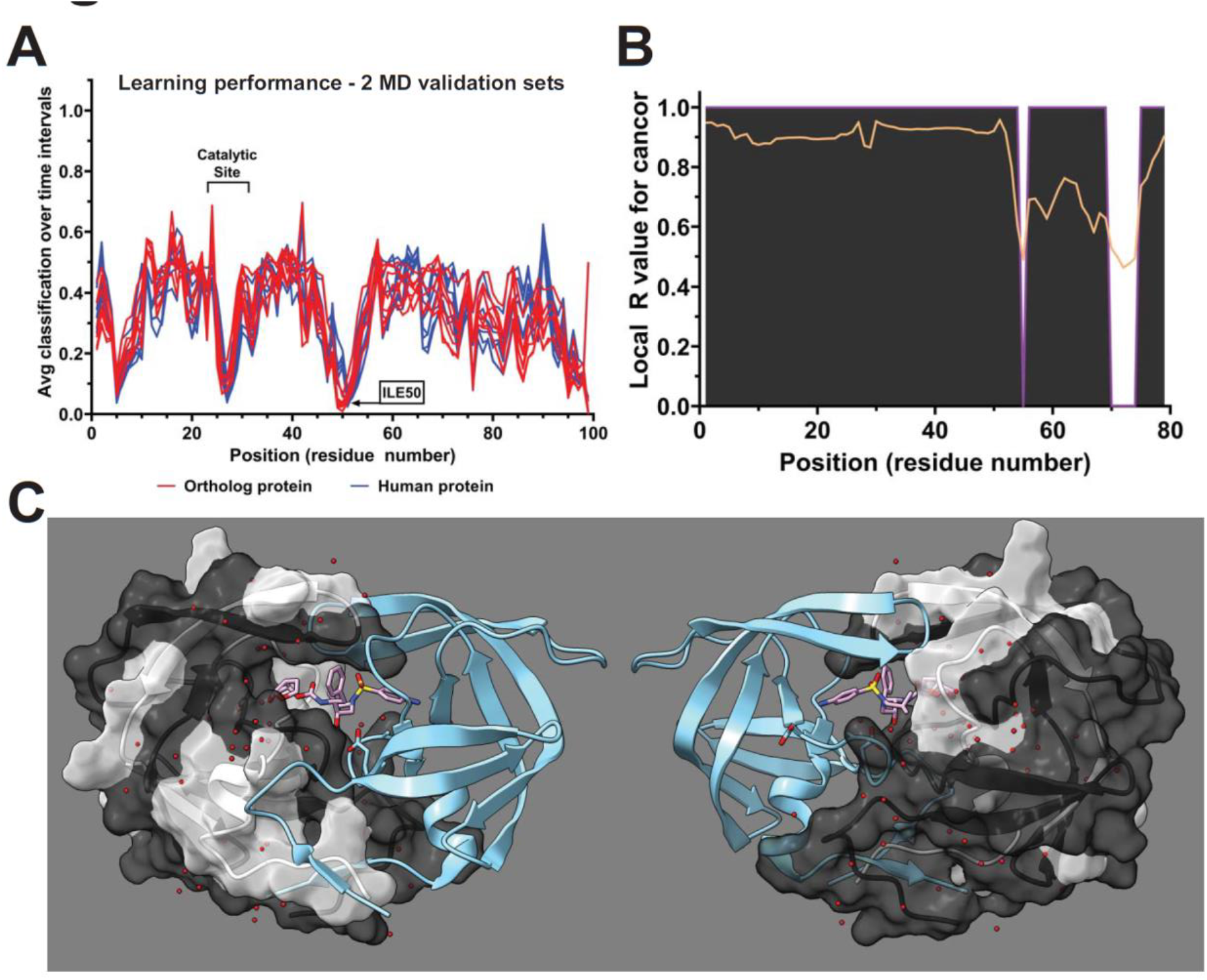
Site-wise identification of functionally conserved dynamics involved in dimerization. **(A)** highly correlated multiagent learning classification profiles for dimerized human HIV-1 protease (blue - PDB 6dgx) and simian SIV protease (red - PDB 1ytg) orthologs indicate that large regions of dynamics are functionally conserved between humans and primates. **(B)** These conserved regions are plotted in dark gray, along with the r-value from the canonical correlation analysis (orange line). **(C)** They are mapped to chain A on the human HIV-1 protease dimer.

To further investigate the importance of Ile50, we identified sites where functionally conserved dynamics are observed using a learning profile. Using the multiagent classifier, we compared the classification of HIV-1 protease and SIV protease. We observed a highly correlated multiagent learning classification for the dimerized HIV-1 and SIV protease (Figure 3A). Notably, a strong majority of amino acid residues(~90%) between the HIV-1 protease and SIV protease dimer show regions of conserved dynamics with high R values from the canonical correlational analysis (Figure 3B and 3C).

### Protease inhibitors cause the amplification of atom motions in the flap region of the HIV-1 protease

The introduction of HIV-1 protease inhibitors and their use in HAART has significantly decreased AIDS-related deaths. HIV-1 protease inhibitors usually bind to the protease’s active site and block the cleavage of viral poly-protein precursors resulting in the formation of immature protein precursors, thus forming noninfectious viral particles (43). The US FDA has approved nine protease inhibitors to treat HIV infection. However, some of the inhibitors are no longer used due to their high dosage requirements and side effects. We employed our comparative dynamics framework to investigate binding and behavior of two PIs in current clinical use: Darunavir, which was approved in 2006, and Lopinavir, approved in 2000. Both of these regimens require boosting with a low-dose of the first generation PI Rotinavir, which functions to alter metabolism of the PIs, rendering them more bioavailable.

Darunavir is one such US FDA-approved protease inhibitor with high binding affinity and can be effective against strains where resistance to other inhibitors has developed (44,45). To understand the binding interaction of Darunavir with HIV-1 protease, we did MD simulations with HIV-1 protease bound and unbound to Darunavir. To compare atomic fluctuation between HIV-1 protease bound and unbound to Darunavir, we used site-wise KL divergence along with multiple test-corrected two-sample KS tests. The more negative the KL divergence value of a specific amino acid residue, the stronger the dampening of atomic fluctuations due to the Darunavir interactions with protease. In contrast, the more positive the KL divergence value of a specific amino acid residue, the stronger the amplification of the atomic fluctuations due to the darunavir interaction with the protease. Comparative protein dynamic simulation of Darunavir bound and unbound to HIV-1 protease shows areas in the protease with amplification of atomic fluctuations (Figure 4A) and other areas where atomic fluctuation is dampened (Figure 4B).

**Figure 4.**
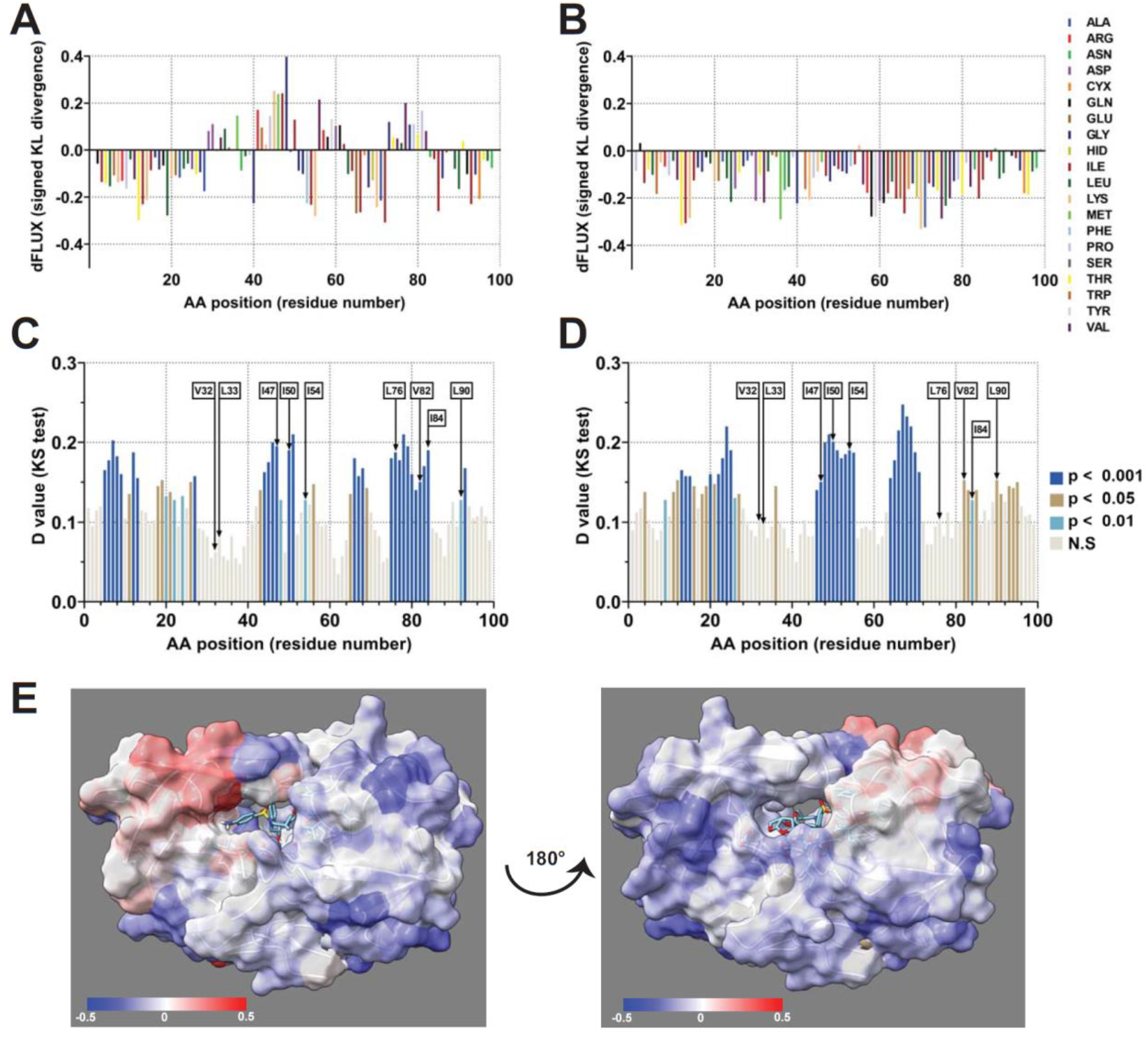
Analysis of atomic fluctuation differences due to Darunavir bound to HIV-1 protease. Sequence positional plotting of atom motion on **(A)** chain A and **(B)** chain B of HIV-1 protease dimer by Darunavir. Multiple tests corrected two-sample KS tests of significance for the difference in atomic fluctuations of **(C)** chain A and **(D)** chain B of HIV-1 protease bound to Darunavir. N.S. no significance Arrows point to the location of amino acids most common to mutate, causing resistance to Darunavir (V32, L33, I47, I50, I54, L76, V82, I84, and L90) **(E)** Changes in atom fluctuation due to darunavir binding are color mapped to HIV-1 protease (PDB 6DGX). Dark blue denotes a KL divergence value of −0.5, while red denotes a KL divergence value of +0.5. Models have been rotated 180° to show the front and back view of the protease.

Interestingly, we only see a dampening of atomic fluctuation in chain B (Figure 4B). On the other hand, we see both amplification and dampening of atomic motion in chain A of the protease (figure 4A). As mentioned previously, we also calculated significance tests and p-values for these site-wise differences. We observed significant atomic fluctuation motion in some regions of chains A and B (Figure 4C, 4D). Several of the areas of significant atomic motion correspond to the location of Darunavir resistance mutation in the protease (7). Lastly, we also color-mapped the fluctuation of the atom motion. We observed amplification of the atom motion in the flap region of the protease when Darunavir is bound to HIV-1 protease, indicating that the flap region plays a role in the binding of Darunavir (Figure 4E). Other studies have also found that binding of Darunavir alters the flap region with some studies finding that Darunavir binding exhibits a unique curling confirmation of the flap region, while others observing extended flap conformations (46,47).

Like Darunavir, Lopinavir is another HIV-1 protease inhibitor with high specificity for the protease. We wanted to investigate the importance of the flag region further. Therefore, we performed MD simulation using multi-drug resistant 769 (MDR769) strain bound and unbound to Lopinavir. Unlike our previous simulation with Darunavir, MDR769 is resistant to Lopinavir. Compared to wildtype (WT) HIV-1 protease, MDR769 has about 4.3-fold drug resistance against Lopinavir (38). Comparative MD simulation MDR769 bound and unbound to Lopinavir show amplification of atomic motion in both chains of the protease (Figure 5A, 5B). Even though both chains of MDR769 have significant residues with atomic motion fluctuation, chain B has a higher portion of significant residues (Figure 4C, 4D). Furthermore, when the atomic fluctuations were color mapped to the MDR769 protease, we see the amplification of atom fluctuations concentrated to the flap region of the MDR769, even though MDR769 is resistant to Lopinavir (Figure 5E).

**Figure 5.**
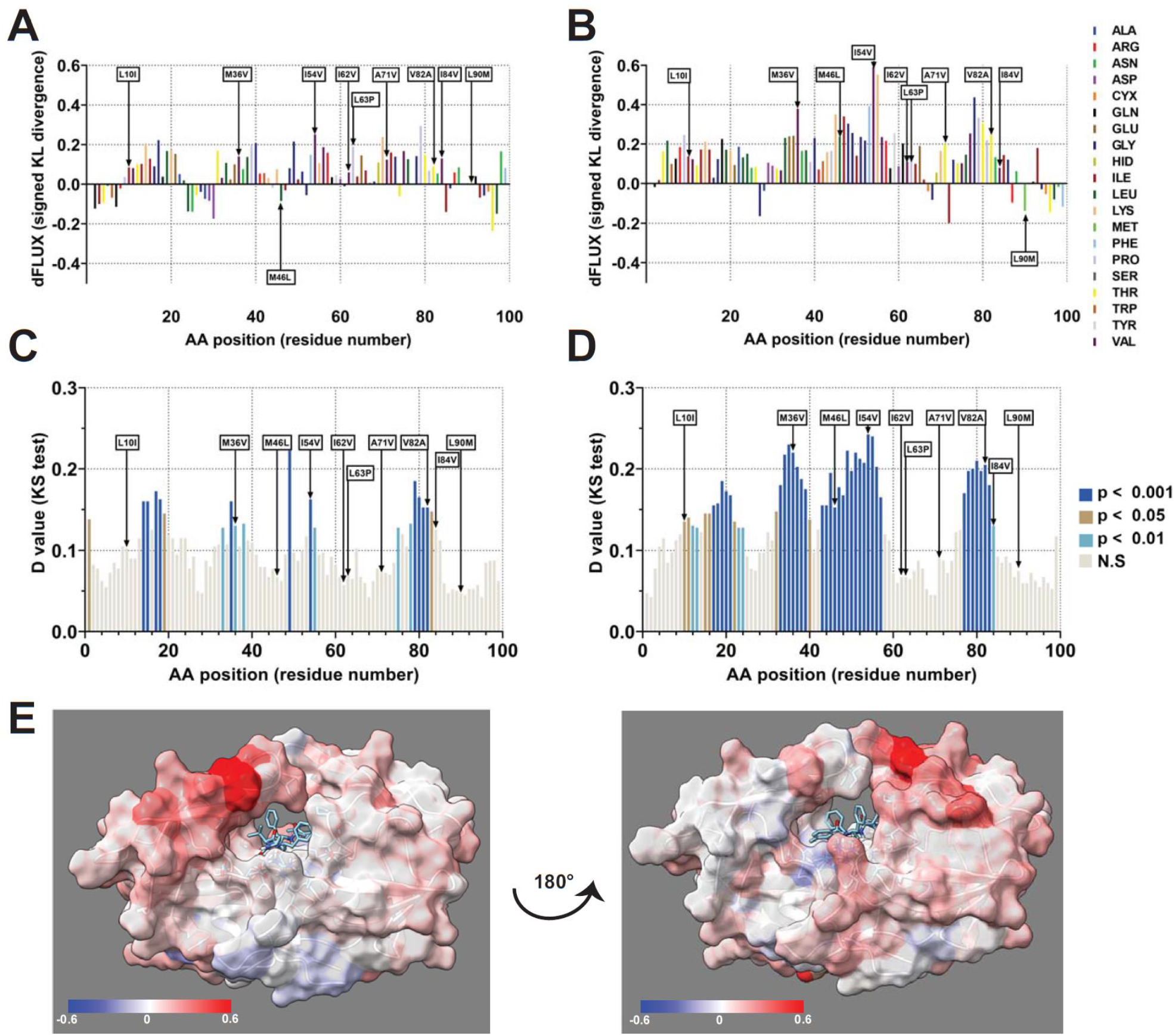
Analysis of atomic fluctuation amplification due to Lopinavir bound to MDR769. Sequence positional plotting of atom motion on **(A)** chain A and **(B)** chain B of MDR769 dimer by Lopinavir. Multiple tests corrected two-sample KS tests of significance for the difference in atomic fluctuations of **(C)** chain A and **(D)** chain B of MDR769 bound to Lopinavir. N.S. no significance. Arrows points to MDR769 mutations: L10I, M36V, M46L, I54V, I62V, L63P, A71V, V82A, I84V, and L90M. **(E)** Changes in atom fluctuation due to lopinavir binding are color mapped to MDR769 (PDB 4L1A). Dark blue denotes a KL divergence value of −0.6, while red denotes a KL divergence value of +0.6. Models have been rotated 180° to show the front and back view of the protease.

### Drug-resistant mutations impact the atom motions in the flap region of the HIV-1 protease

MDR769 is an HIV strain that has accumulated multiple drug resistance mutations in the protease, which has resulted in the decreased potency of protease inhibitors against HIV. MDR769 consists of 10 amino acid substitutions in the protease: L10I, M36V, M46L, I54V, I62V, L63P, A71V, V82A, I84V, and L90M (48). We performed similar MD simulations to understand the effect of various mutations in MDR769 and its ability to resist several protease inhibitors. In this comparative MD simulation, we used the WT protease as the reference state and MDR769 as the query state. To compare atomic fluctuations between the WT and MDR769, we used site-wise KL divergence along with multiple test-corrected two-sample KS tests (Figure 6A-6D). Even though MDR769 is a homodimer, we see several residues with significant dampening of atomic fluctuation in chain B. Of those significant residues, numerous correspond to the location of MDR769 substitution mutations (Figure 6B, 6D). The significant dampening of the atomic motions most commonly happened at the flap region of the MDR769 (Figure 6E). The dampening of the atomic motion in the flap region can be attributed to two reasons. First, the mutation of MDR769 in the flap region results in a “wide-open” structure representing an opening that is 8 Å wider than the “open” structure of the wildtype protease (49). This prevents clashing of amino acid side chains in the flap region. Next, the MDR769 is associated with a decrease in the volume of amino acid side chains with the active site cavity (34). This provides additional space for the amino acid side chains of the flap region to stabilize.

**Figure 6.**
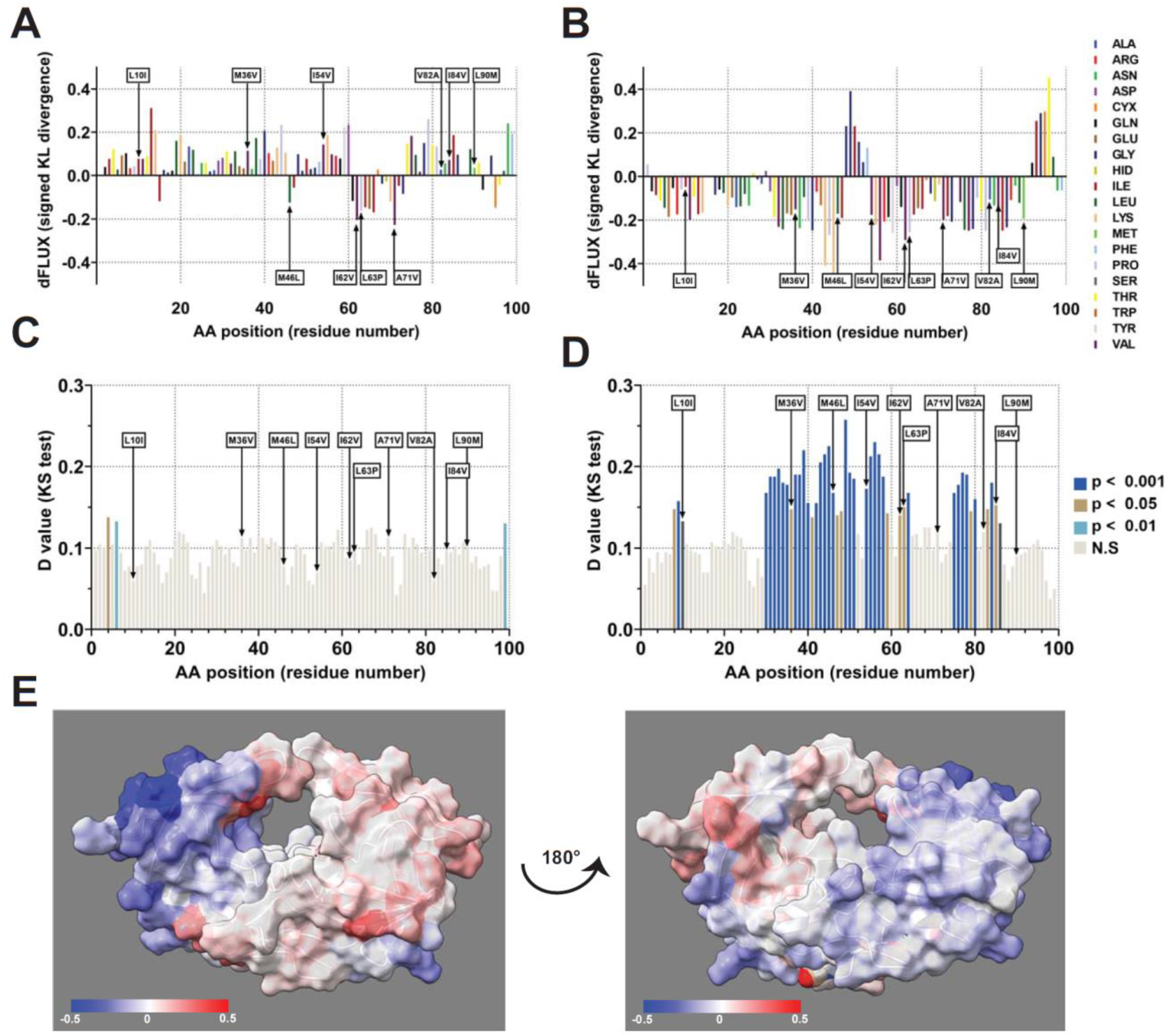
Analysis of atomic fluctuation differences due to mutational impacts of MDR769 on HIV-1 protease. Sequence positional plotting of atom motion, due to MDR769 mutational impacts, on **(A)** chain A and **(B)** chain B of HIV-1 protease. Multiple tests corrected two-sample KS tests of significance for the difference in atomic fluctuations due to MDR769 mutations of **(C)** chain A and **(D)** chain B of HIV-1 protease. N.S. no significance. Arrows points to MDR769 mutations: L10I, M36V, M46L, I54V, I62V, L63P, A71V, V82A, I84V, and L90M. **(E)** Changes in atom fluctuation due to mutational impacts of MDR769 are color mapped onto the mutant structure (PDB 4L1A). Dark blue denotes a KL divergence value of −0.5, while red denotes a KL divergence value of +0.5. Models have been rotated 180° to show the front and back view of the protease.

We also characterized two different drug-resistant HIV-1 strains. Unlike MDR769, these two strains (6x mutant and 8mut) were only resistant against the drug/inhibitor they were treated with. The 6x mutant was resistant to the broad-based inhibitor TL-3, while 8mut was resistant to Darunavir (39,50).

As the name suggests, 6x mutant encodes for six mutations in the HIV-1 protease. Those six mutations are L24I, M46I, F53L, L63P, V77I, and V82A (39). MD simulations were performed with the WT protease as the reference state and the 6x mutant as the query state. Contrary to the simulations performed comparing MDR769, we saw significant amplification of atomic motions in 6x mutant (Figure 7A-7D). This amplification was only focused on one chain of the 6x mutant (Figure 7A, 7C). When mapped onto the protease structure, the amino acid residues with significant fluctuations of the atom motion were concentrated in the flap region (Figure 7E). Heaslet et al. note that the 6x mutant protease flap regions adopt a curled region and lack the intermolecular packing interactions present in the WT protease. The authors also note that in the 6x mutant, the side chain of Ile50 and residues Gly48 and Gly49 are disordered (39). The lack of intermolecular packing interactions and disordered side chains can possibly explain the increased fluctuation of atomic motions in the mutant.

**Figure 7.**
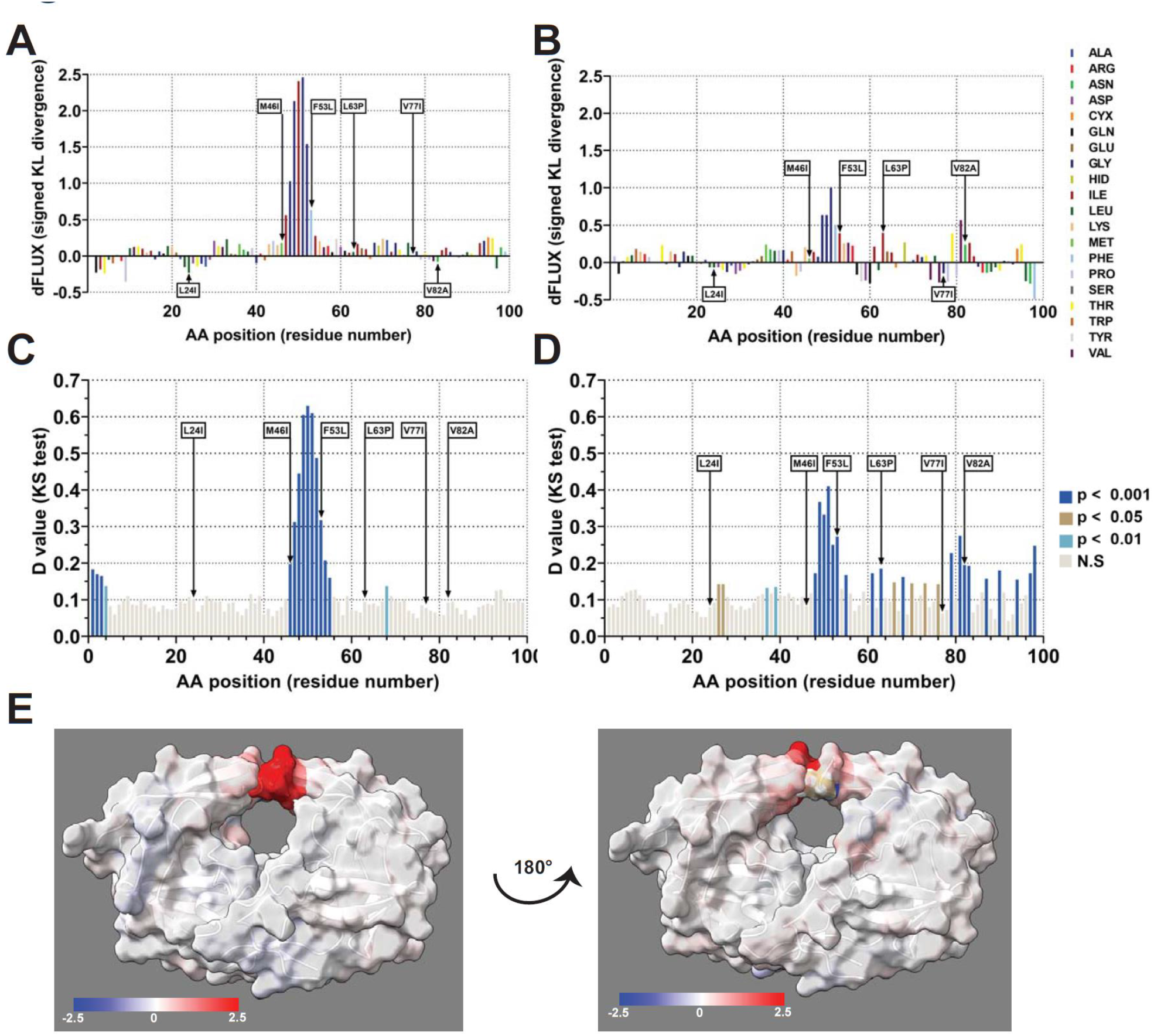
Analysis of atomic fluctuation amplification due to mutational impacts of 6x mutant on HIV-1 protease. Sequence positional plotting of atom motion, due to 6x mutant mutational impacts, on **(A)** chain A and **(B)** chain B of HIV-1 protease. Multiple tests corrected two-sample KS tests of significance for the difference in atomic fluctuations due to 6x mutant mutations of **(C)** chain A and **(D)** chain B of HIV-1 protease. N.S. no significance. Arrows points to 6x mutant mutations: L24I, M46I, F53L, L63P, V77I, and V82A. **(E)** Changes in atom fluctuation due to mutational impacts of 6x mutant are color mapped on the mutant structure (PDB 2HB2). Dark blue denotes a KL divergence value of −2.5, while red denotes a KL divergence value of +2.5. Models have been rotated 180° to show the front and back view of the protease.

Lastly, 8mut is an HIV strain that encodes eight mutations in the HIV-1 protease. Those eight mutations are I13V, G16E, V32I, L333F, K45I, M46I, V82F, and I84V (50). Similar to MDR769, comparative MD simulation of 8mut and WT show significant dampening of atomic motion in the 8mut protein (Supplemental Figure 4A – 4D). The dampening of the atom motions is prominent in one of the two chains (Supplemental figure 4A, 4C). When examined closely, similar to the two mutants discussed above, the difference in atomic motions is concentrated in the flap region (supplemental figure 4E). MD simulation using different statistical analysis by a different group have also found similar results. They found that 8mut has reduced flap fluctuations compared to the WT (50).

## Discussion

Substantial effort has gone into elucidating the important role of protein structure and motion in the function and resistance development of HIV-1 protease. Our comparative dynamics study identifies and underscores some of the key molecular motions that play a role in protease inhibitor drug resistance.

Our DROIDS analytical tool is capable of locating critical dynamics at a residue-level of specificity using site-wise divergences computed from ensembles of MD runs comparing a query and reference protein structures along with statistical tools for evaluation. The inclusion of the stacked machine learner maxDemon provides additional information on which protein regions possess functionally conserved dynamics, maintained over evolutionary time since the divergence of simian and feline lineages from that of our own. We investigated dimerization in HIV-1 protease, a process with strong experimental evidence for structure stabilization upon dimerization with concomitant motion dampening, as a confirmatory check on our methods (refs). Our study detects the global stabilization of the dimeric form, seen in signatures of damped motion of negative Kullback Leibler divergence in atom fluctuation across the entire sequence for the protease monomer compared to the dimer, with functionally important sites like ILE50 showing stronger relative compaction (see Figure 2D,E).

Analyses of drug-bound compared to unbound proteases revealed several important trends. Previous research has focused on expansion of the active site as a mechanistic driver of resistance development (51,52), and discovered the role of the substrate envelope in influencing drug resistance (53). Our analyses indicate differential behavior between the individual chains of the homodimeric protease bound to both PIs studied, an experimental result seen previously experimentally for Darunavir(54). In addition, we were able to observe in the case of Darunavir binding both localized expansive and compressive behaviors of the protease bound to drug in Chain A (Fig 4A). Lopinavir also showed both enhanced and dampened fluctuations, although overall was dominated by increased motion (especially in the flap region), consistent with the expansion of the active site expected for this multi-drug resistant example. Very few studies have been able to tease out the more nuanced picture of resistance development that extends beyond the ‘expanded pocket’ model (but see Shiek, cited above). However our method adds additional perspective provided by comparative dynamics indicating that the evolution of drug resistance is also generally accompanied by significant changes in the soft matter biophysics of this region that may not be entirely related to expansion of the shape of the binding pocket, but that still might create disruptive changes in response to competitive inhibition induced by current small molecule drugs.

Viral-host interactions are well known to drive adaptive evolution at the protein level (55,56). This has been well-documented by decades of work in comparative genomics. However, while comparative sequence analyses can easily determine local signals of natural selection acting on proteins (i.e. dN/dS type approaches, they have always been hard pressed to determine what the functional drivers of this evolution are (57,58). We developed our methods largely in response to this ‘black box’ problem in molecular evolutionary studies. Function in molecular biology is ultimately defined by both the structure and soft matter dynamics of proteins. Therefore, we have long conjectured that a proper comparative approach to molecular dynamics simulations conducted on well defined functional states (e.g. bound vs. unbound or wildtype vs. mutant) can help provide this missing perspective to the functional evolution of many protein systems. In this work, we have clearly demonstrated that adaptive evolution in the HIV-1 main protease in response to the selective pressures of drug therapy, have subsequently driven series of directional mutations that combine to alter the stability of the flap region of the proteolytic binding pocket, thereby altering its ability to function. Our work also demonstrates that while this evolution is directed, it can act over time to either stabilize or alternatively destabilize the biophysics in this region depending upon the various drug therapies involved. We speculate that the functional progression of these series of mutations that can lead to drug resistance may even depend to some degree on random chance as to whether the initial mutation(s) in this progression tended to stabilize or destabilize the flap region of the protease. Nonetheless, in concert, our overall findings suggest a common functional evolutionary route leading to most HIV drug resistance to competitive protease inhibitors. Future drug therapies that can avoid this rapid viral evolution may have more lasting benefit during the lifespan of the patient.

---

**Supplemental Figure 1.**
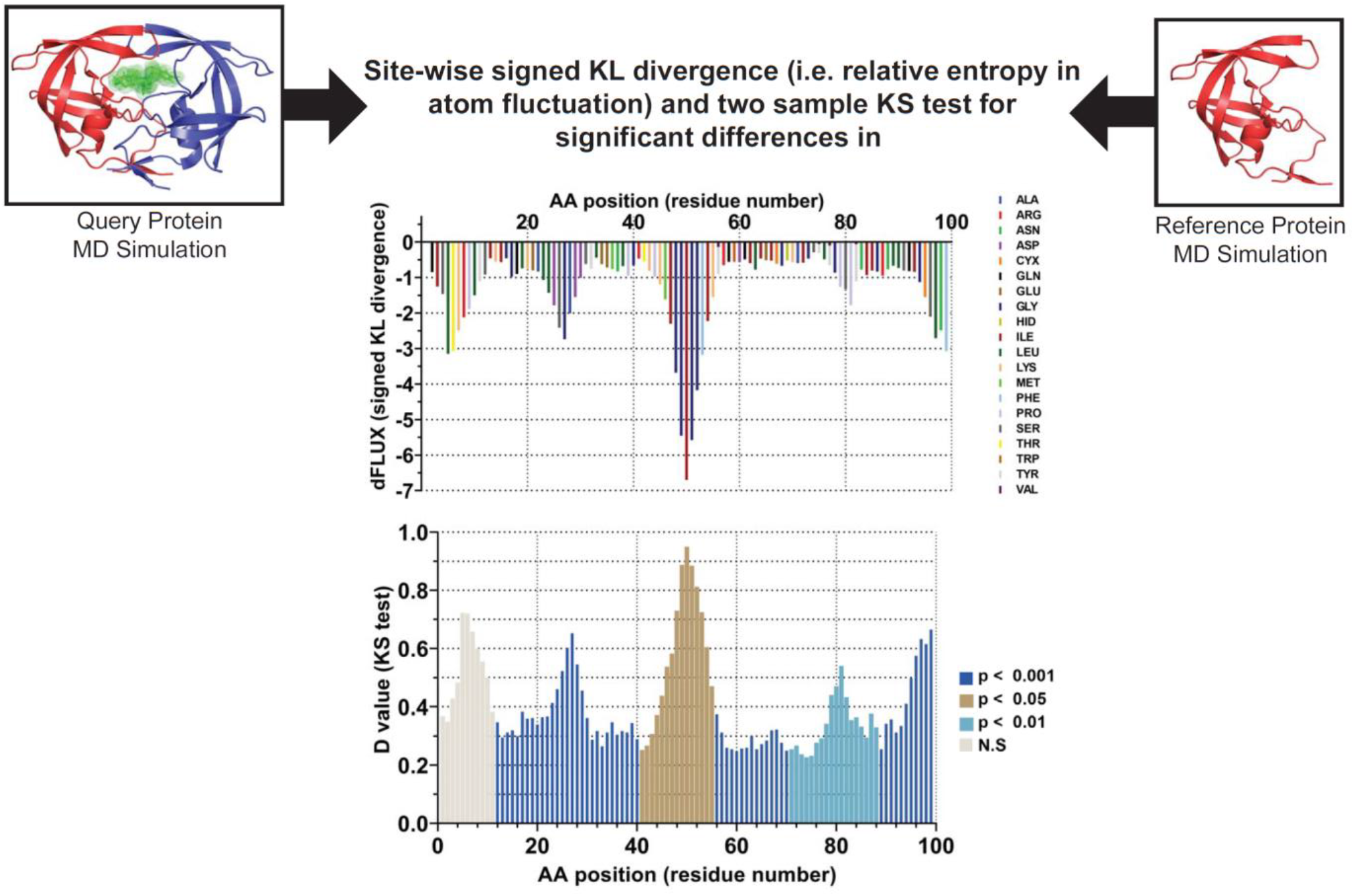
Comparative molecular dynamics with DROIDS 4.0. Our software pipeline uses a signed symmetric Kullback-Leibler divergence (i.e., also called Jensen-Shannon distance or relative entropy) to conduct site-wise comparisons of protein backbone atom fluctuations during molecular dynamics simulations representing two functional protein states (e.g., bound vs. unbound or mutant vs. wildtype). Negative values indicate the dampening of protein motion in the query state compared to the reference state, while positive values indicate amplified motions. Site-wise, two sample Kolmogorov-Smirnov tests are used to determine the statistical significance of the divergence in dynamics. The p-values are corrected via Benjamini-Hochberg to adjust the protein’s number of amino acid sites.

**Supplemental Figure 2.**
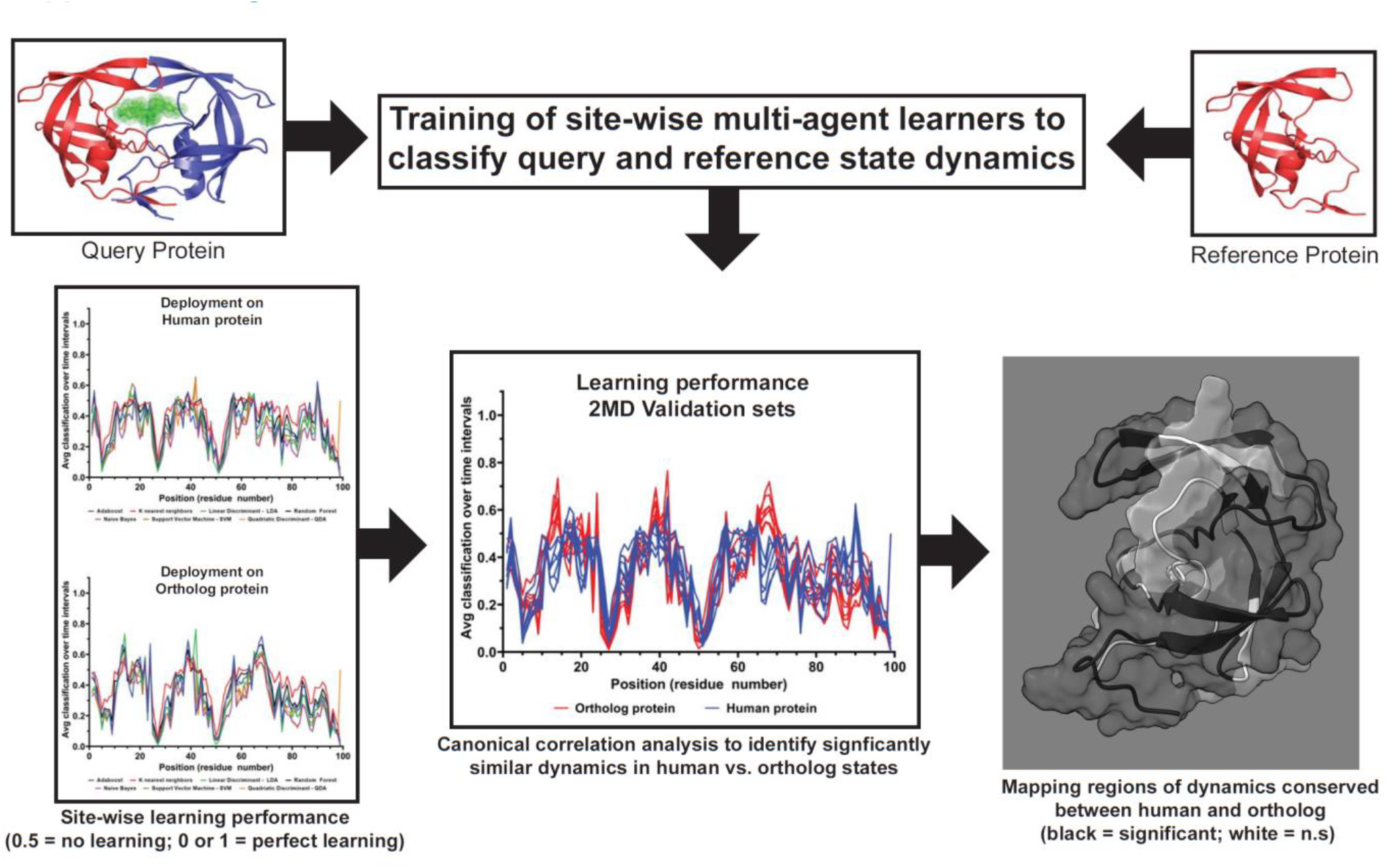
Identification of functionally conserved dynamics with maxDemon 2.0. Our software pipeline trains a multiagent or stacked machine learner classifier on features defining local atom fluctuations derived from molecular dynamics simulations in the protein’s query and reference states. The learner is deployed upon two molecular dynamics simulations of an ortholog pair (e.g., human vs. primate) in a similar but undetermined functional state. Correct and incorrect classifications are collected over sub-sampled time slices of these simulations to create two site-wise learning profiles for the ortholog pair. An average classification of 0.5 indicates no learning was achieved at a given site, while values of 0 or 1 would indicate perfect learning classification. Canonical correlation analysis in the ortholog pair learning profiles is then used to identify regions where molecular dynamics are significantly similar in the ortholog pair. This indicates that the motions in these regions were ‘functionally conserved’ throughout their evolutionary divergence from a common ancestor.

**Supplemental Figure 3.**
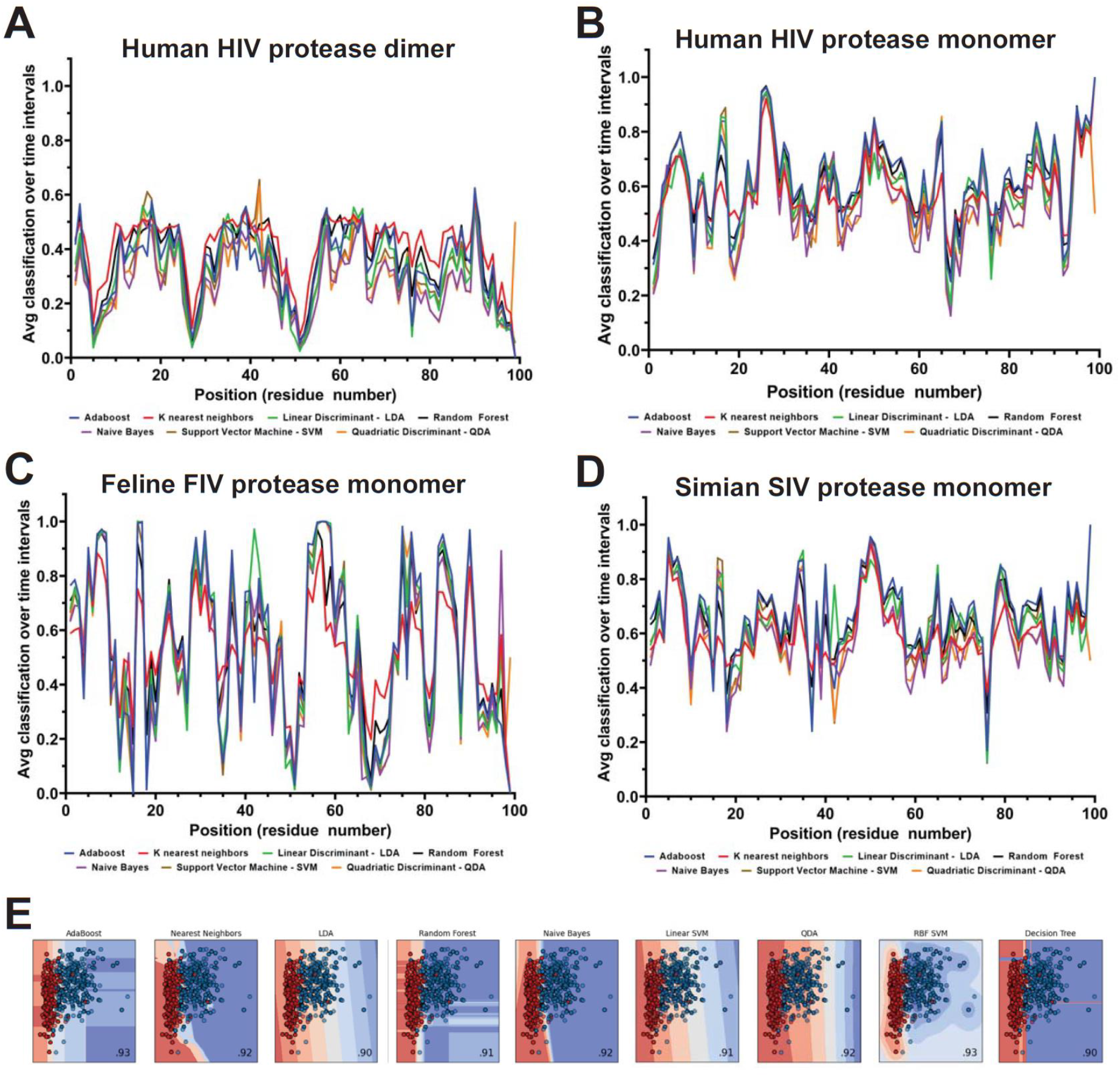
Multiagent learning profiles in human orthologs. The performance of seven machine learning methods is compared across **(A-B)** HIV-1 protease dimer and monomer (PDB 6DGX) and **(C)** feline immunodeficiency virus (FIV) protease monomer (PDB 1B11) and (D) simian immunodeficiency virus (SIV) protease monomer (PDB 1YTI). An average classification of 0.5 indicates no learning was achieved at a given site, while values of 0 or 1 would indicate perfect learning classification. Strong learning at Ile50 is easily observed when deployed upon the dynamics of the protease monomers and dimers. (E) The learning space on reduced local atom fluctuations for seven machine learning methods is also shown.

**Supplemental Figure 4.**
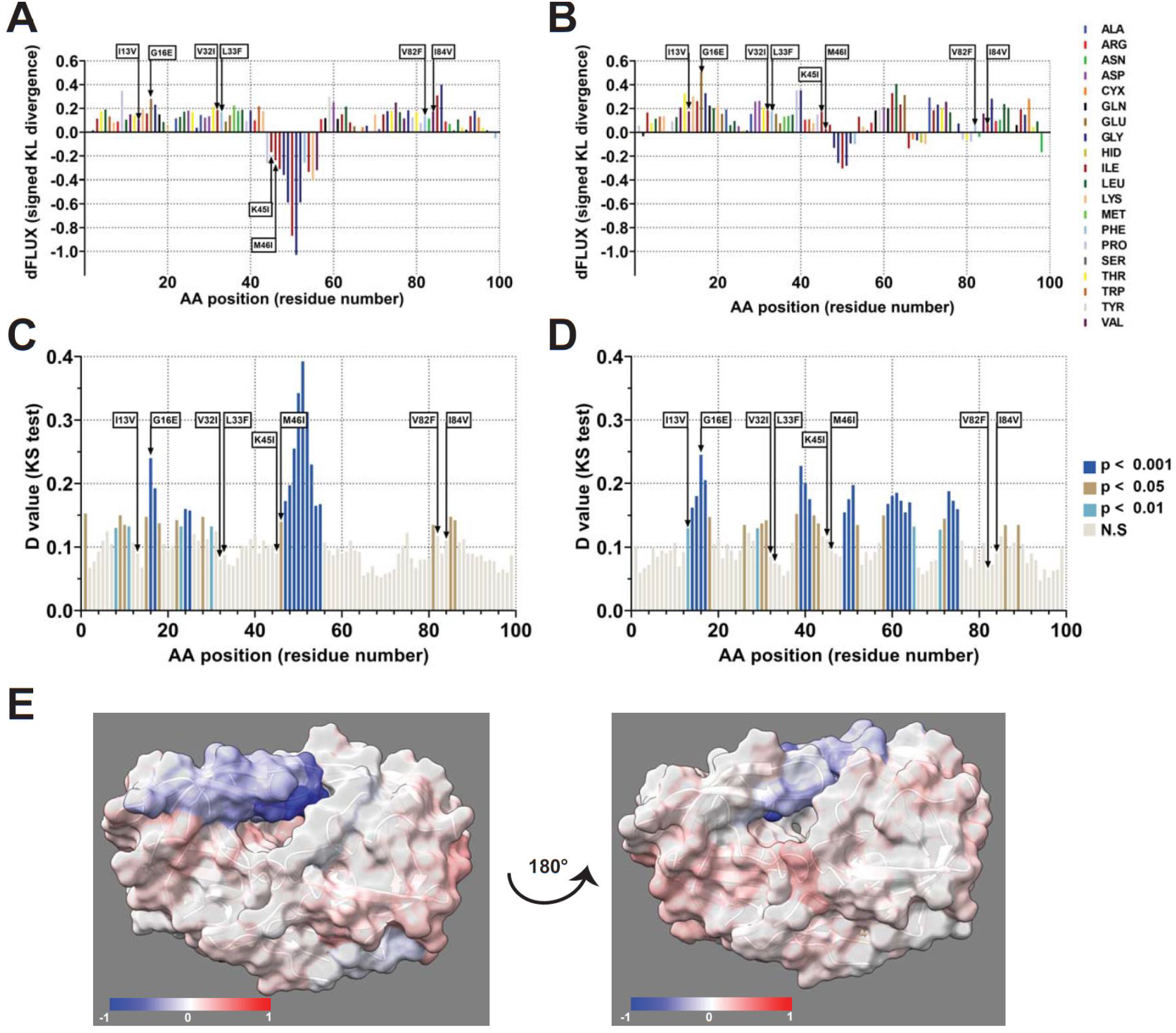
Analysis of atomic fluctuation amplification due to mutational impacts of 8mut mutant on HIV-1 protease. Sequence positional plotting of atom motion, due to 8mut mutant mutational impacts, on **(A)** chain A and **(B)** chain B of HIV-1 protease. Multiple tests corrected two-sample KS tests of significance for the difference in atomic fluctuations due to 8mut mutant mutations of **(C)** chain A and **(D)** chain B of HIV-1 protease. N.S. no significance. Arrows points to 6x mutant mutations: I13V, G16E, V32I, L33F, K45I, M46I, V82F, and I84V. **(E)** Changes in atom fluctuation due to mutational impacts of 8mut mutant are color mapped onto the mutant structure (PDB 6OPV). Dark blue denotes KL divergence value of −1, with red denoting KL divergence value of +1. Models have been rotated 180° to show the front and back view of the protease.

