## Supplementary figures and images for "Evolution of drug resistance drives progressive destabilizations in functionally conserved molecular dynamics of the flap region of the HIV-1 protease"

### DROIDSplot_10runs.pdf

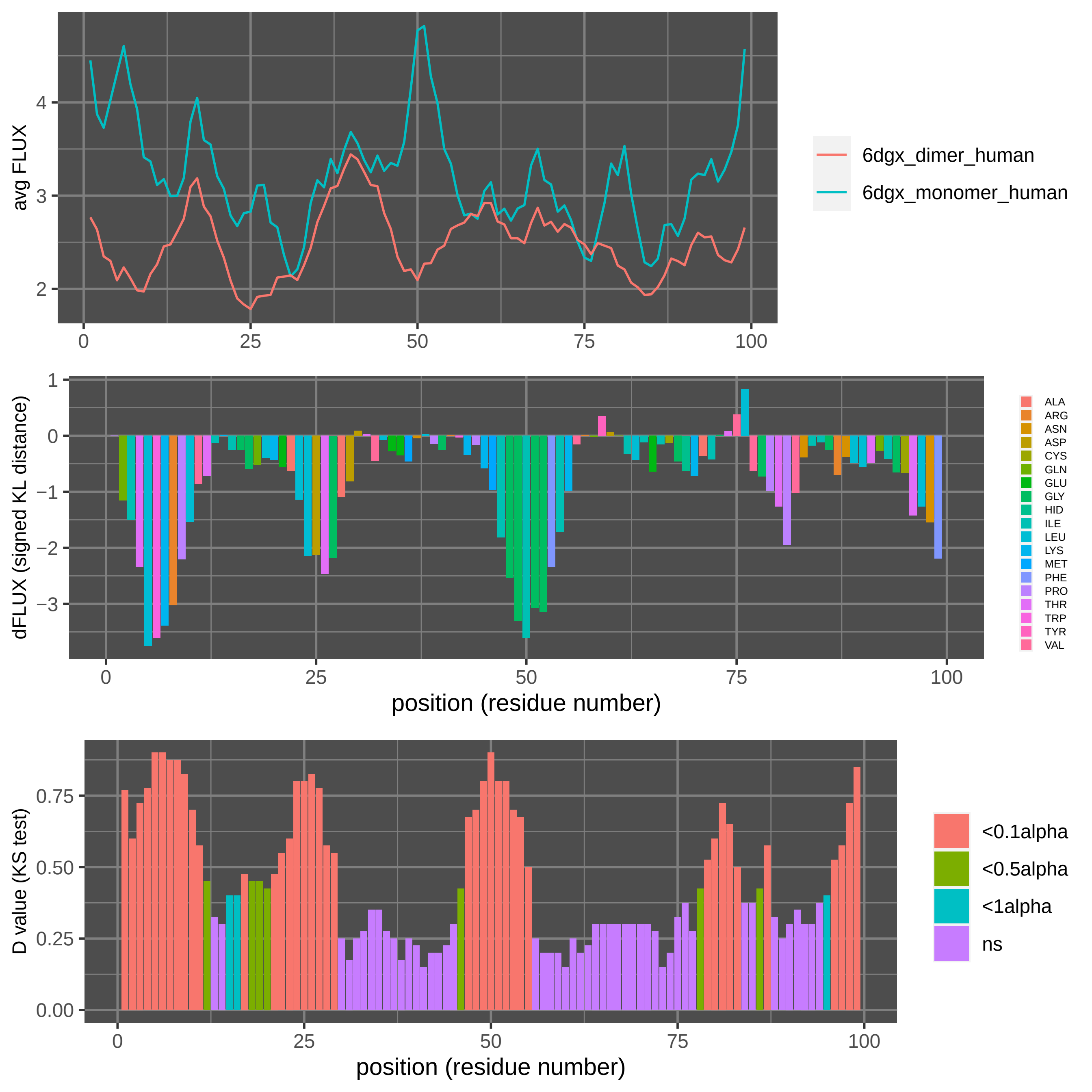

### DROIDSplot_100runs.pdf

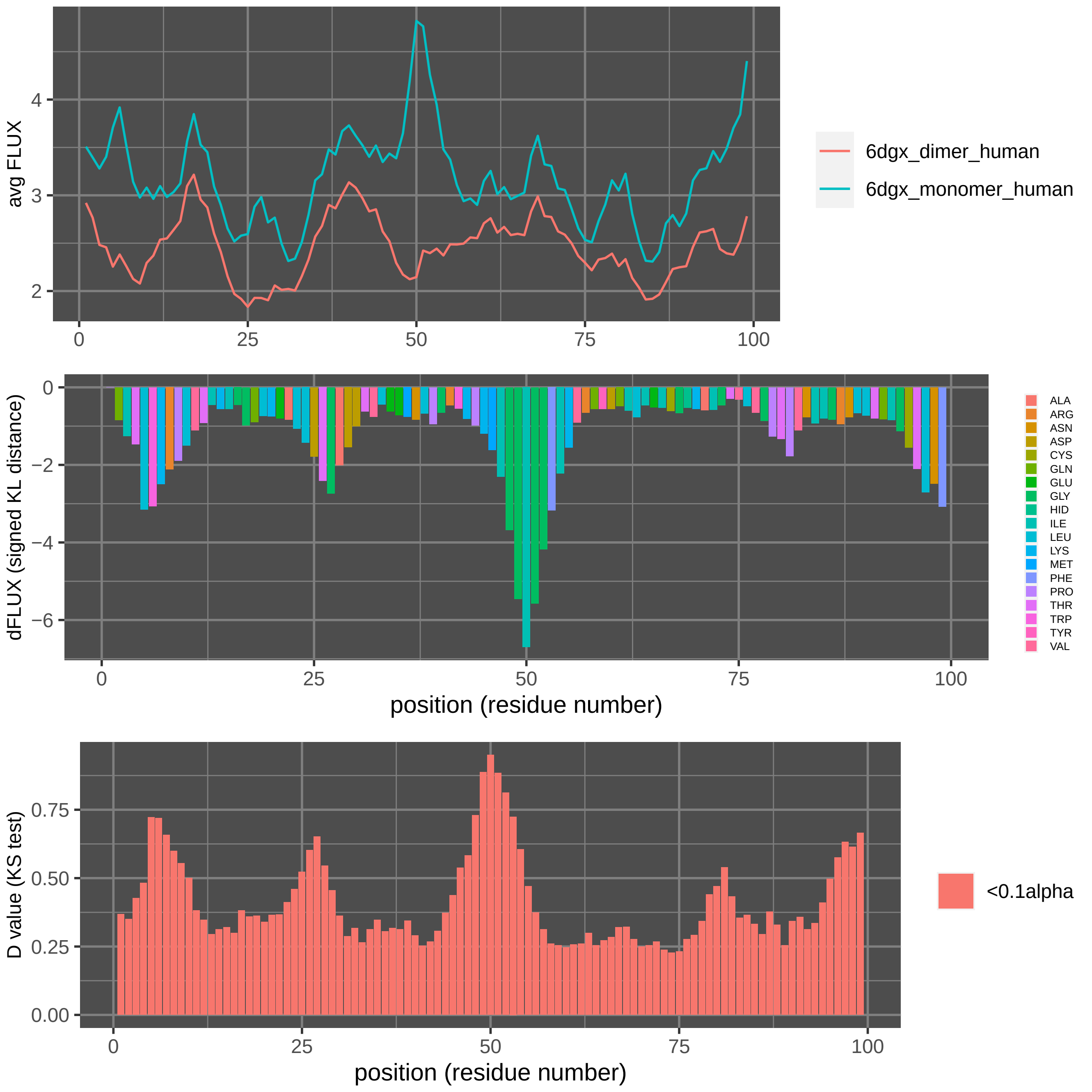

### fluxCorr_learningPLOT_1b11_monomer_feline.pdf

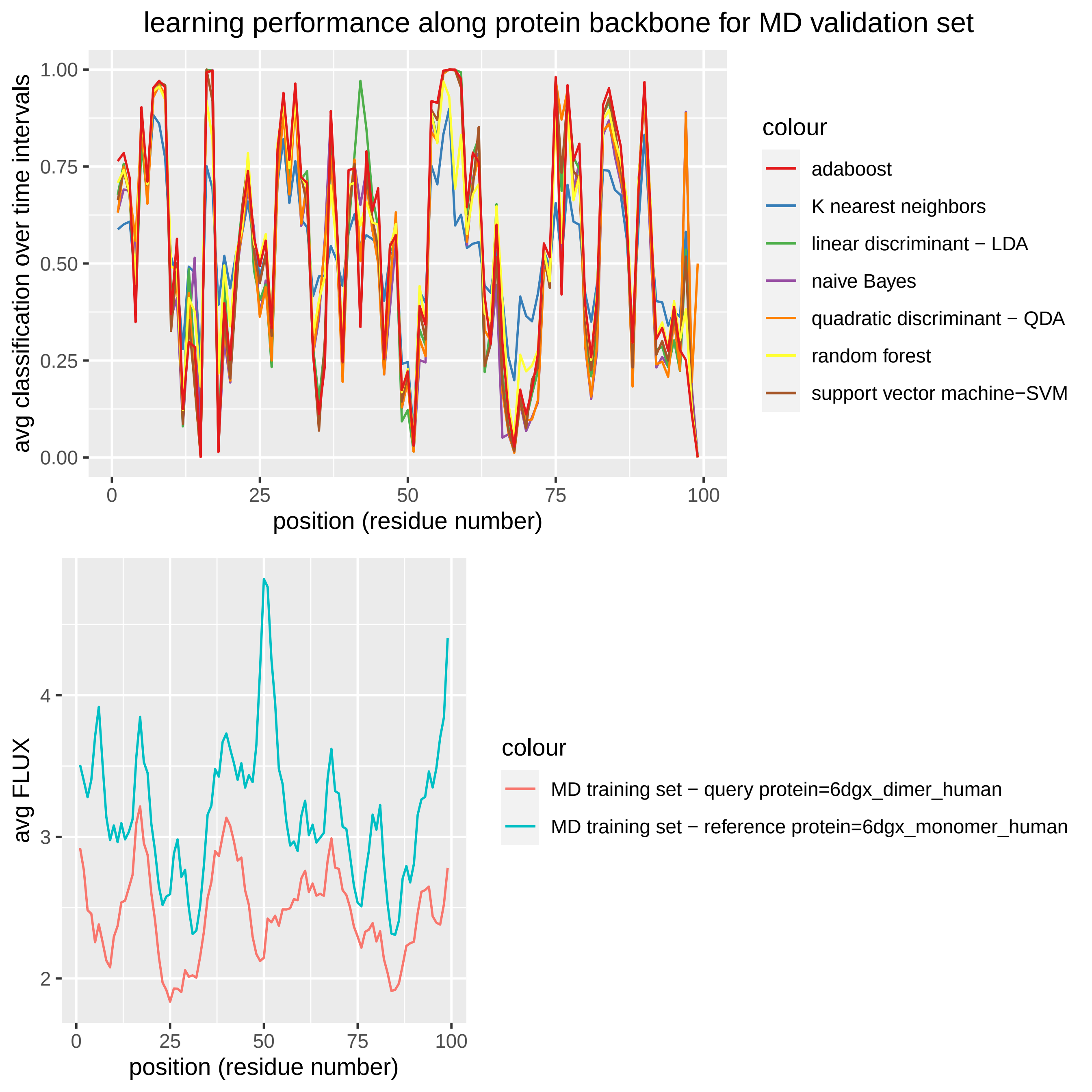

### fluxCorr_learningPLOT_1yti_monomer_simian.pdf

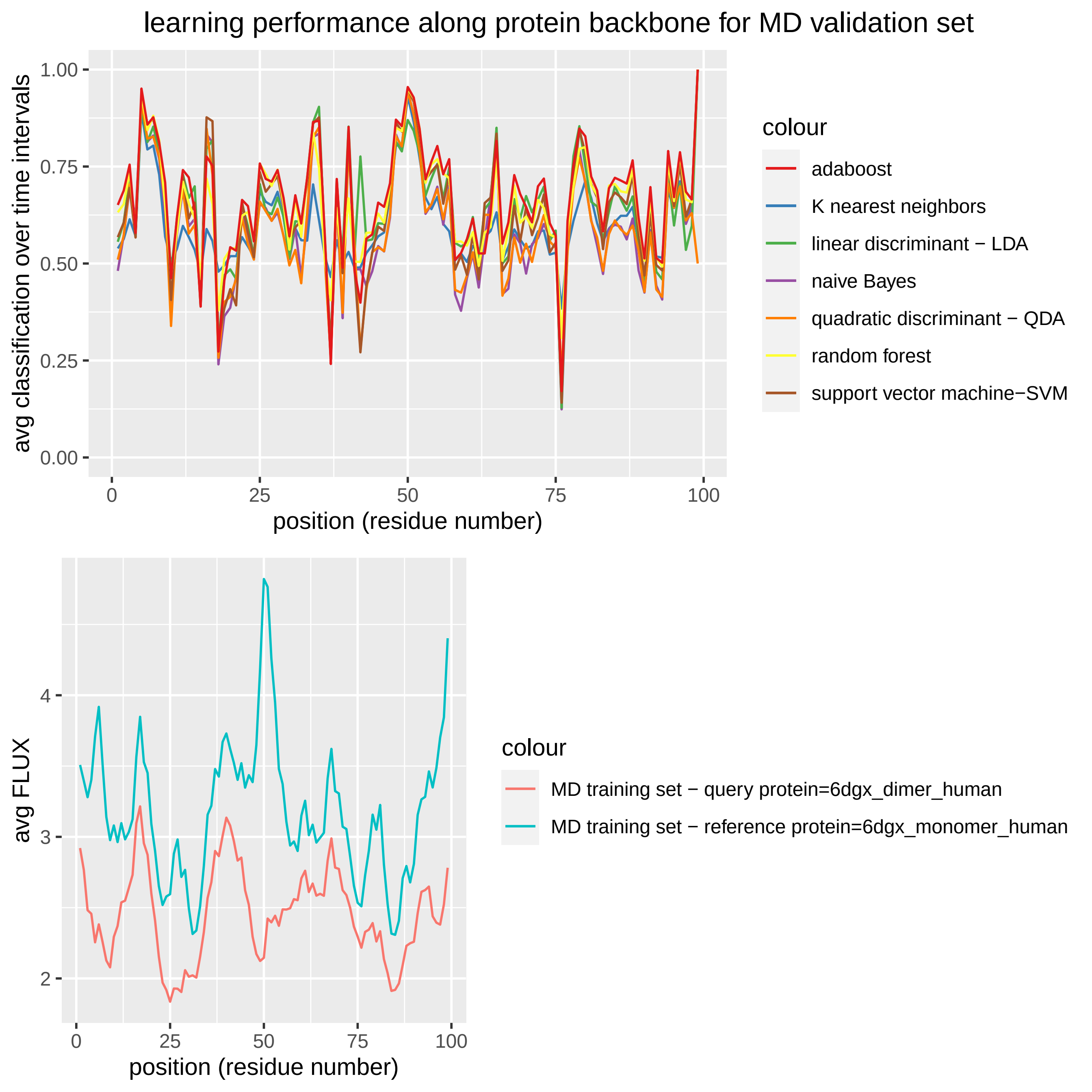

### fluxCorr_learningPLOT_1ytj_monomer_simian.pdf

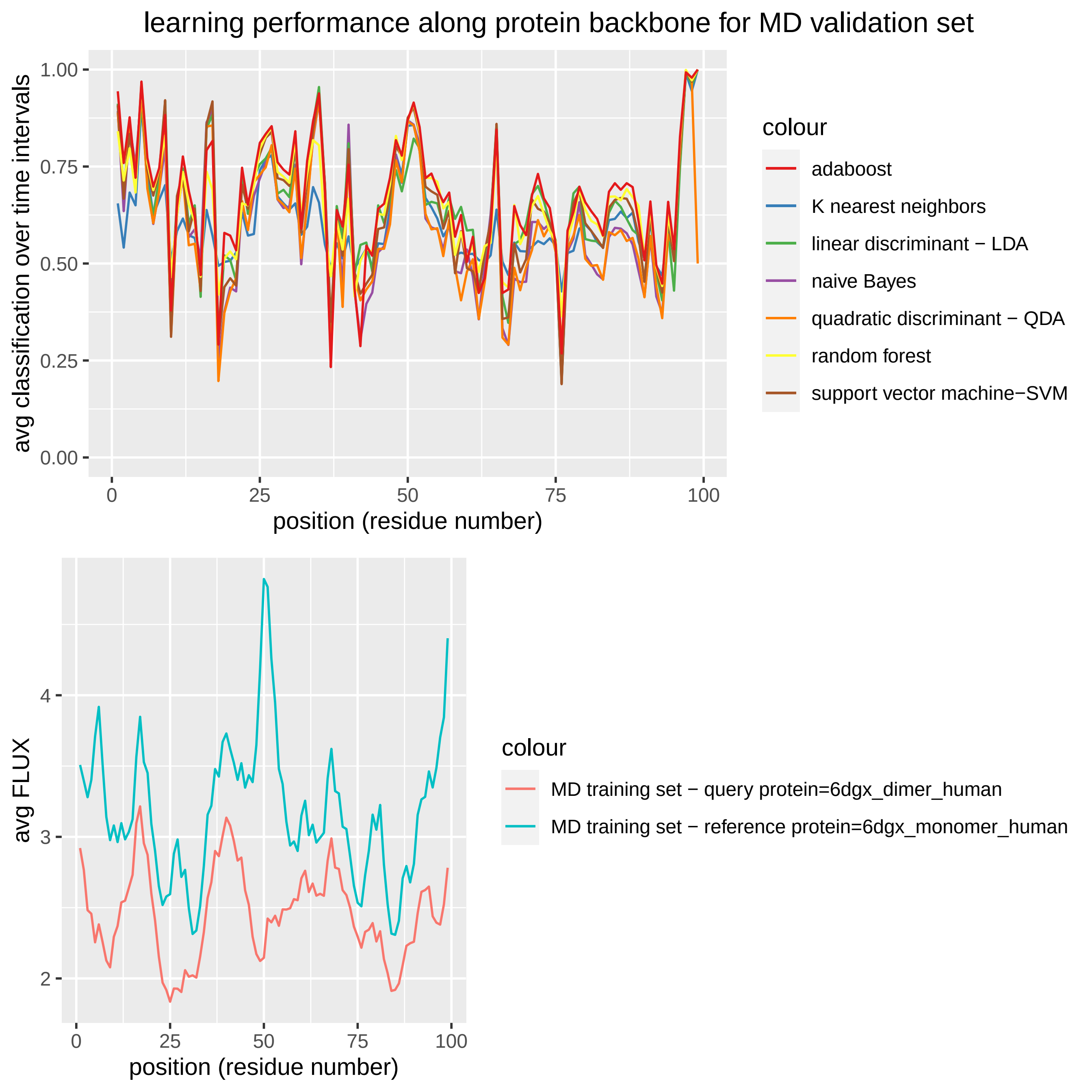

### fluxCorr_learningPLOT_6dgx_monomer_human_1.pdf

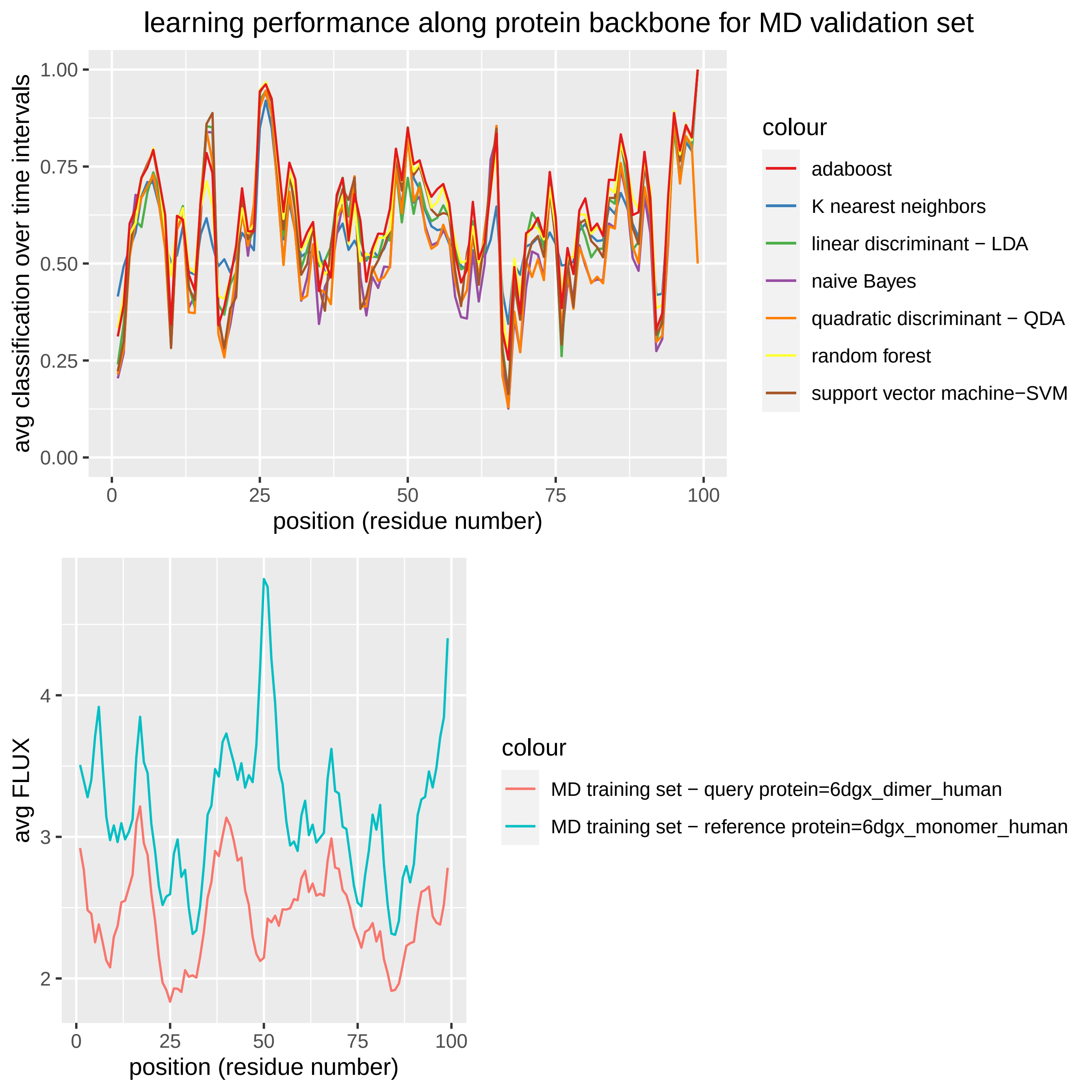

### fluxCorr_learningPLOT_6dgx_monomer_human_2.pdf

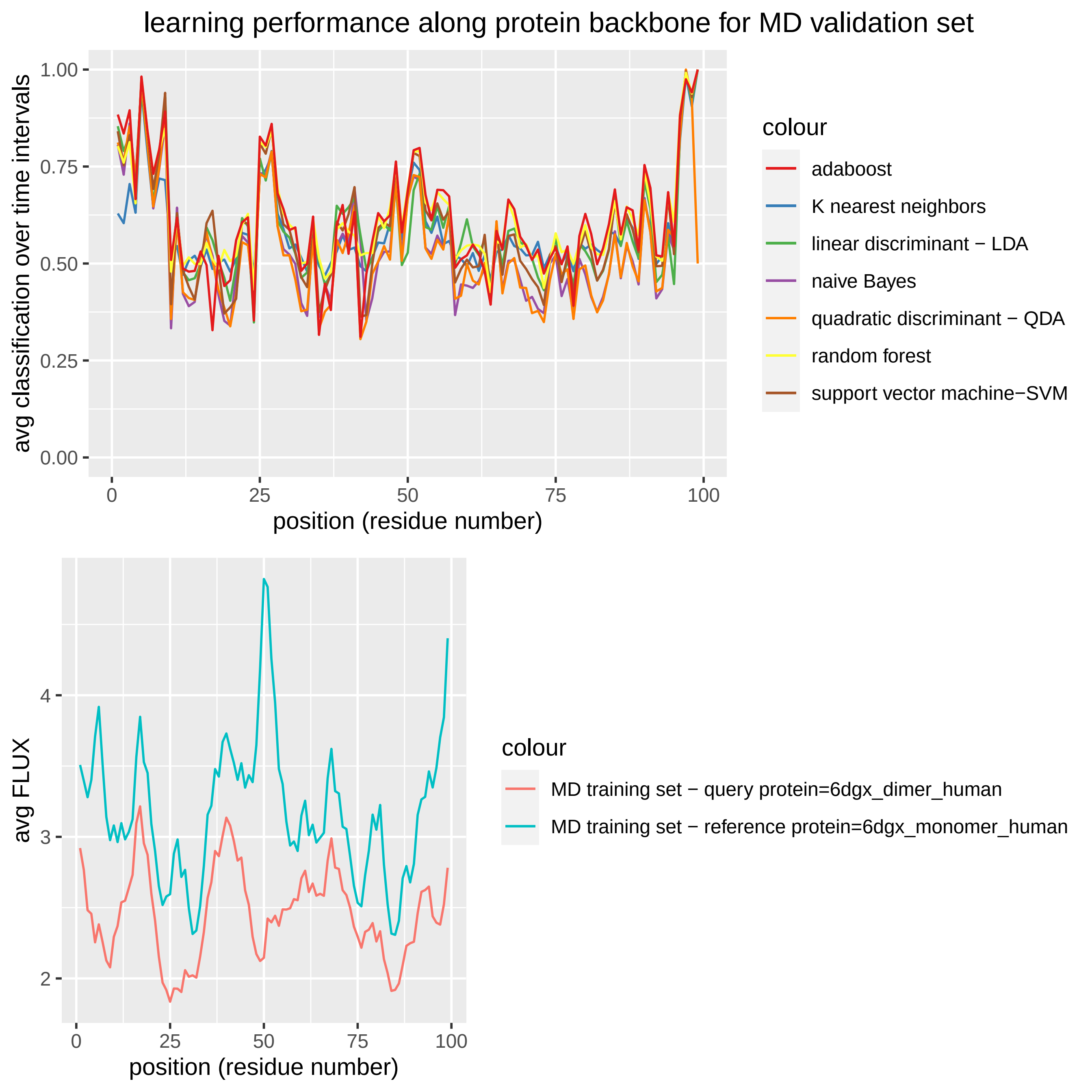

### fluxCorr_learningPLOT_6dgx_nopep_human.pdf

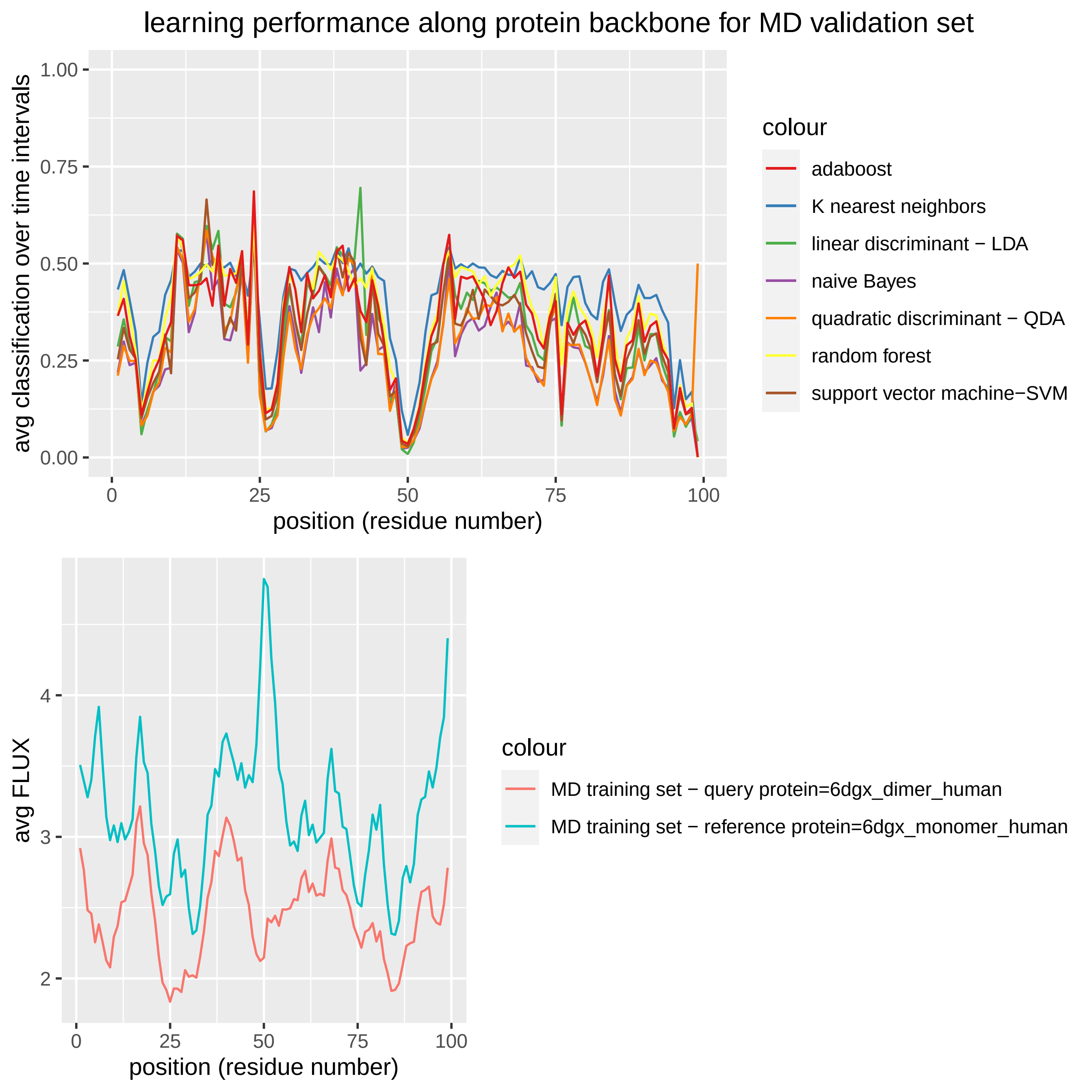

### fluxCorr_learningPLOT_6dgx_nopep_human_1.pdf

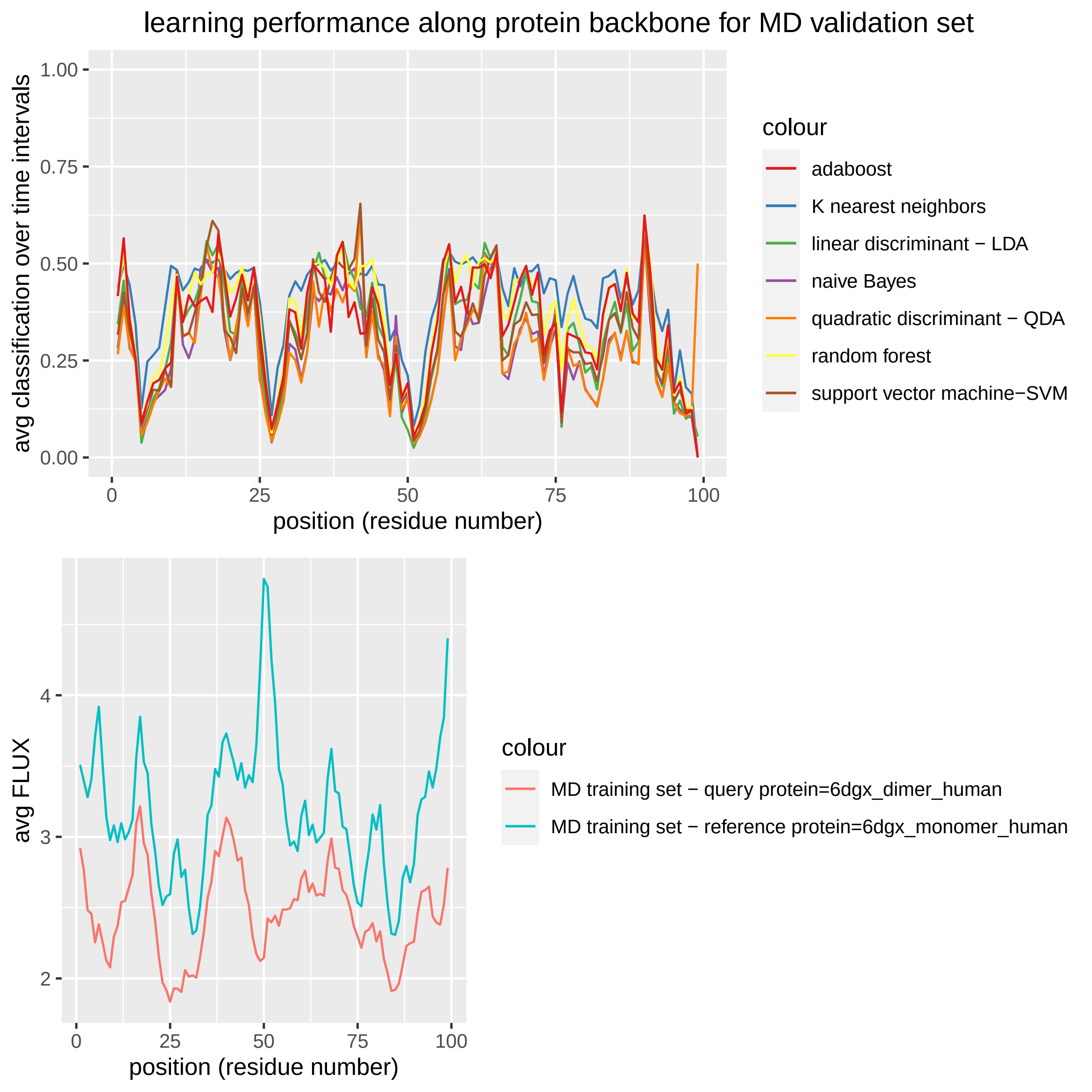

### fluxCorr_learningPLOT_6dgx_nopep_human_2.pdf

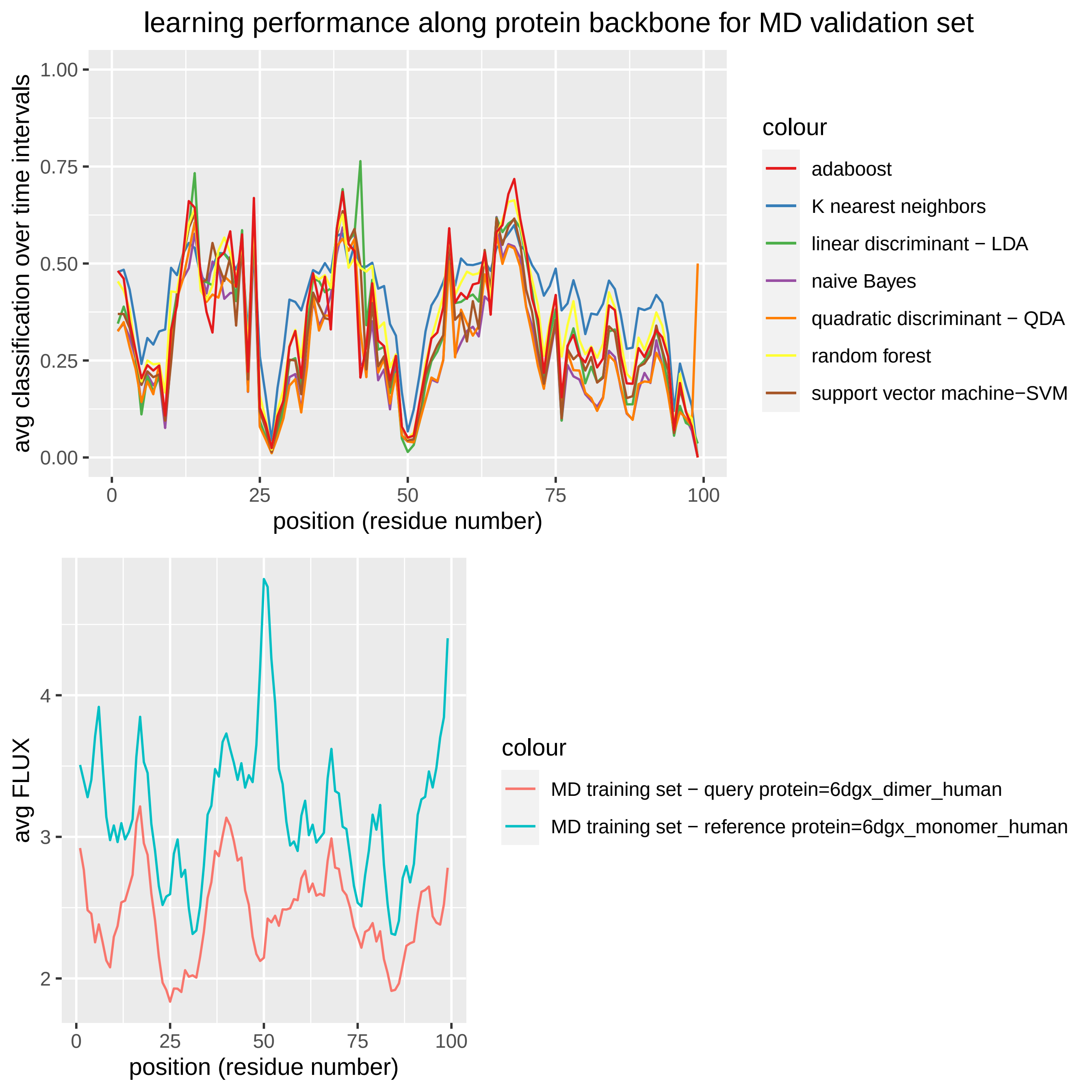

### image1.png

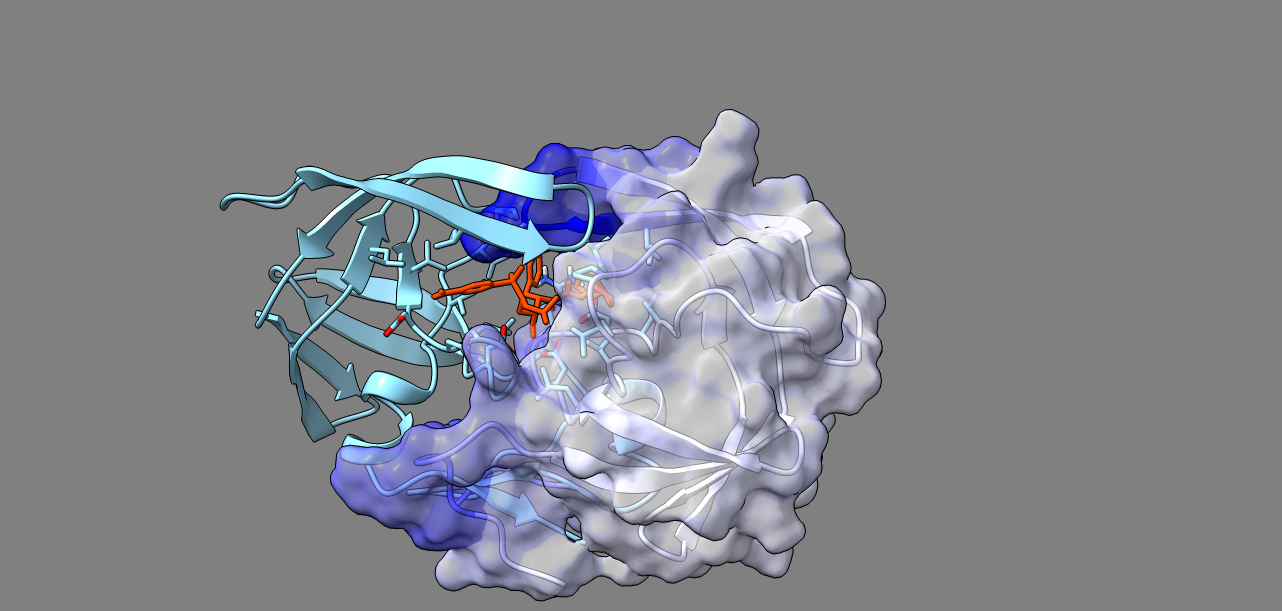

### image1.png

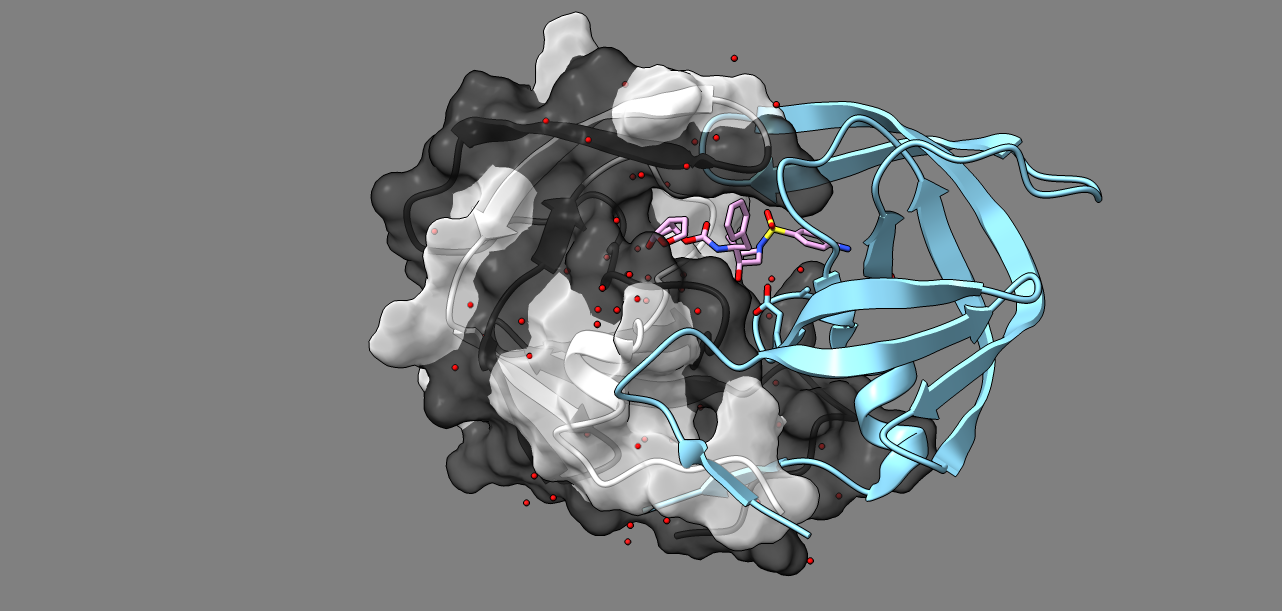

### image1.png

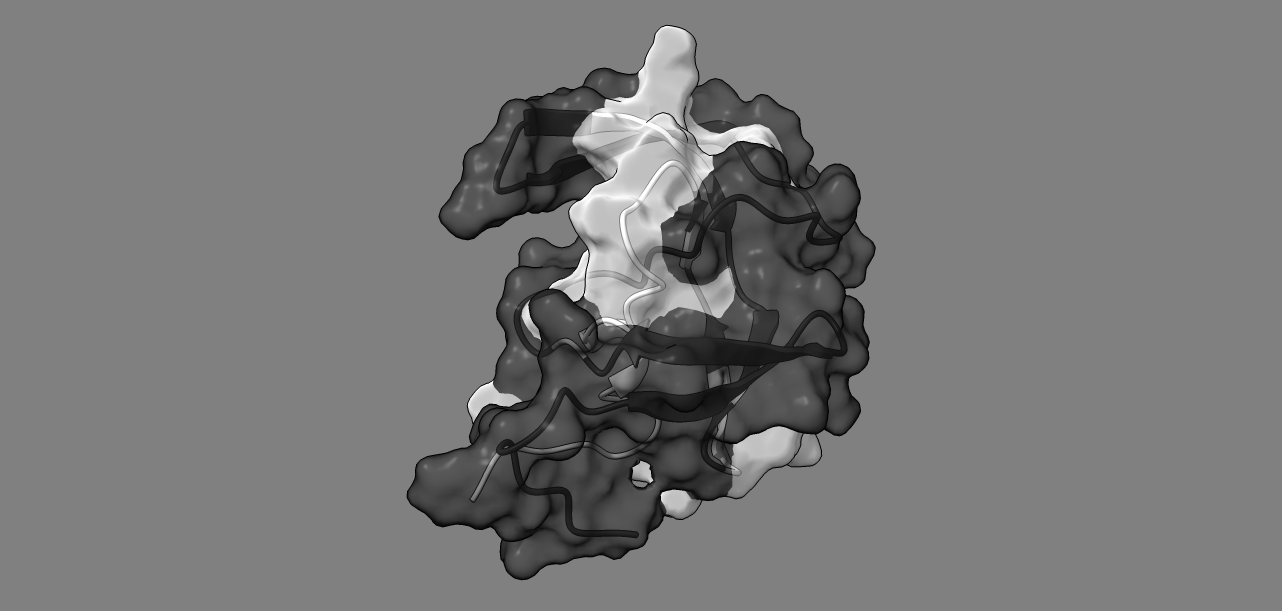

### image2.png

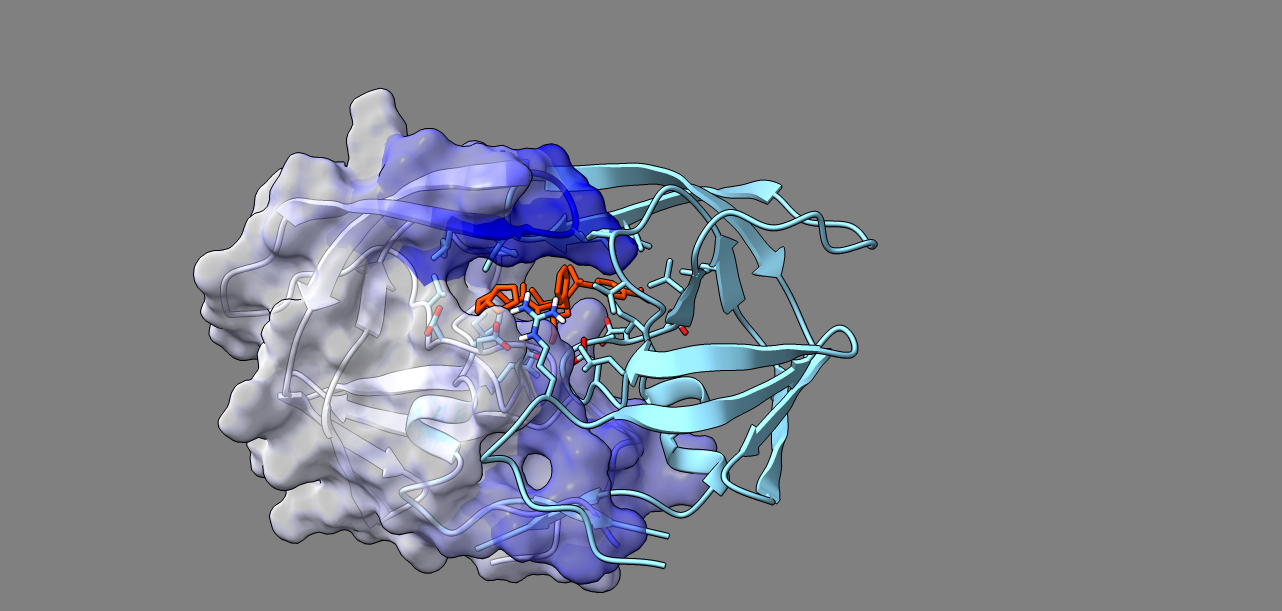

### image2.png

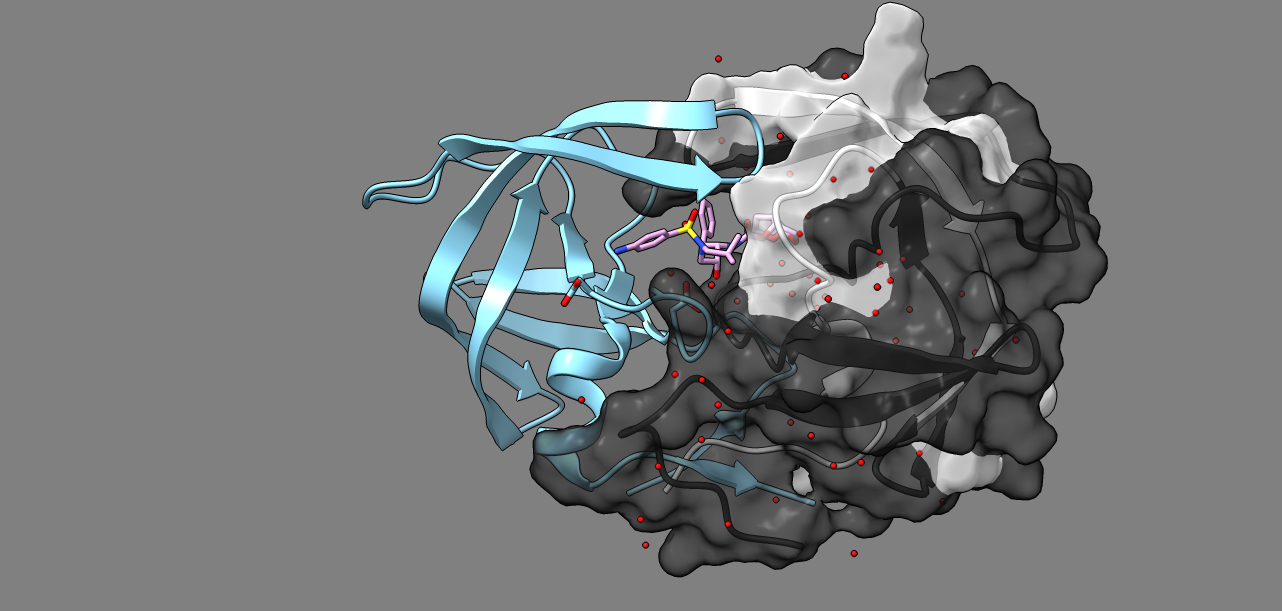

### image2.png

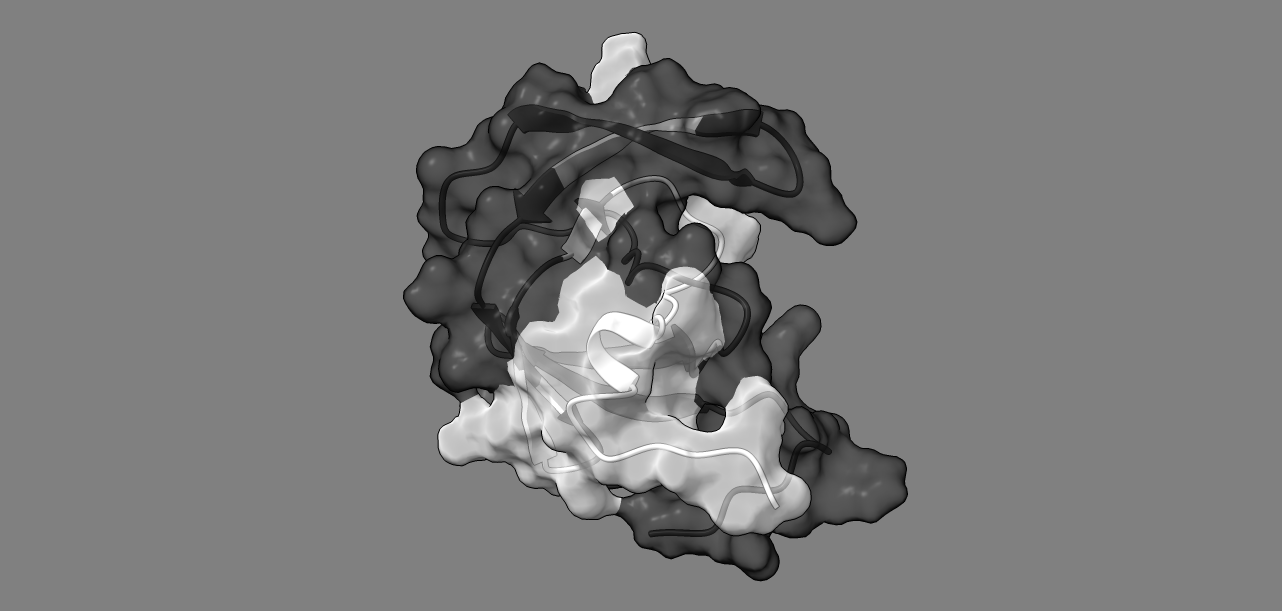

### image3.png

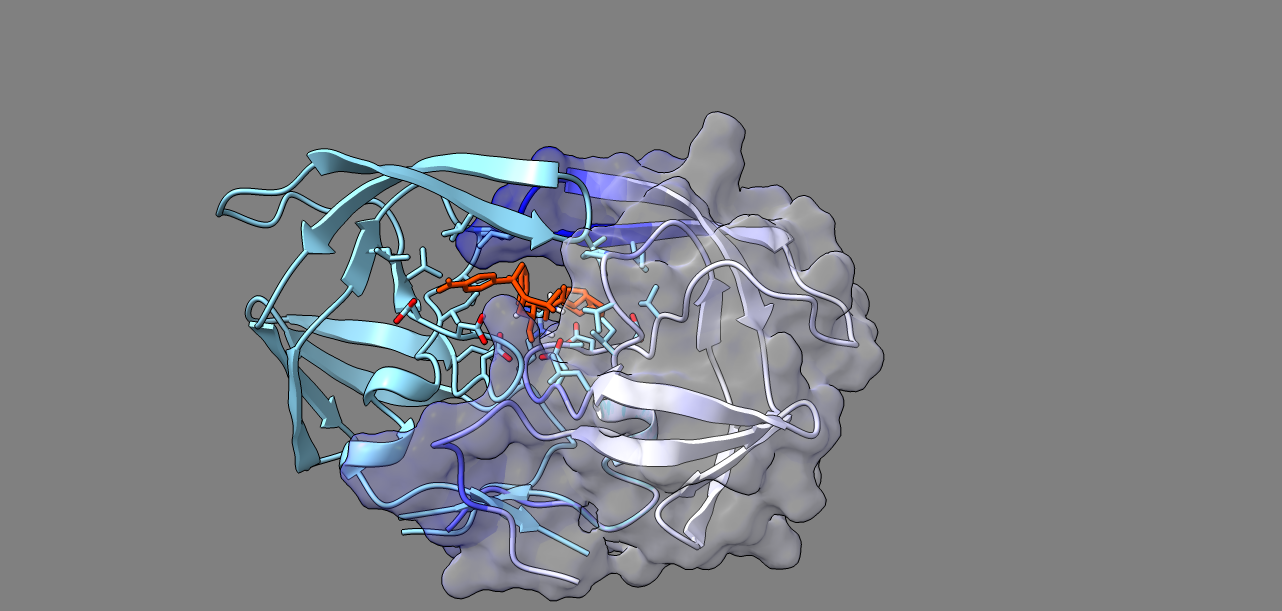

### image3.png

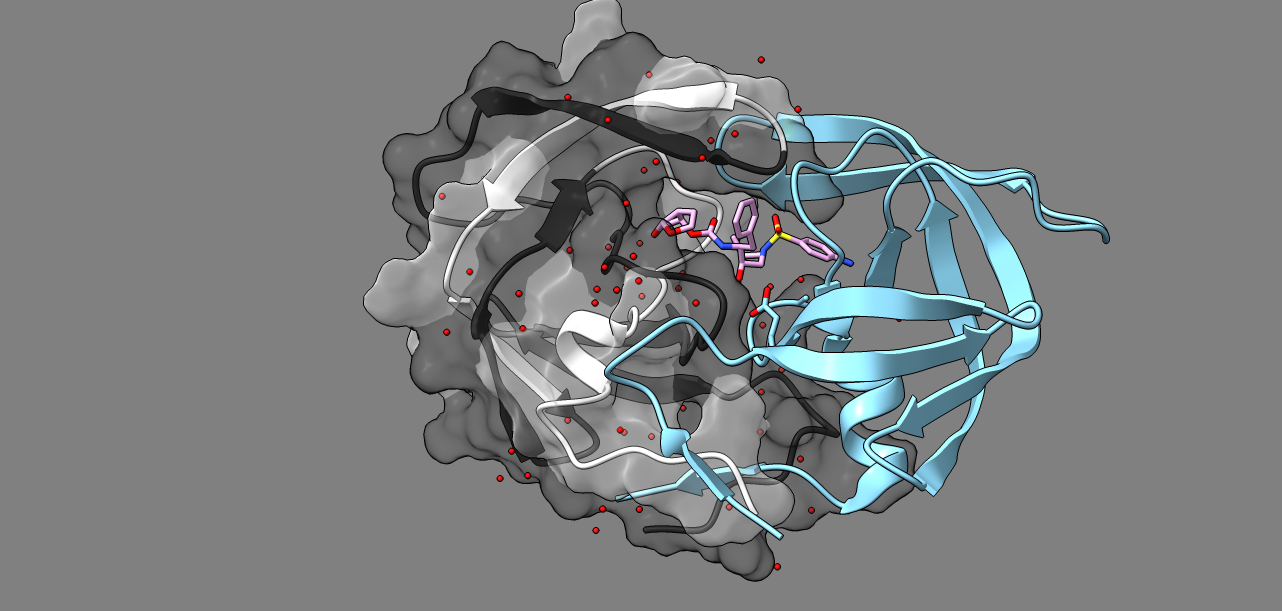

### image3.png

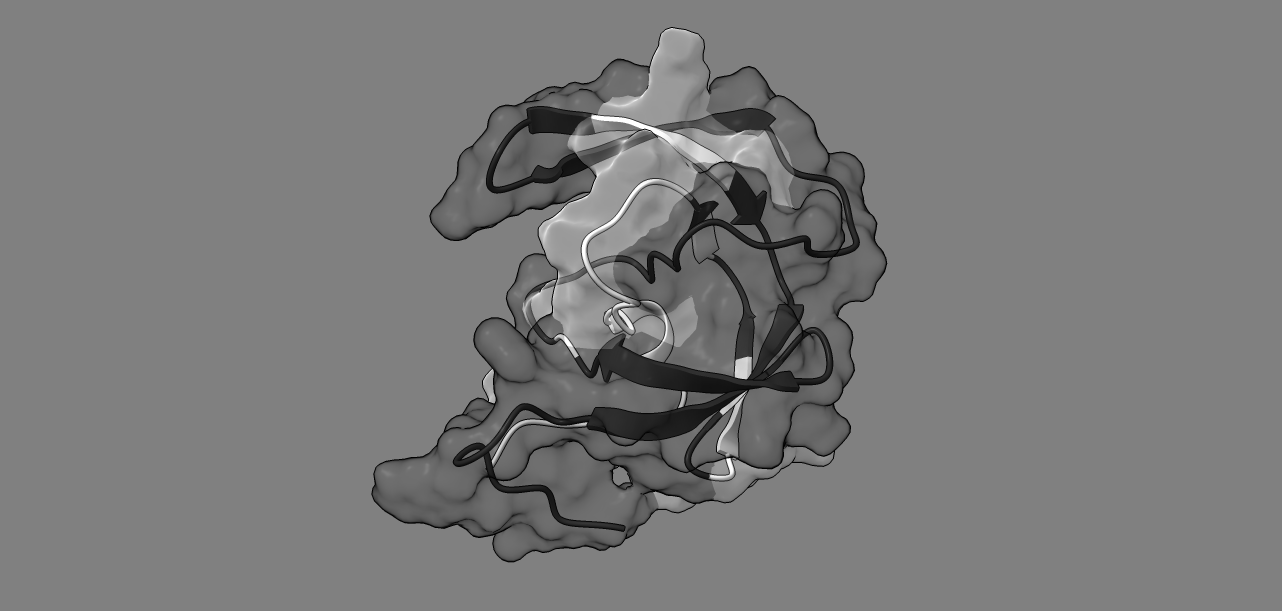

### image4.png

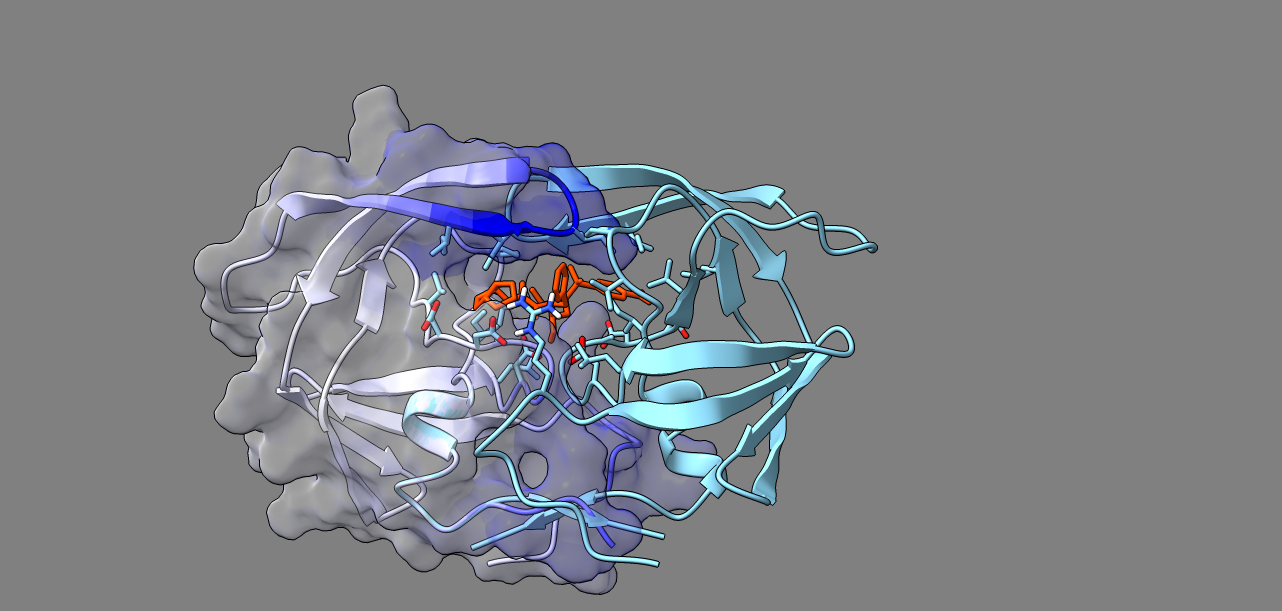

### image4.png

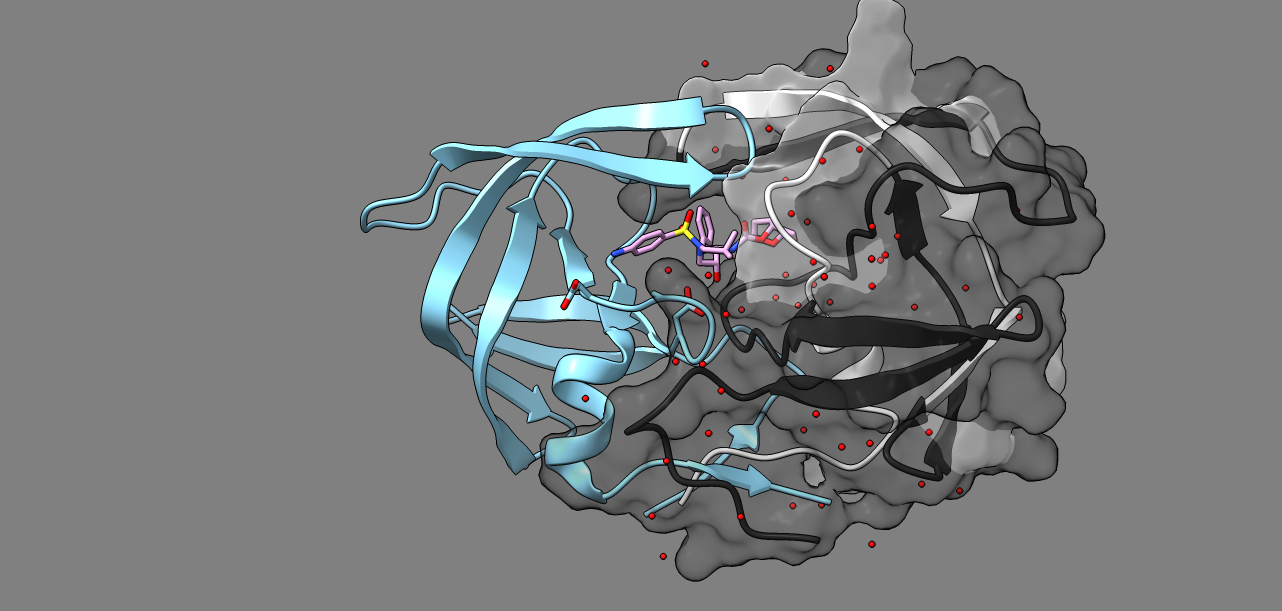

### image4.png

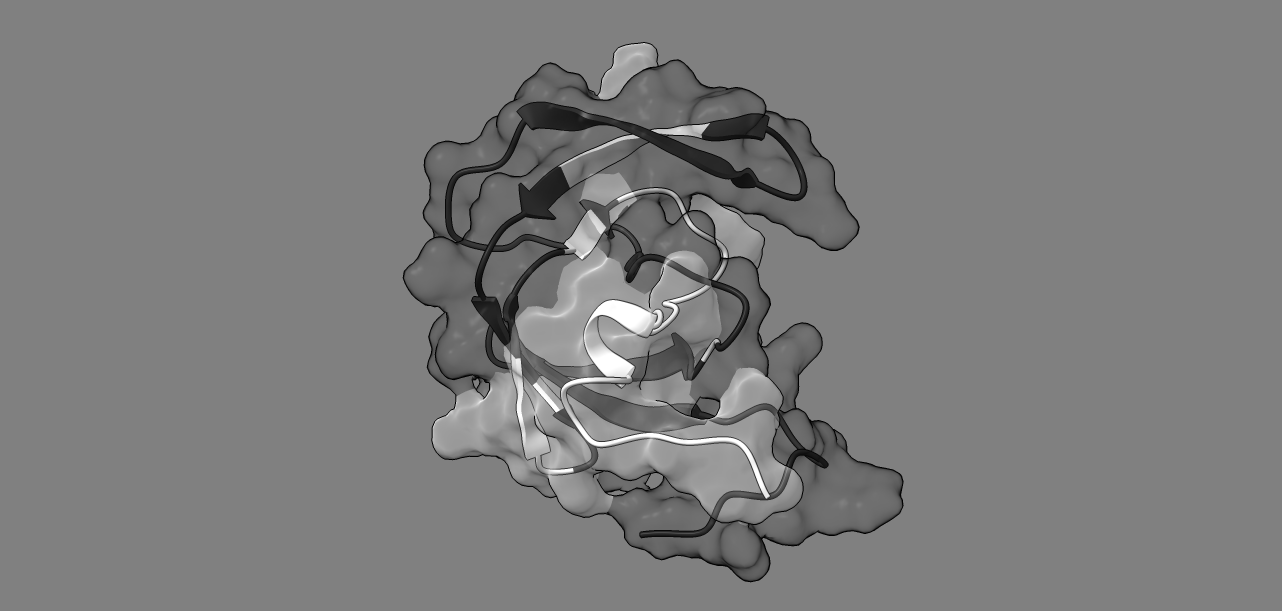

### image5.png

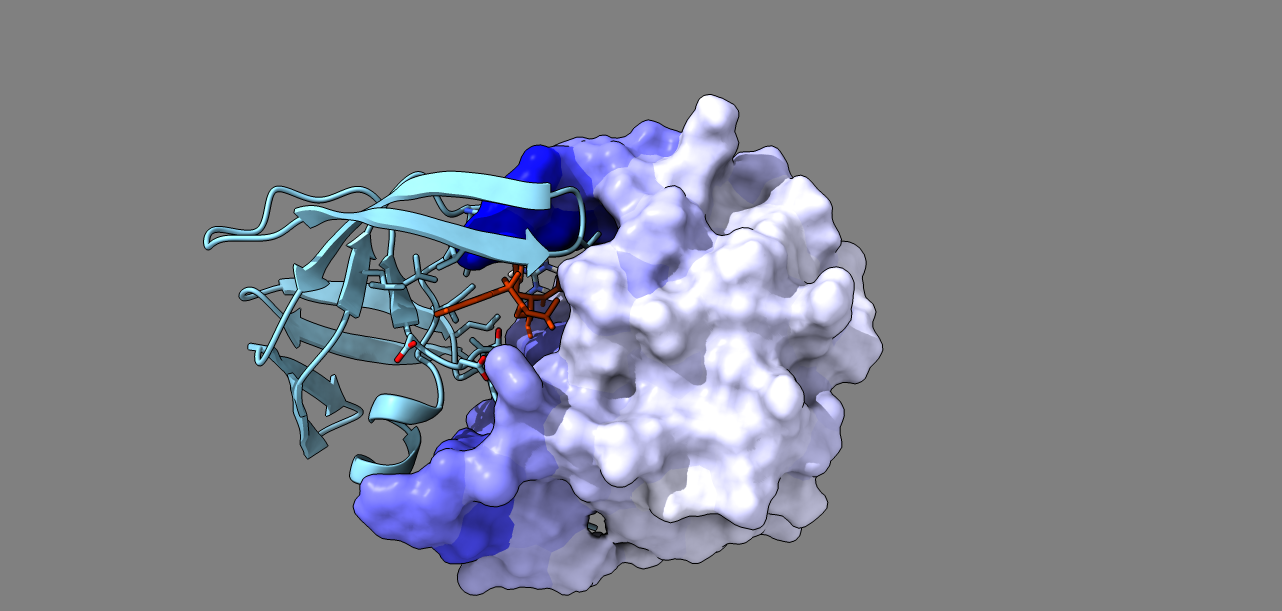

### image5.png

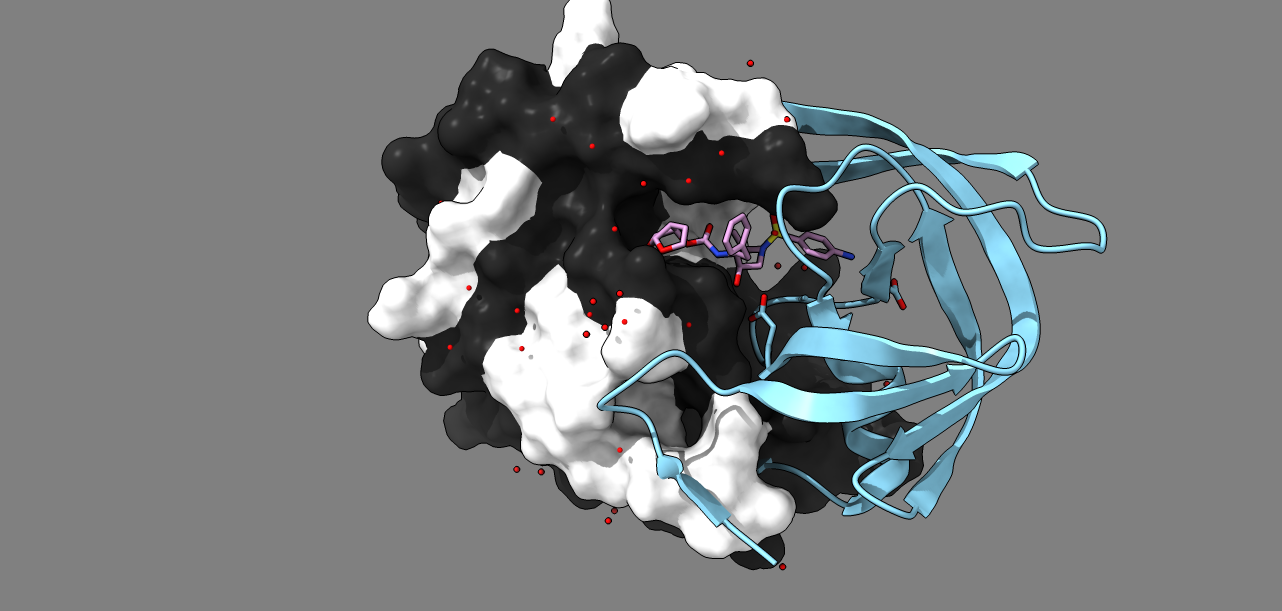

### image5.png

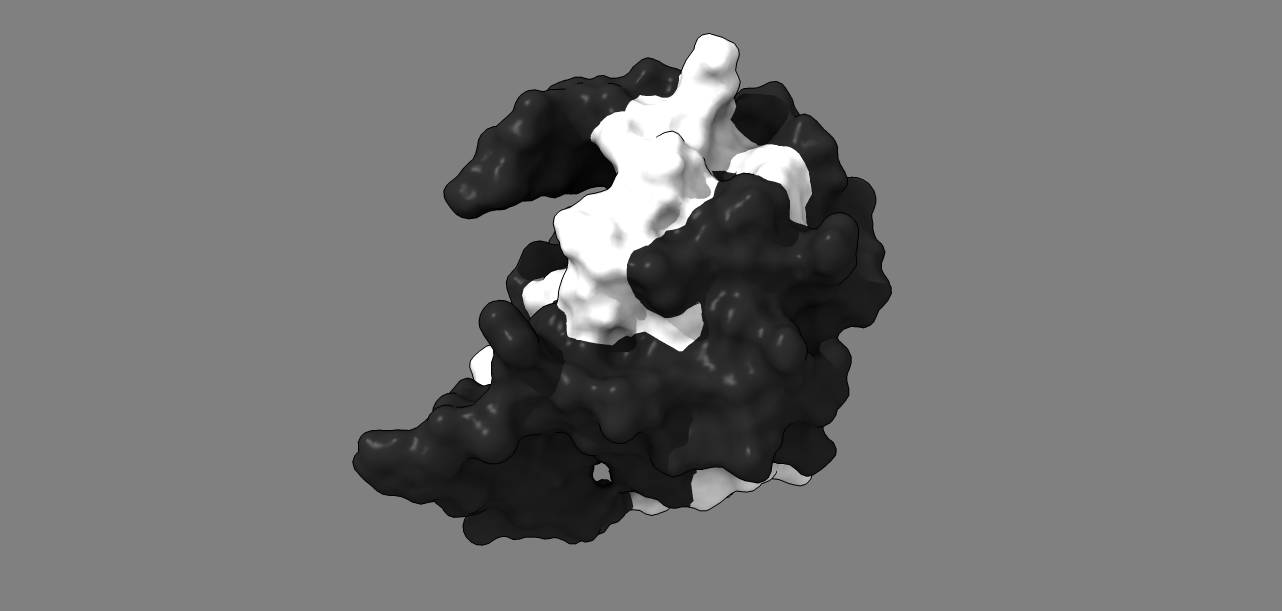

### image6.png

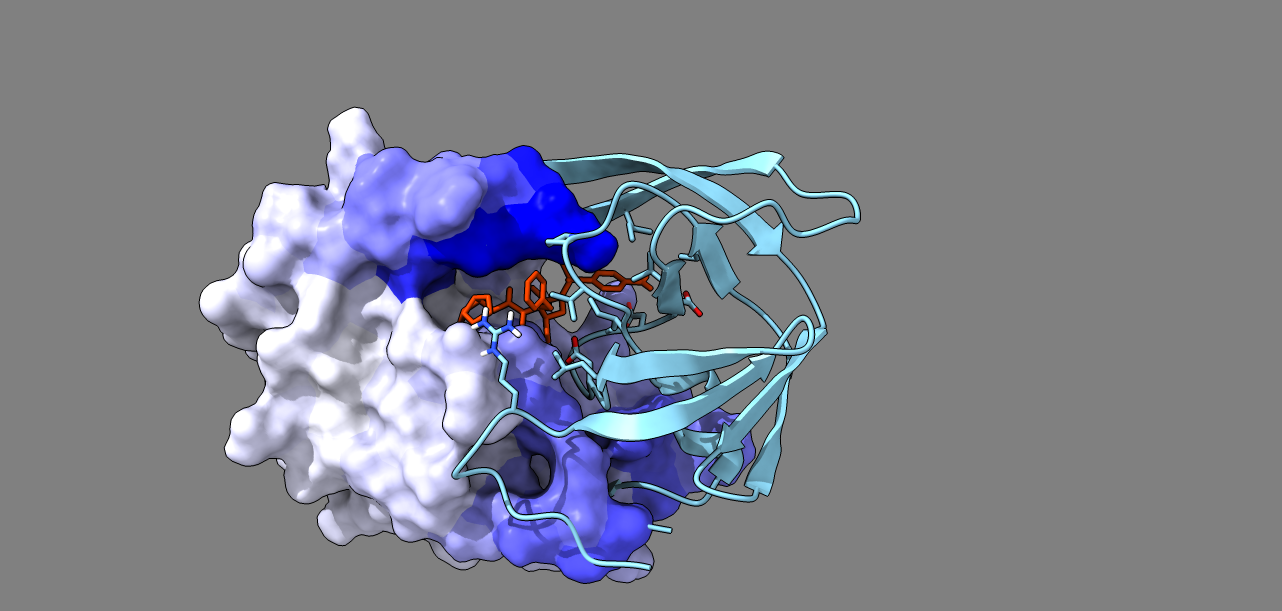

### image6.png

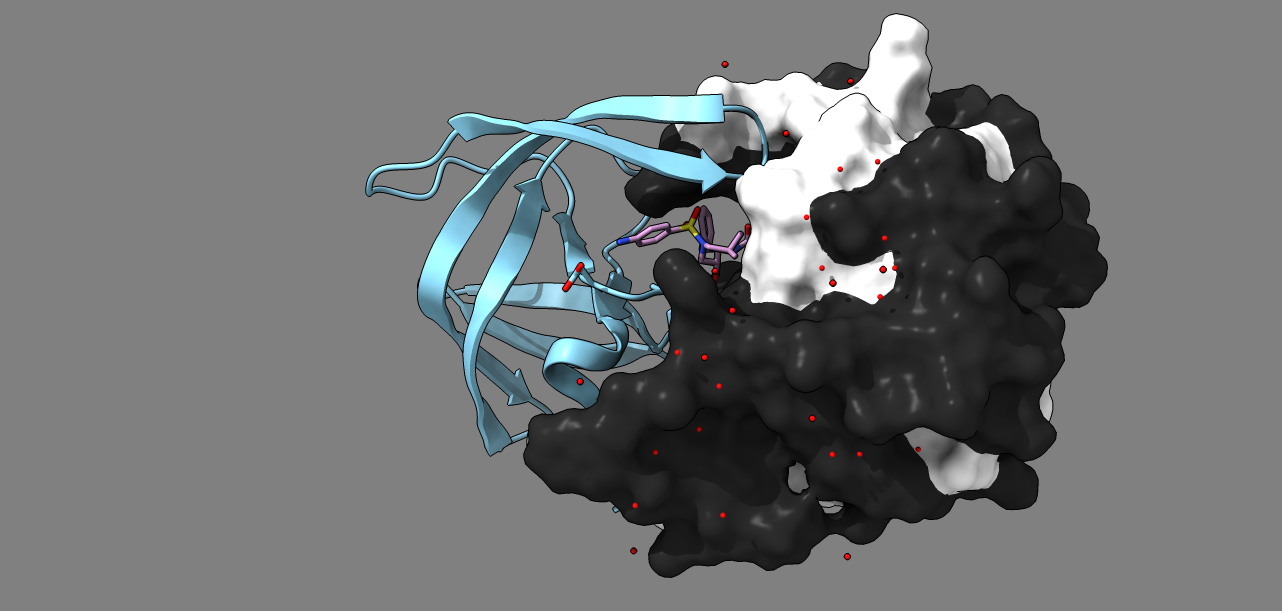

### image6.png

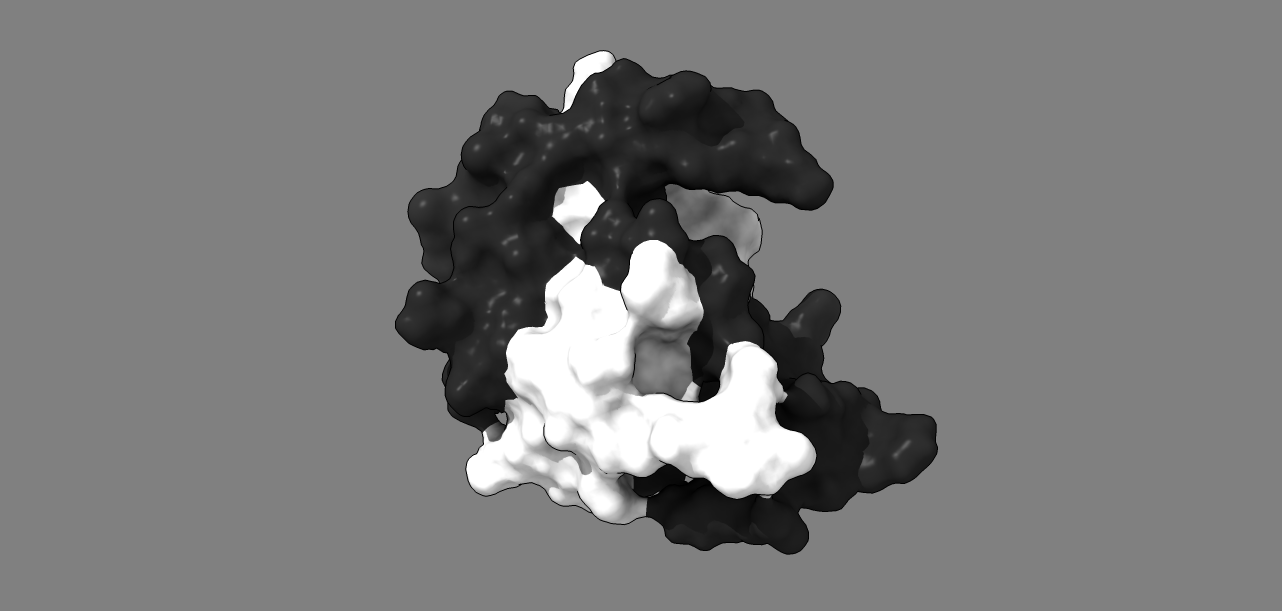

### image7.png

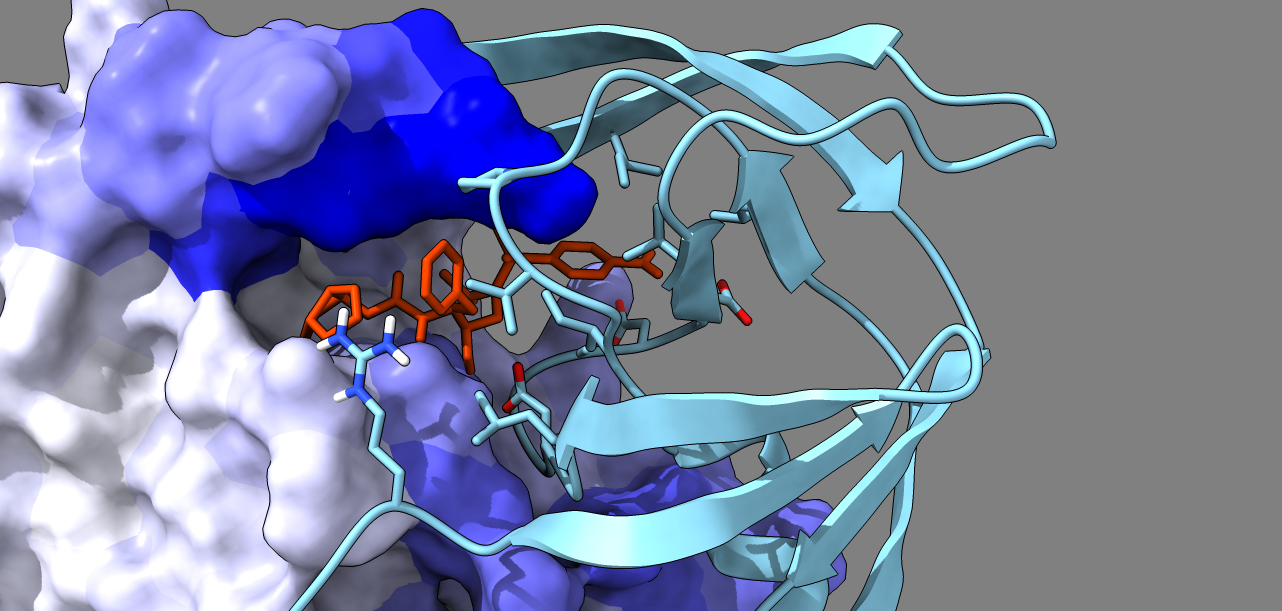

### image7.png

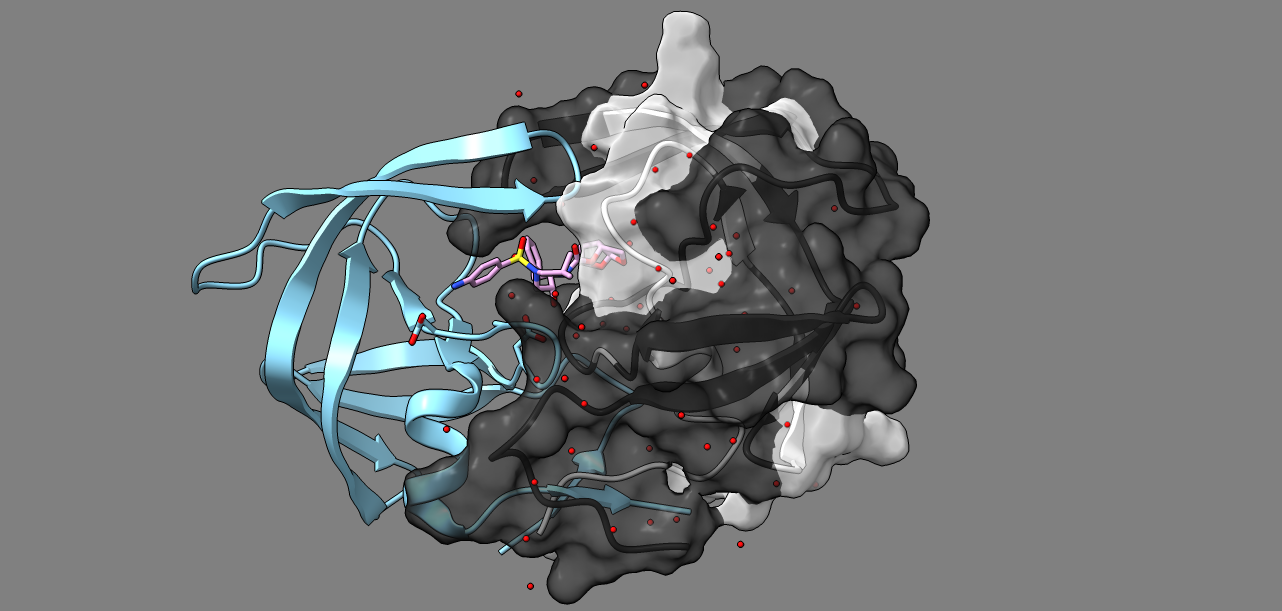

### image8.png

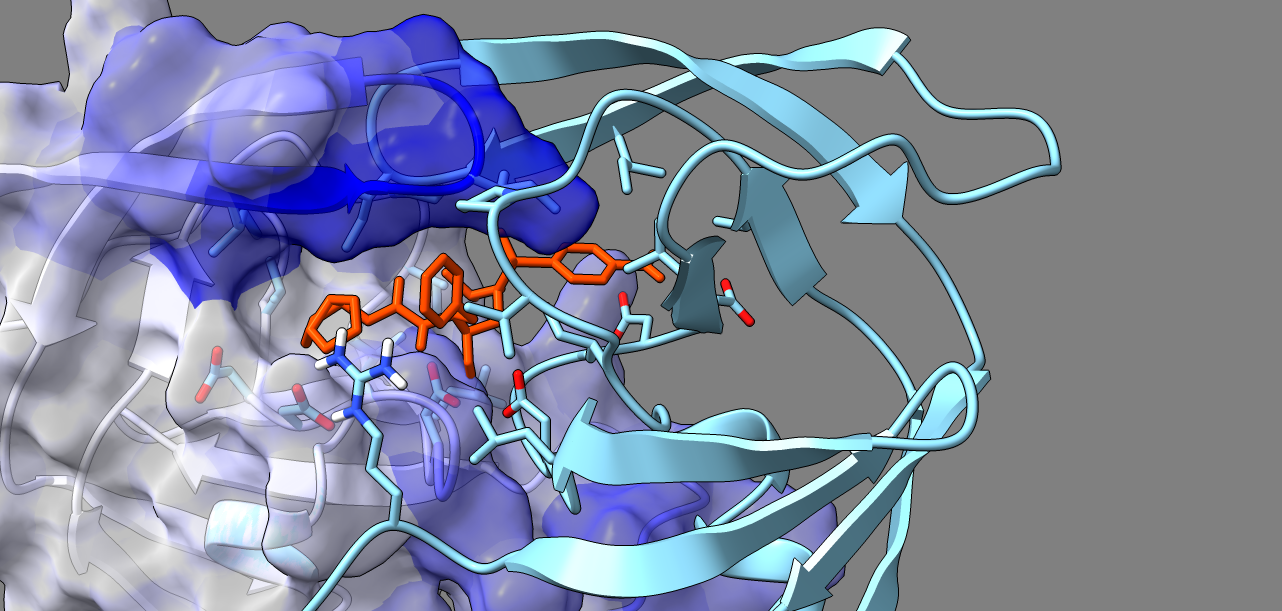

### image8.png

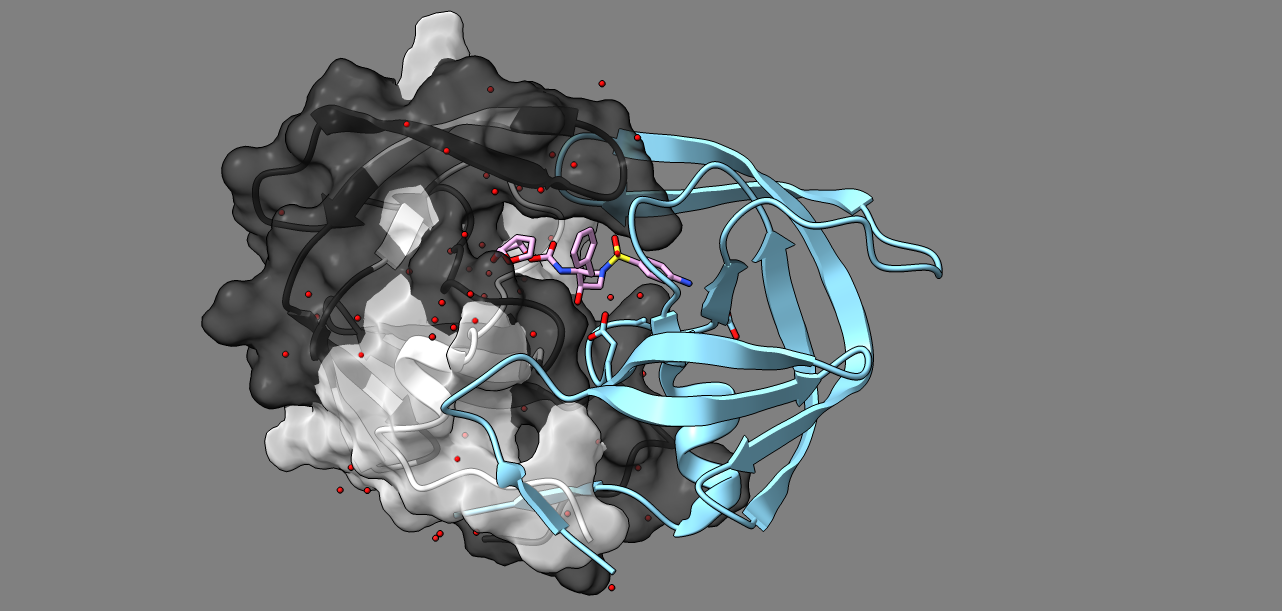

### image9.png

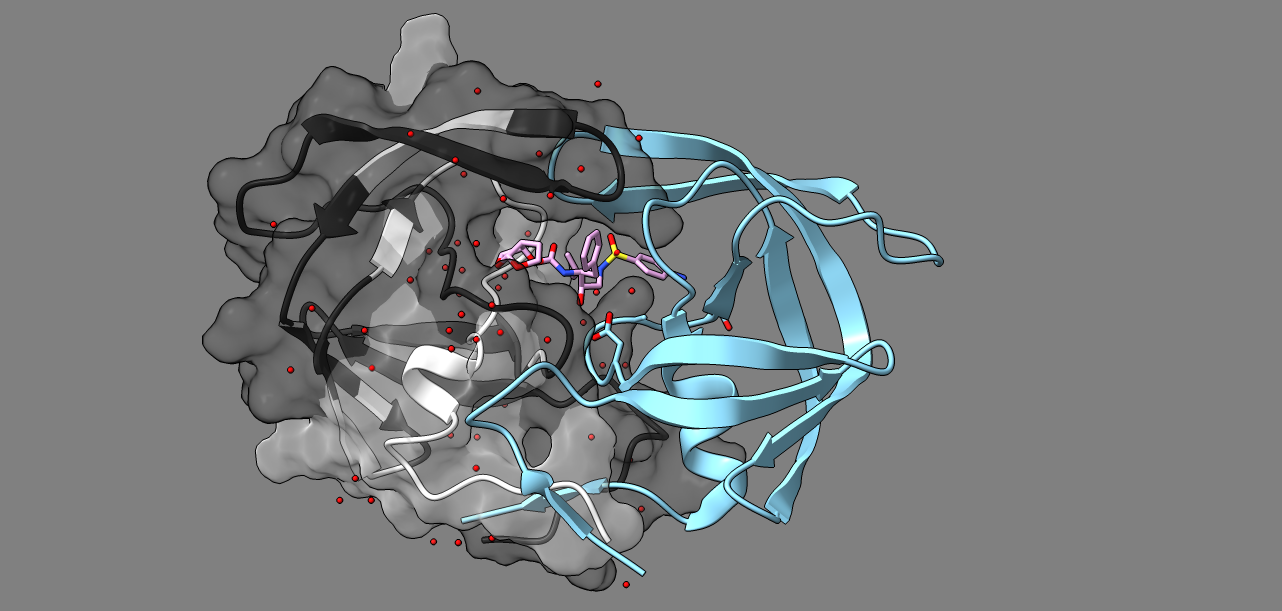

### image10.png

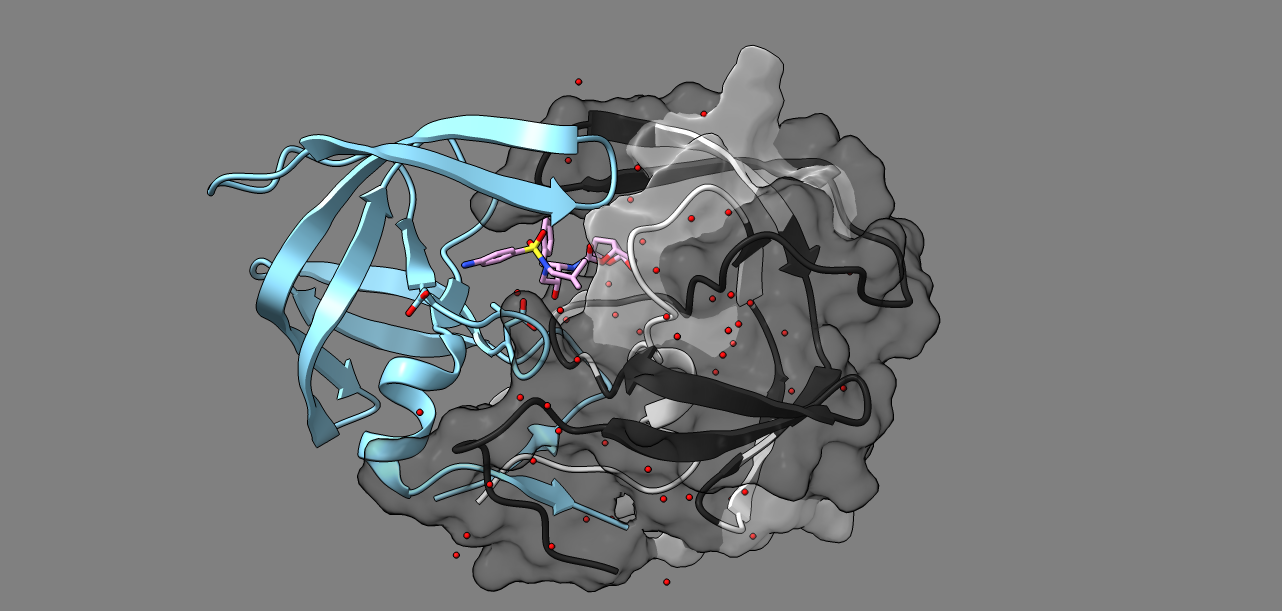

### image11.png

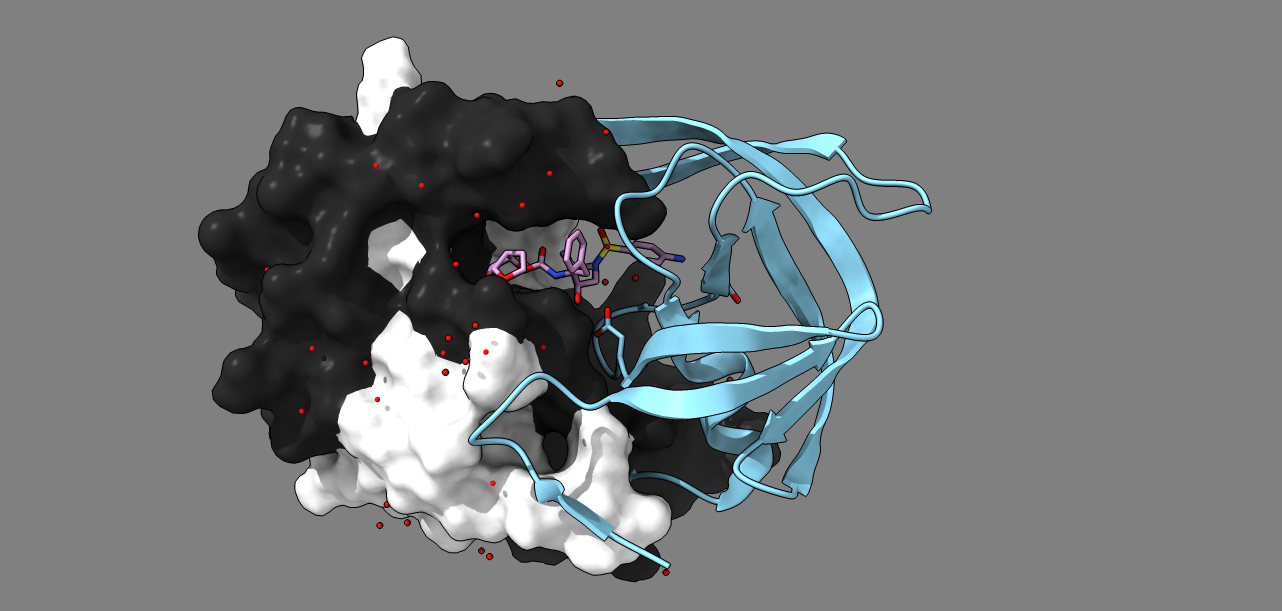

### image12.png

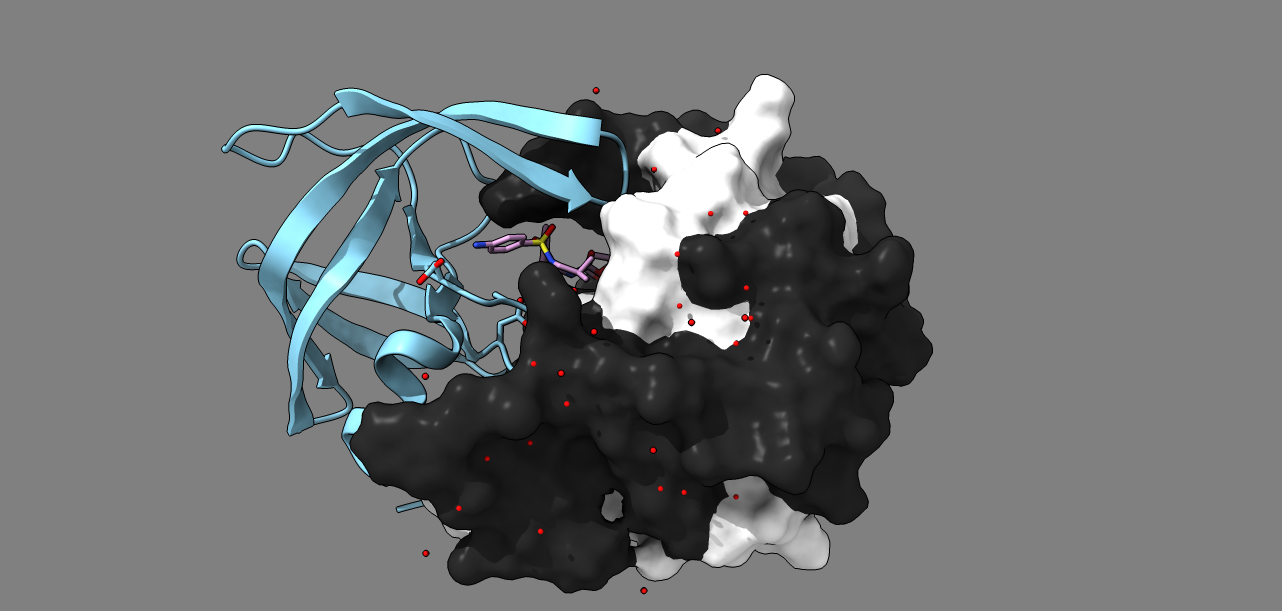

### learningPLOTfunctional.pdf

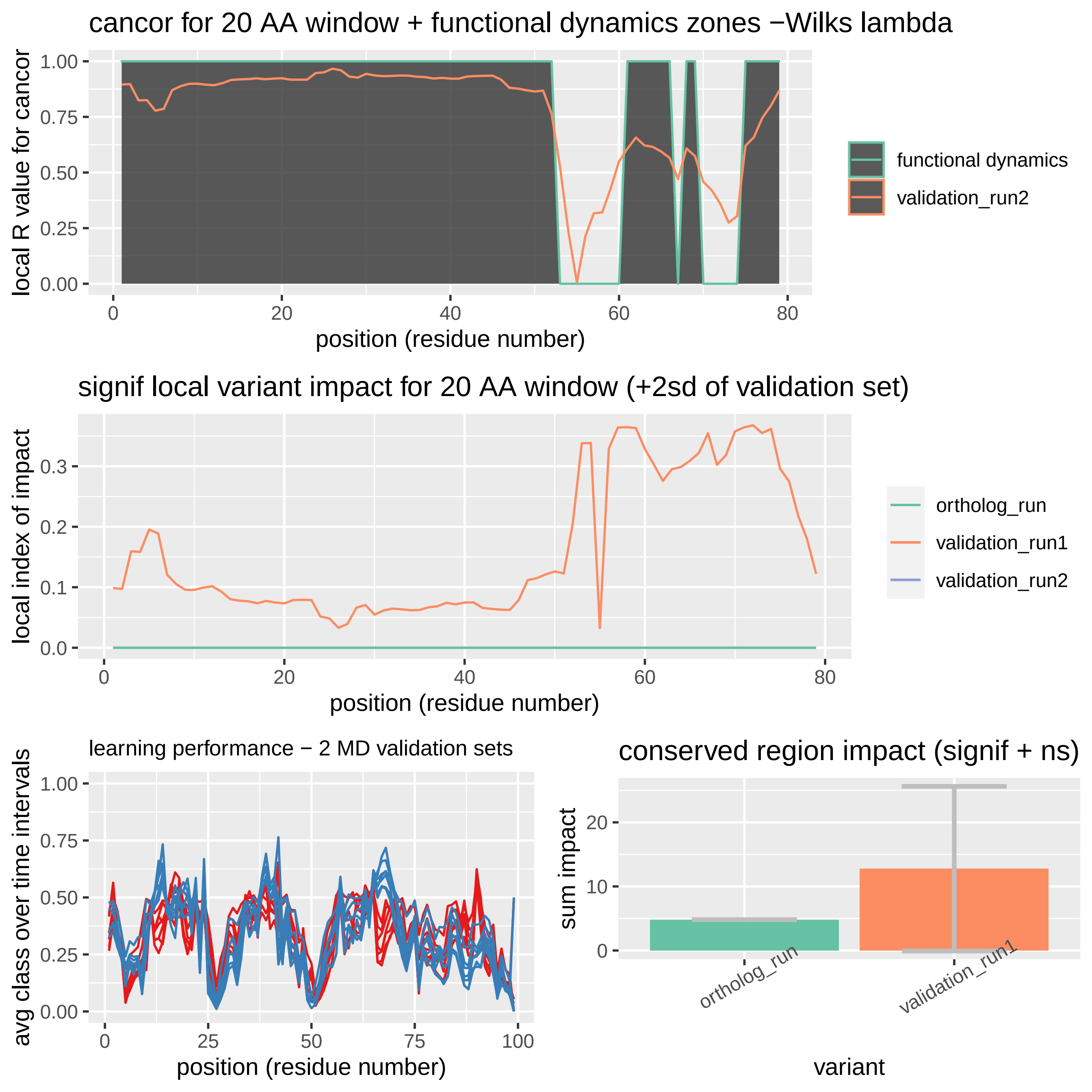

### learningPLOTfunctional.pdf

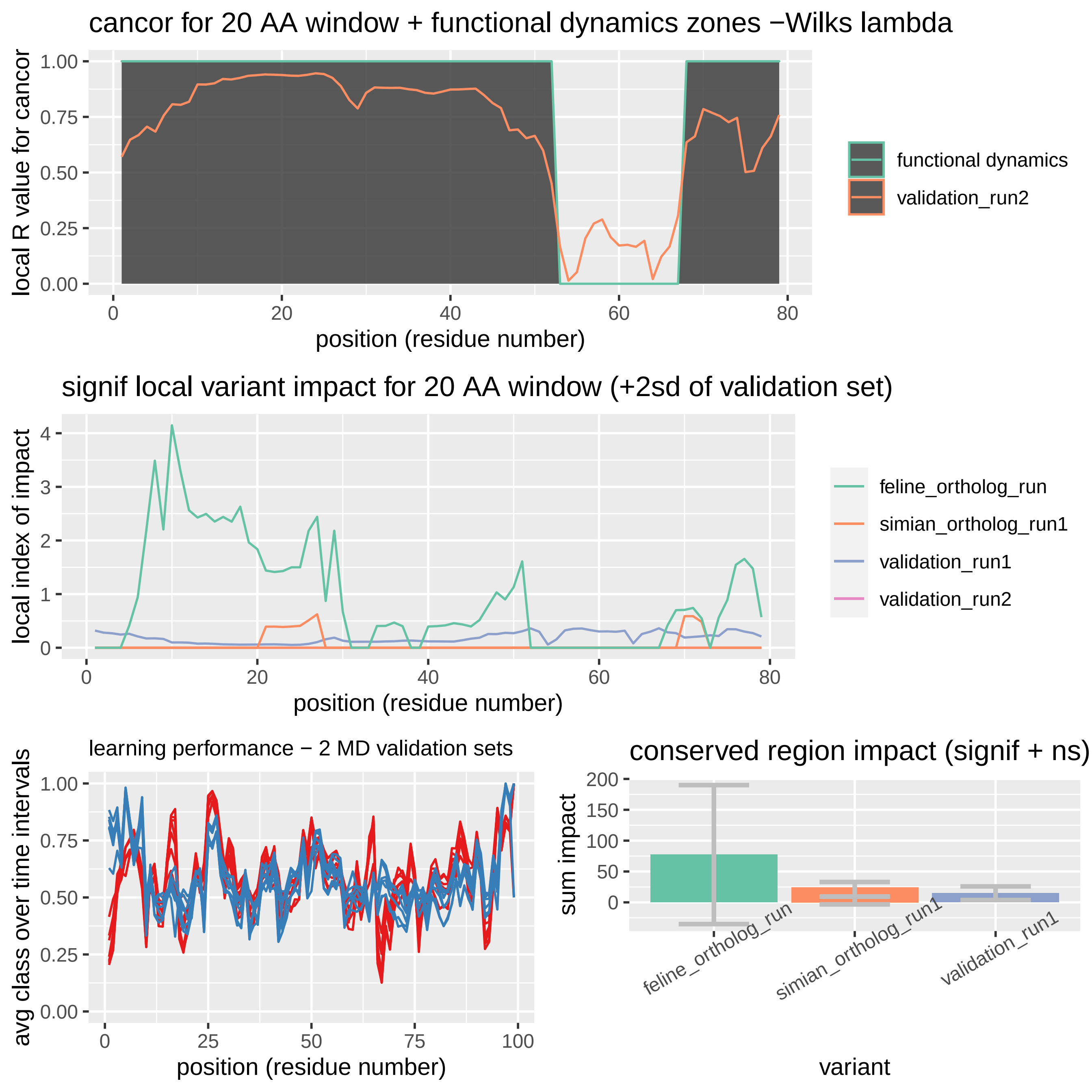

### learningPLOTortholog.pdf

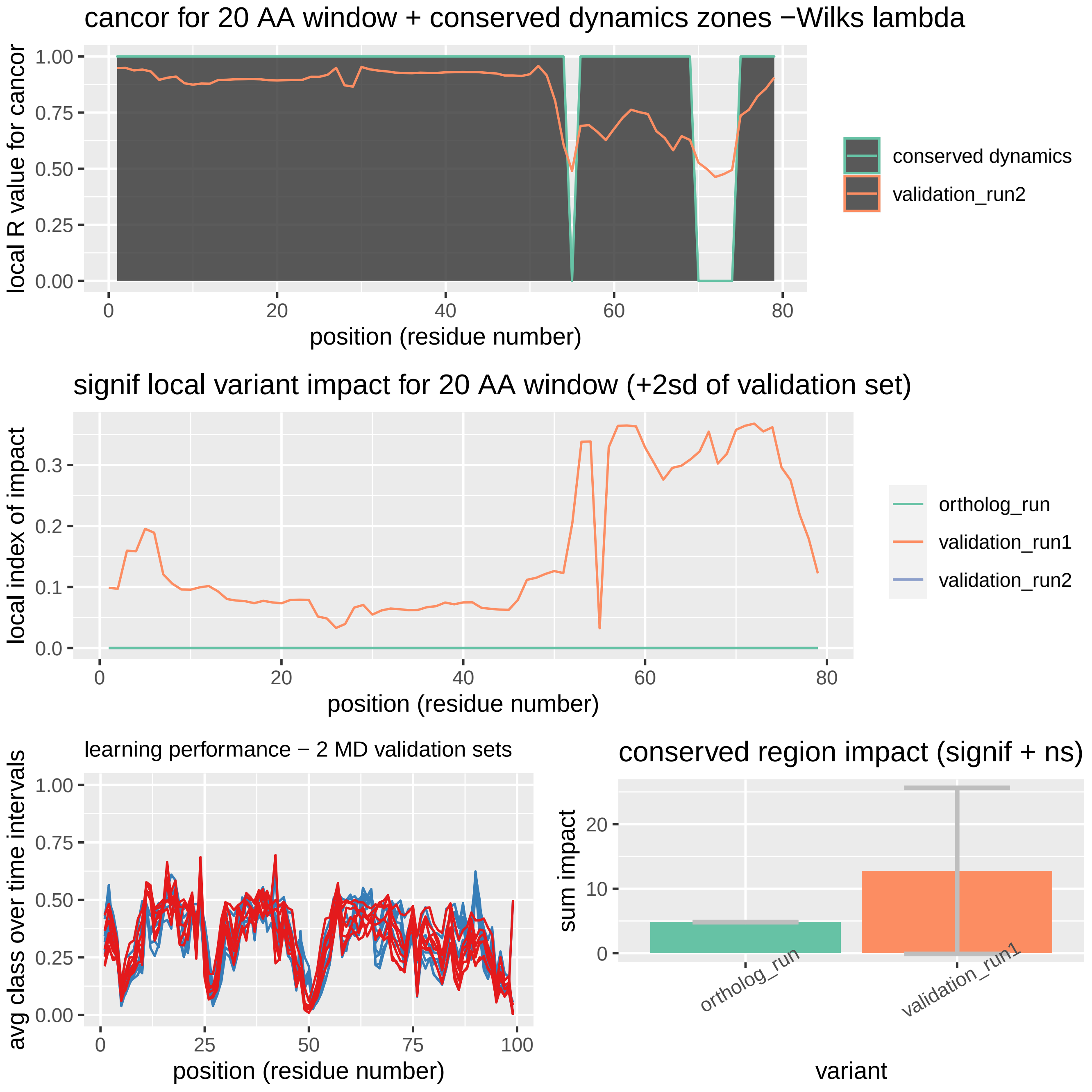

### learningPLOTortholog.pdf

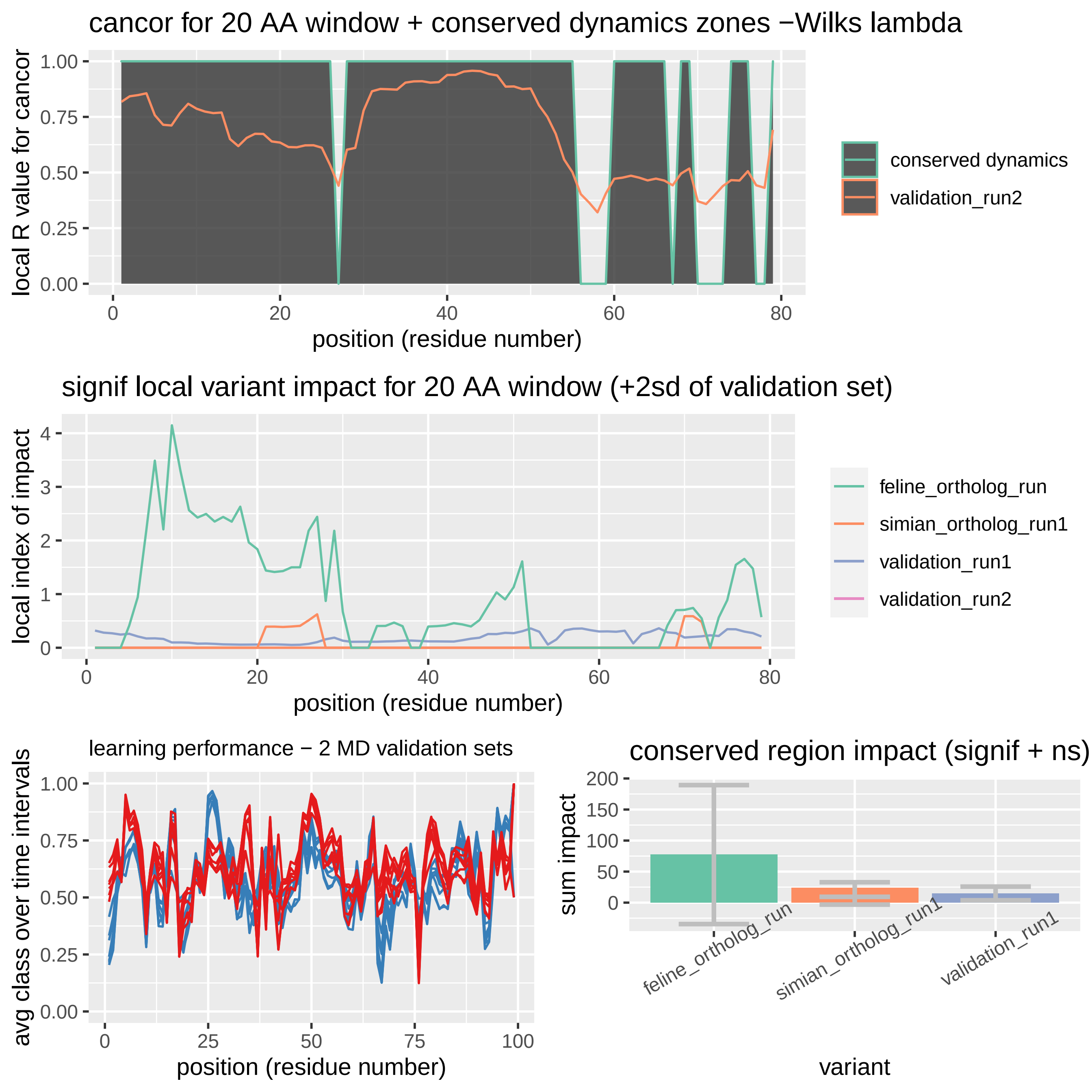

### MImatrix_LDA_1b11_monomer_feline.pdf

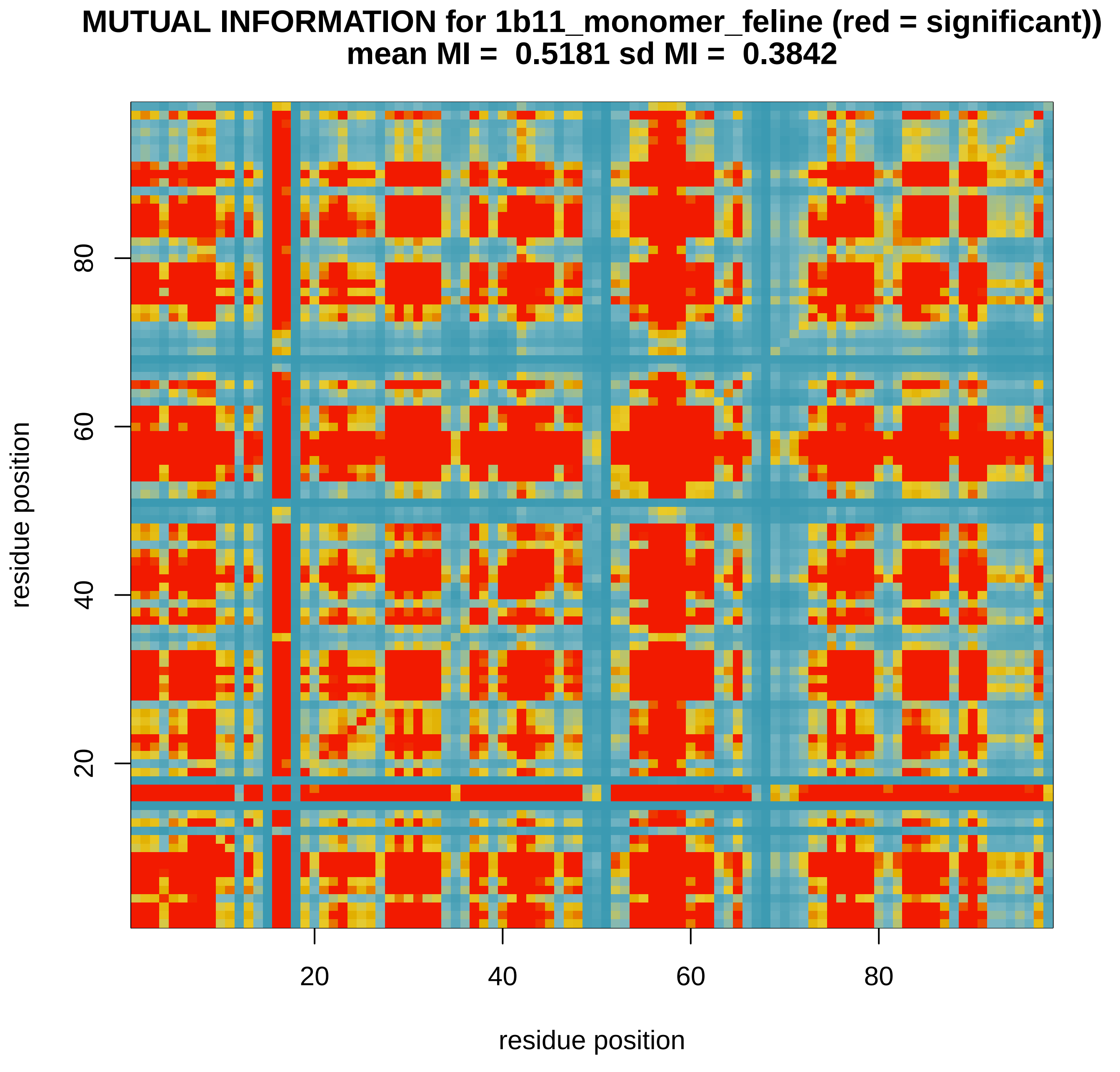

### MImatrix_LDA_1yti_monomer_simian.pdf

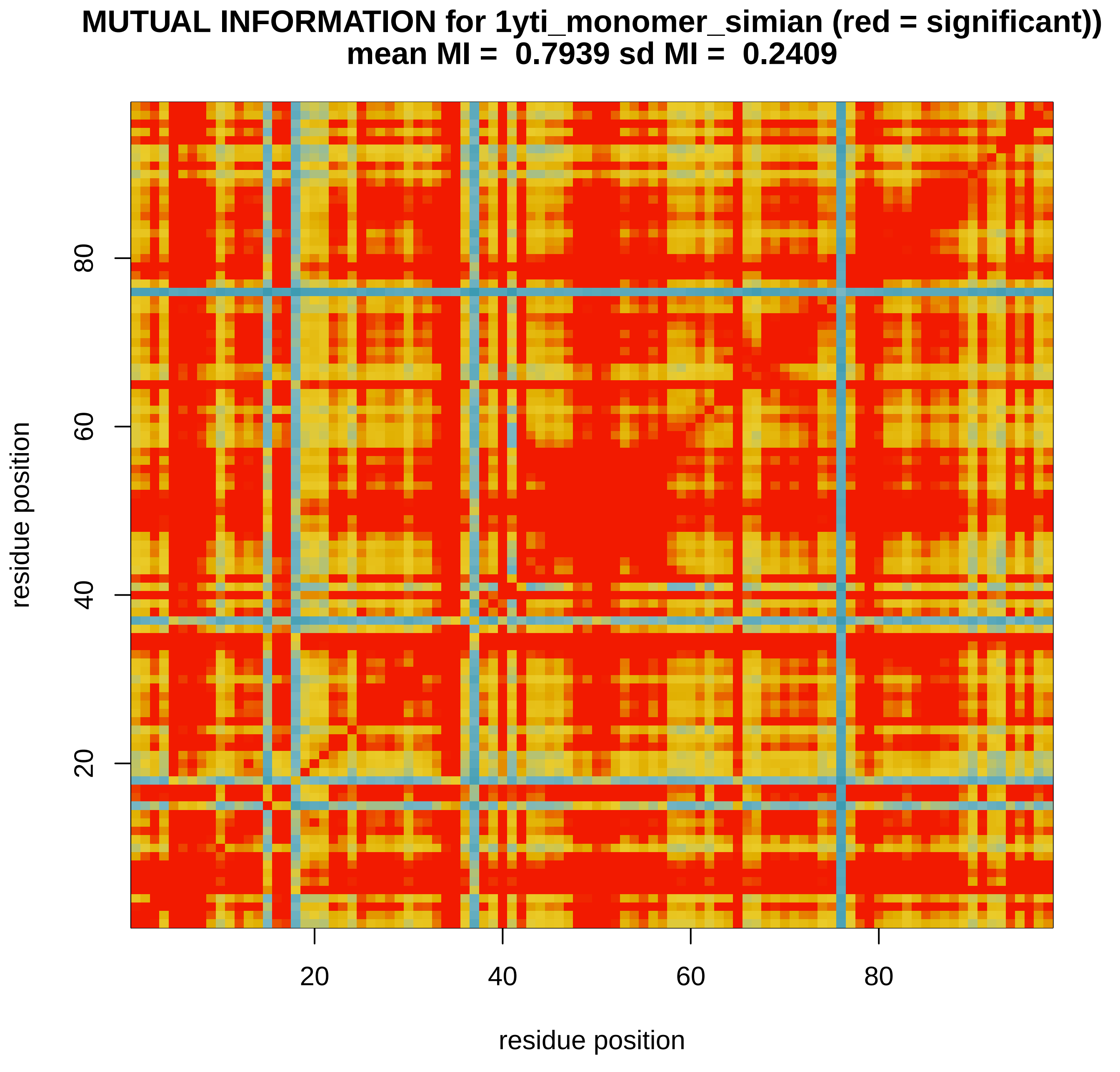

### MImatrix_LDA_1ytj_monomer_simian.pdf

##### MUTUAL INFORMATION for 1ytj_monomer_simian (red = significant))
##### mean MI = 0.7882 sd MI = 0.2424

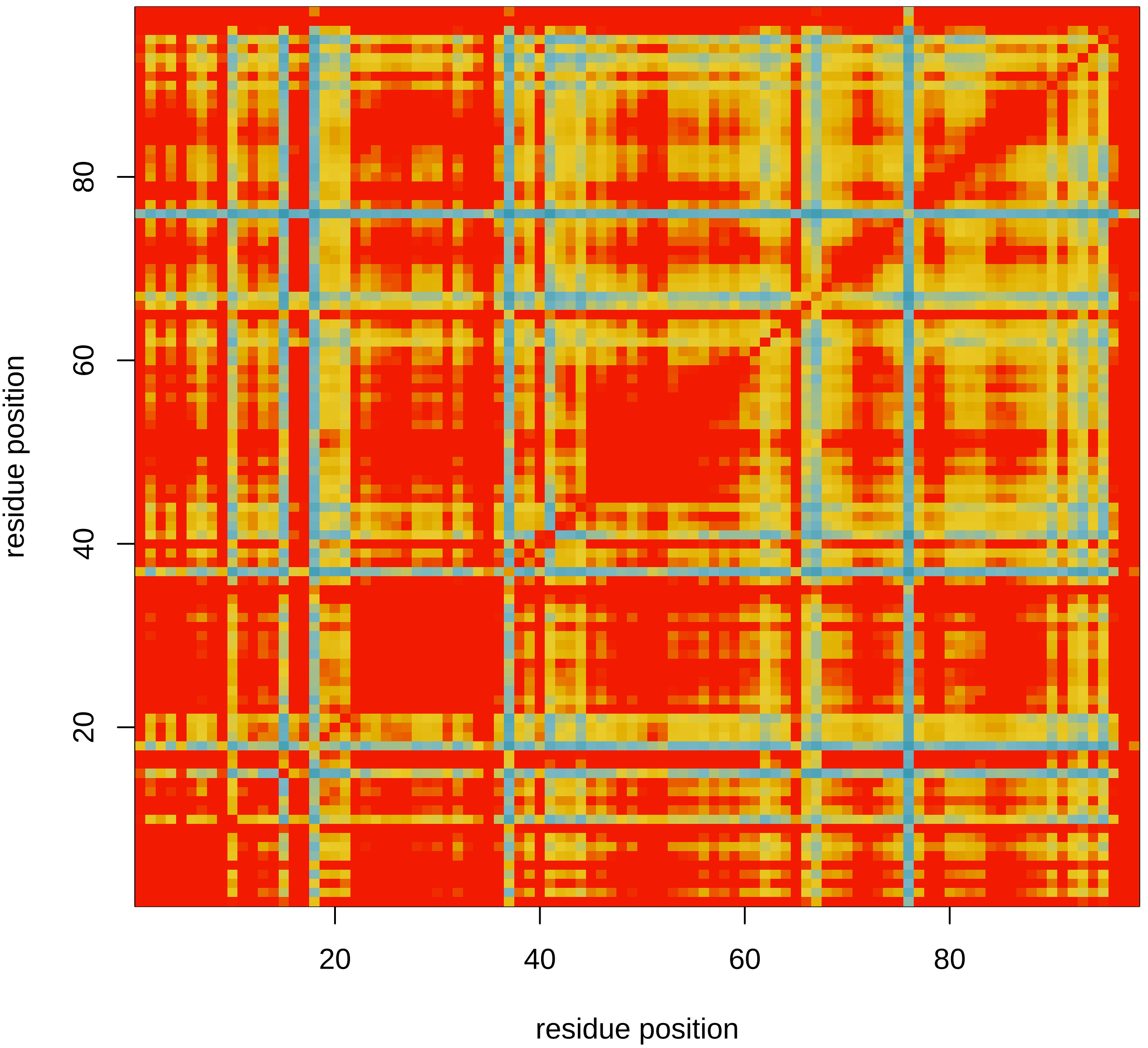

### MImatrix_LDA_6dgx_monomer_human_1.pdf

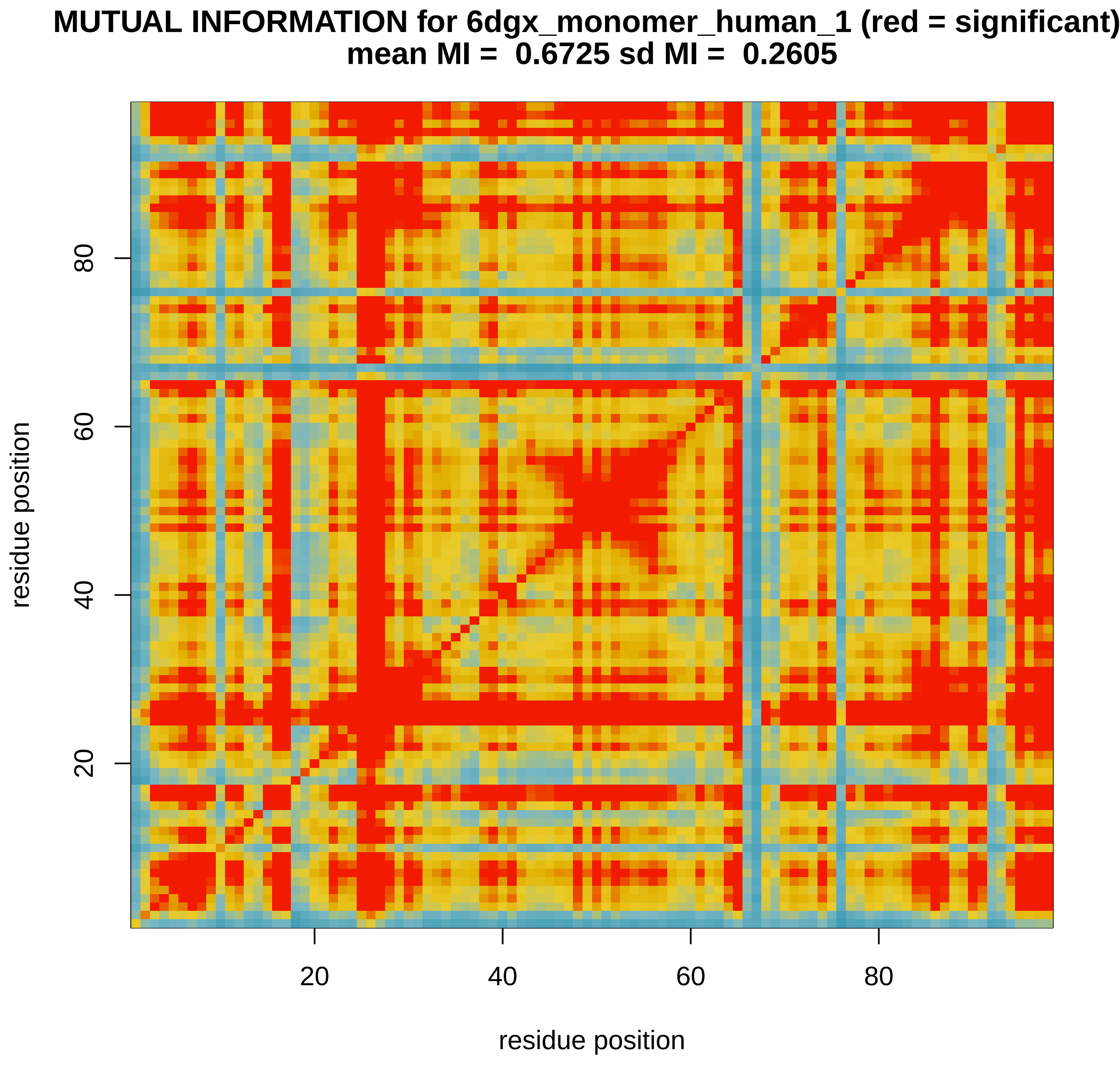

### MImatrix_LDA_6dgx_monomer_human_2.pdf

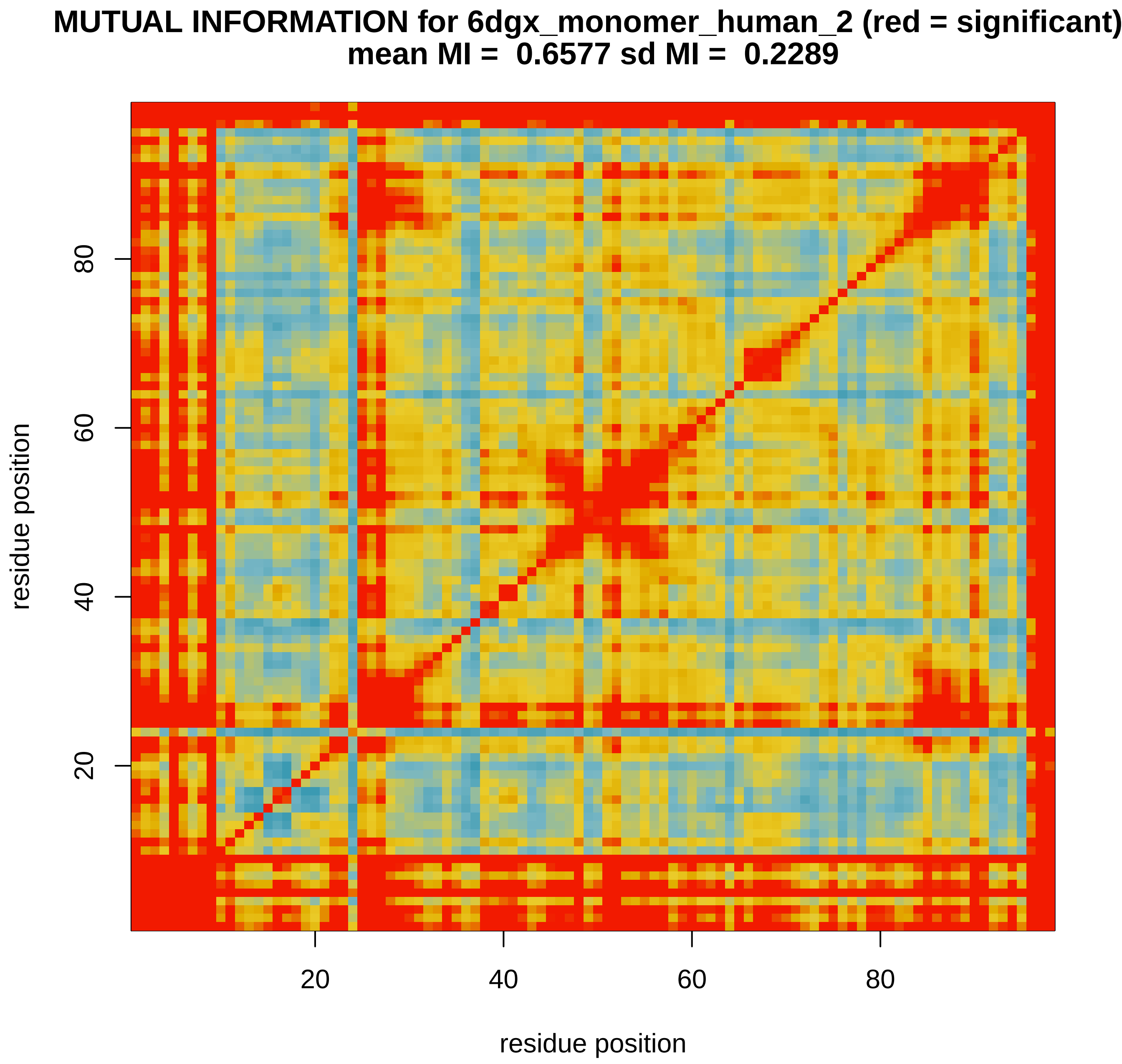

### MImatrix_LDA_6dgx_nopep_human.pdf

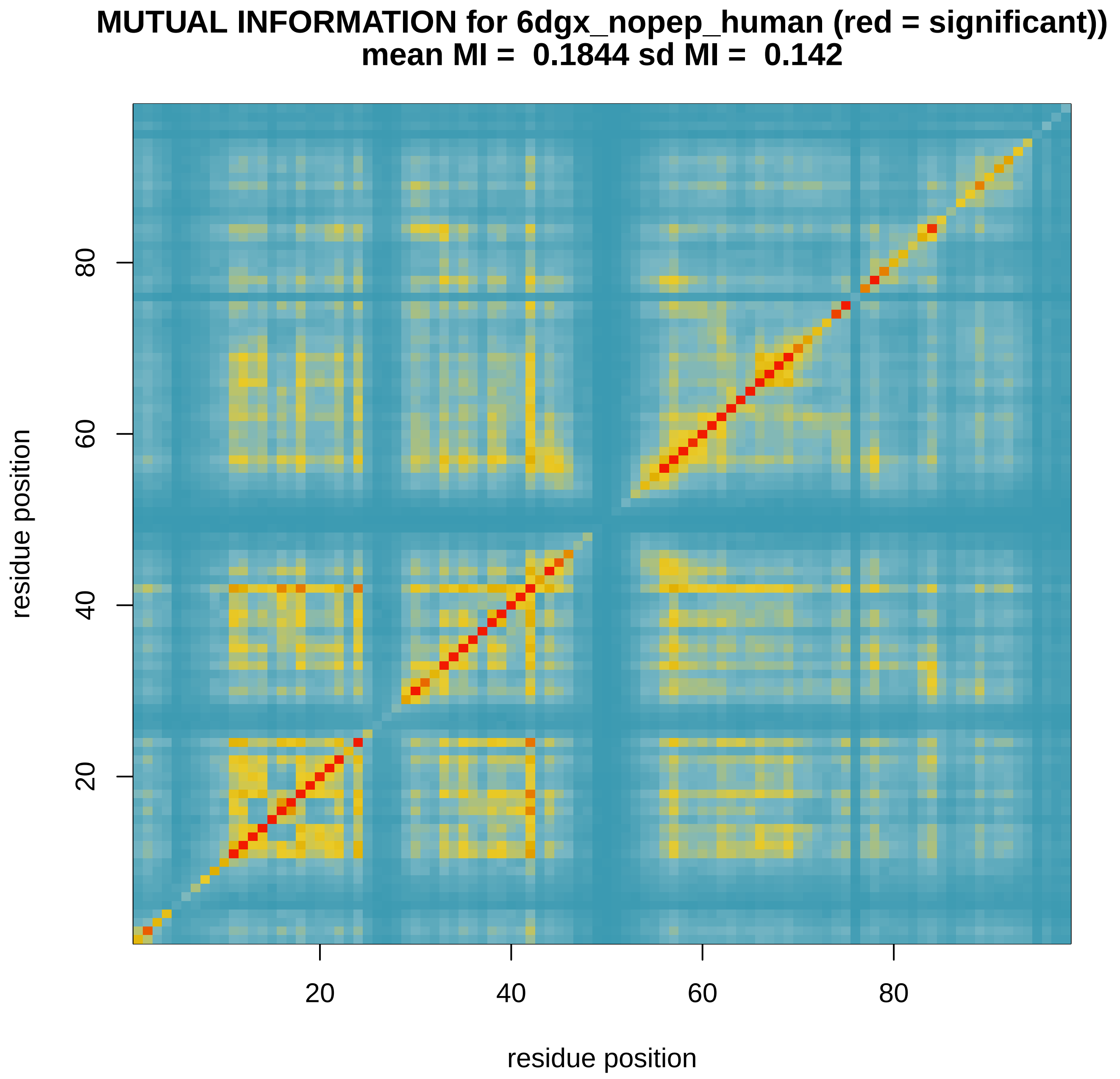

### MImatrix_LDA_6dgx_nopep_human_1.pdf

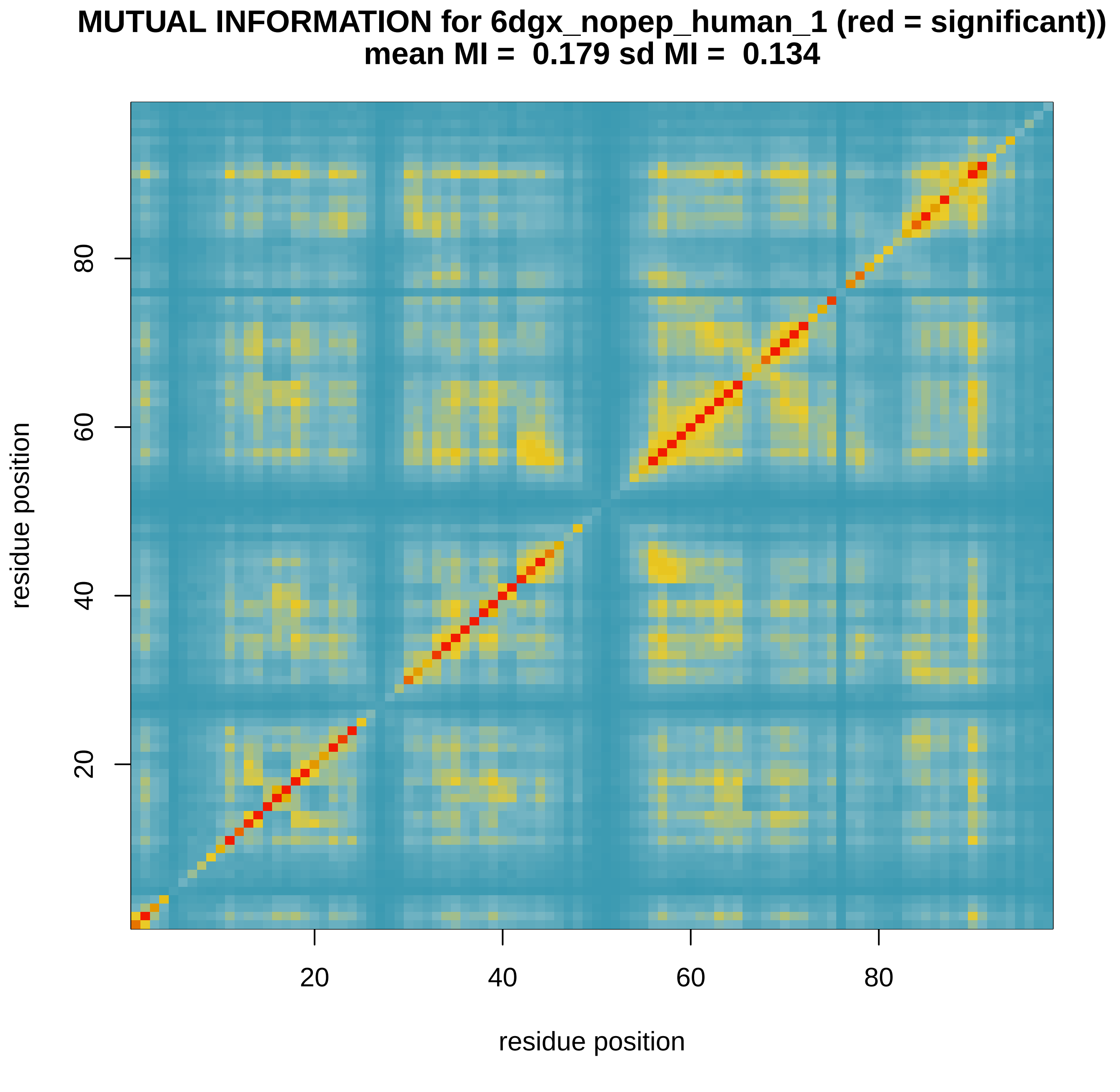

### MImatrix_LDA_6dgx_nopep_human_2.pdf

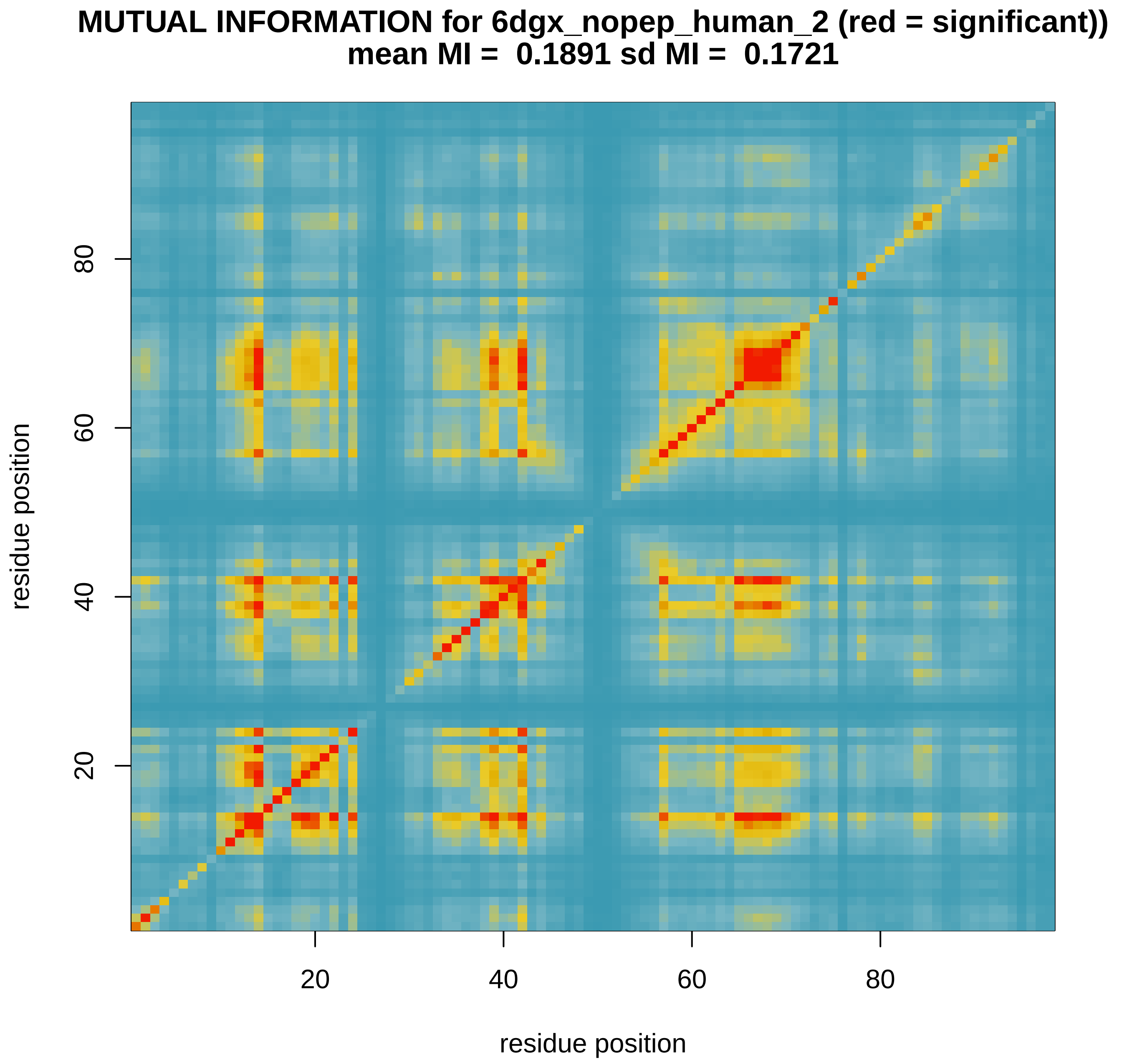

### MImatrix_RFOR_1b11_monomer_feline.pdf

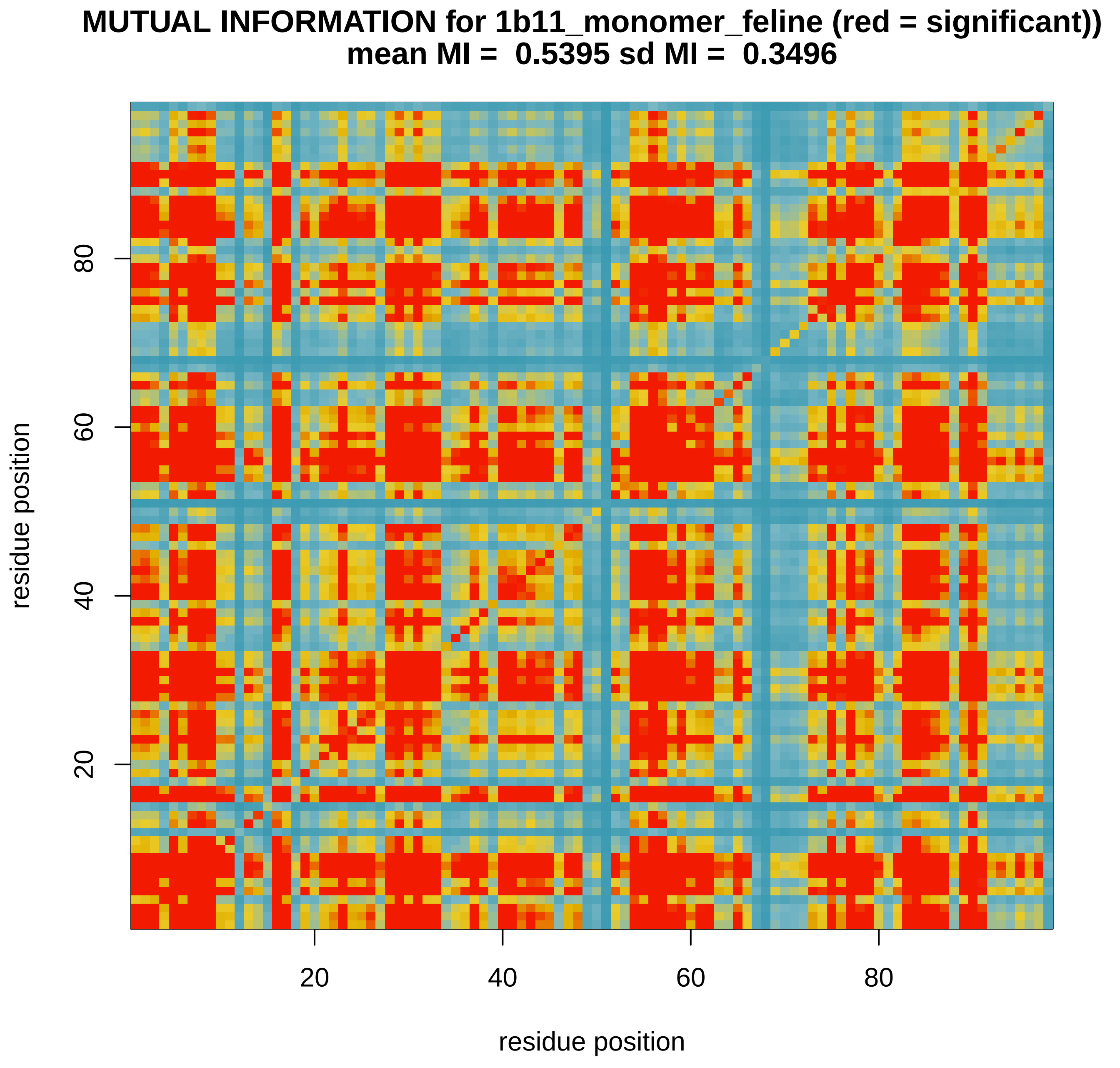

### MImatrix_RFOR_1yti_monomer_simian.pdf

**MUTUAL INFORMATION for 1yti_monomer_simian (red = significant))**
**mean MI =  0.7999 sd MI =  0.2192**

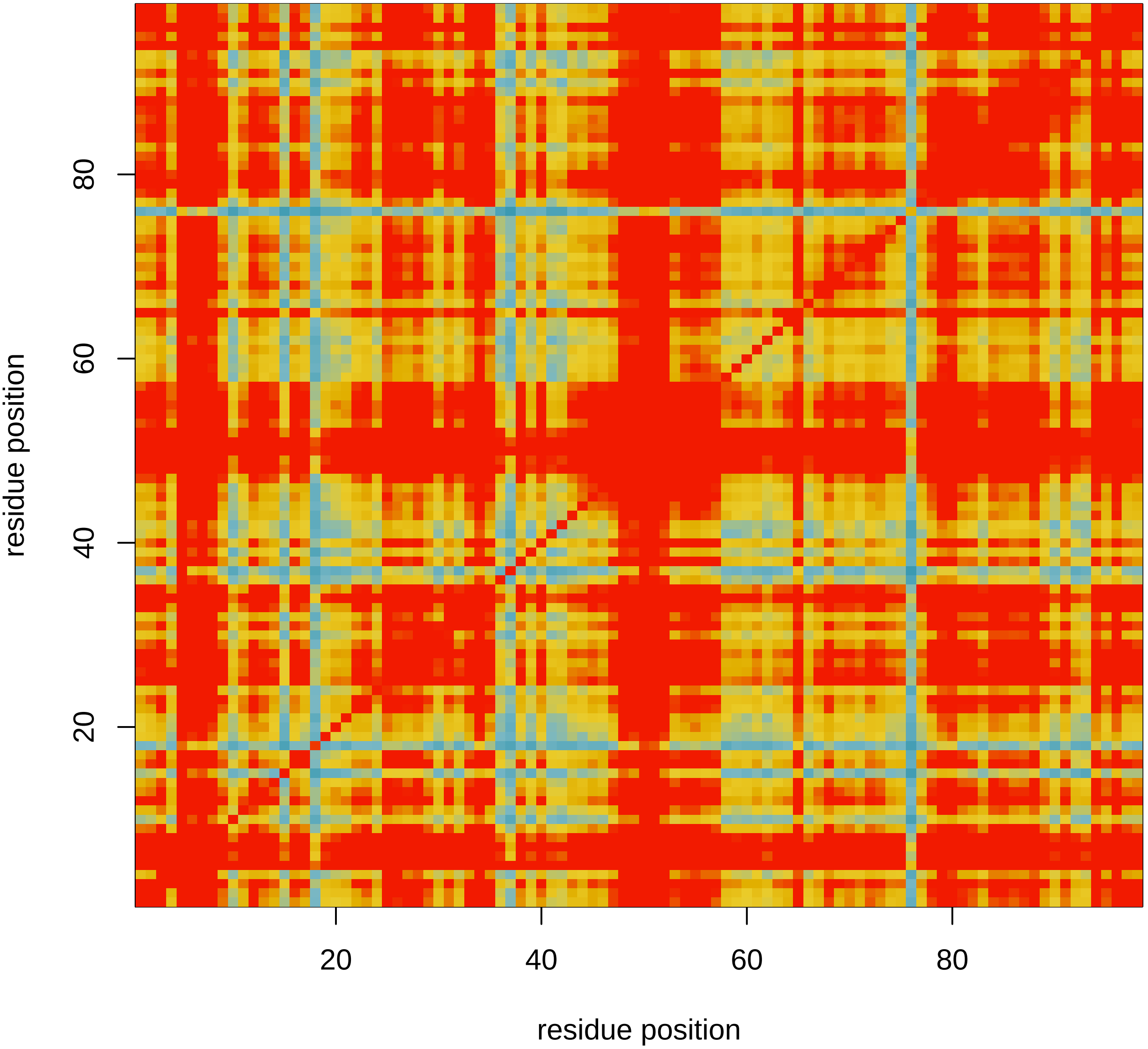

### MImatrix_RFOR_1ytj_monomer_simian.pdf

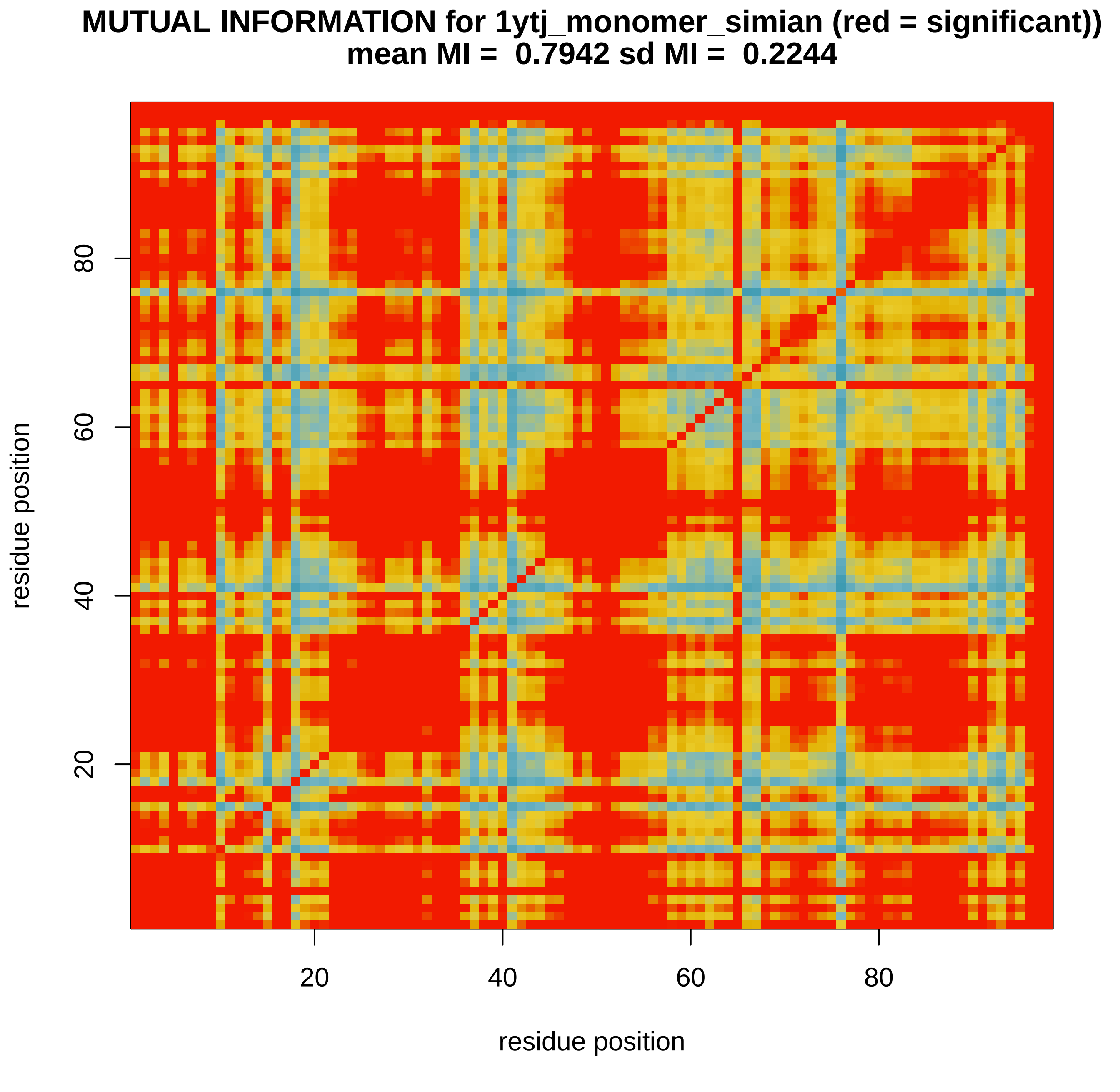

### MImatrix_RFOR_6dgx_monomer_human_1.pdf

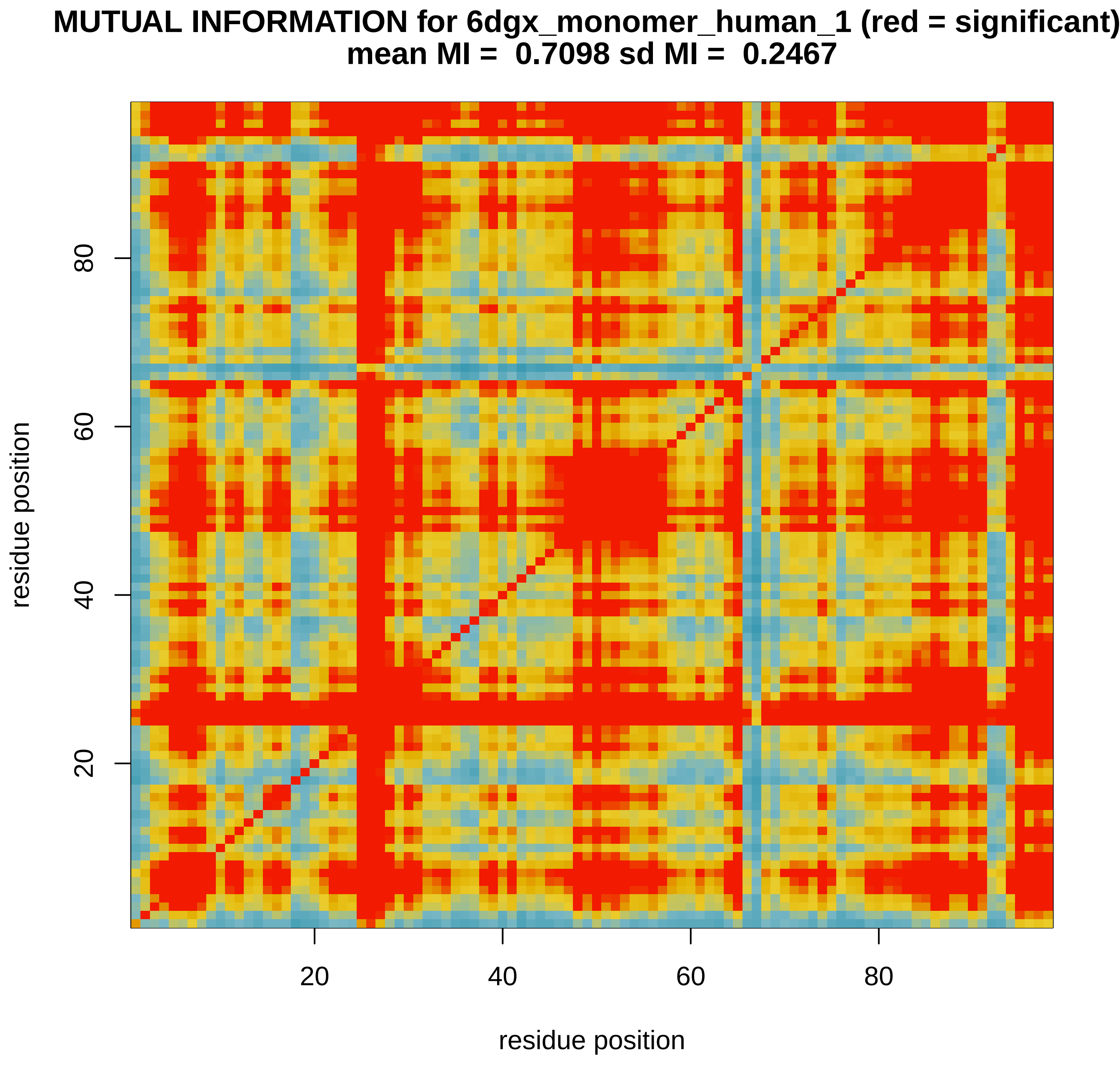

### MImatrix_RFOR_6dgx_monomer_human_2.pdf

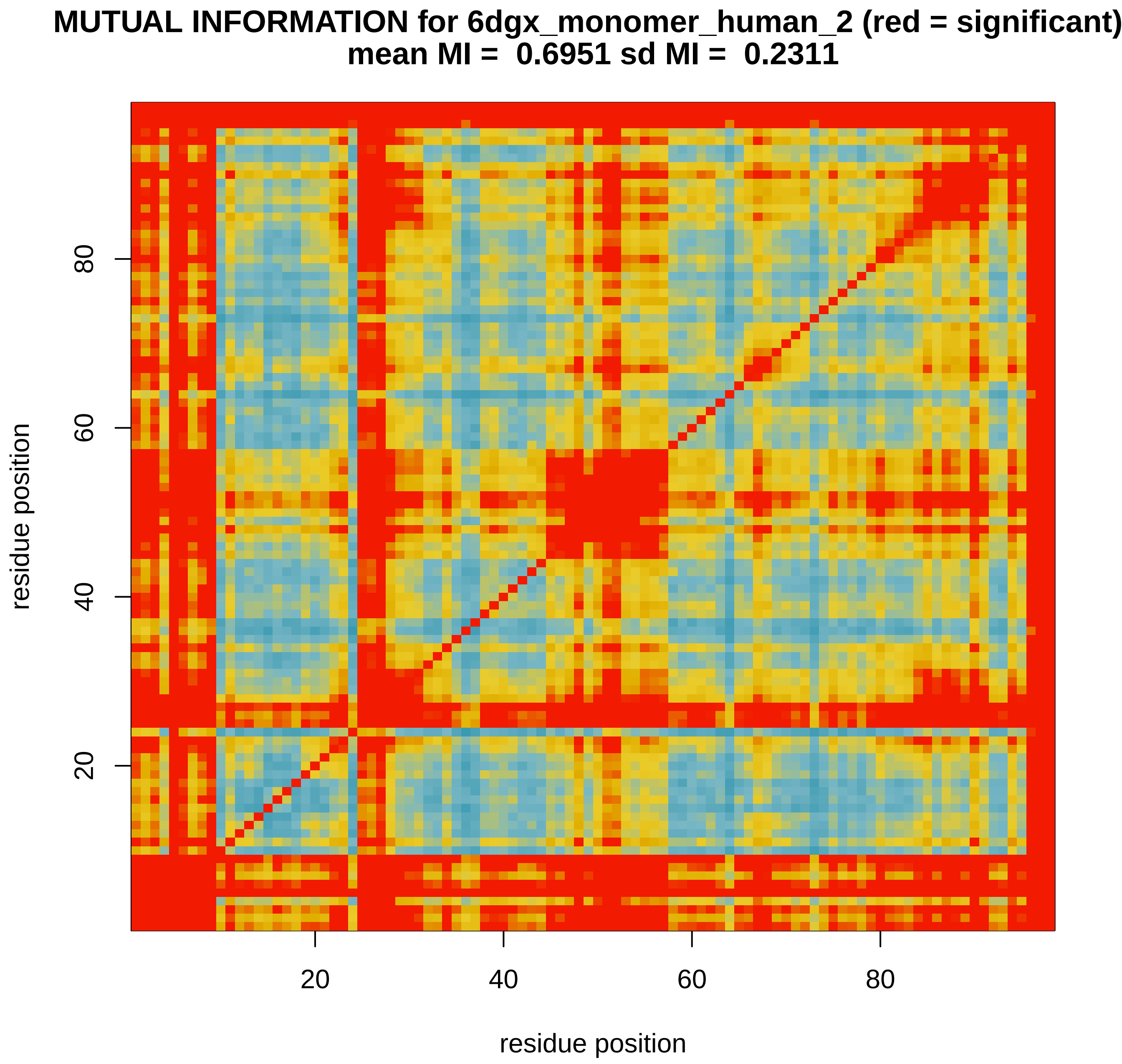

### MImatrix_RFOR_6dgx_nopep_human.pdf

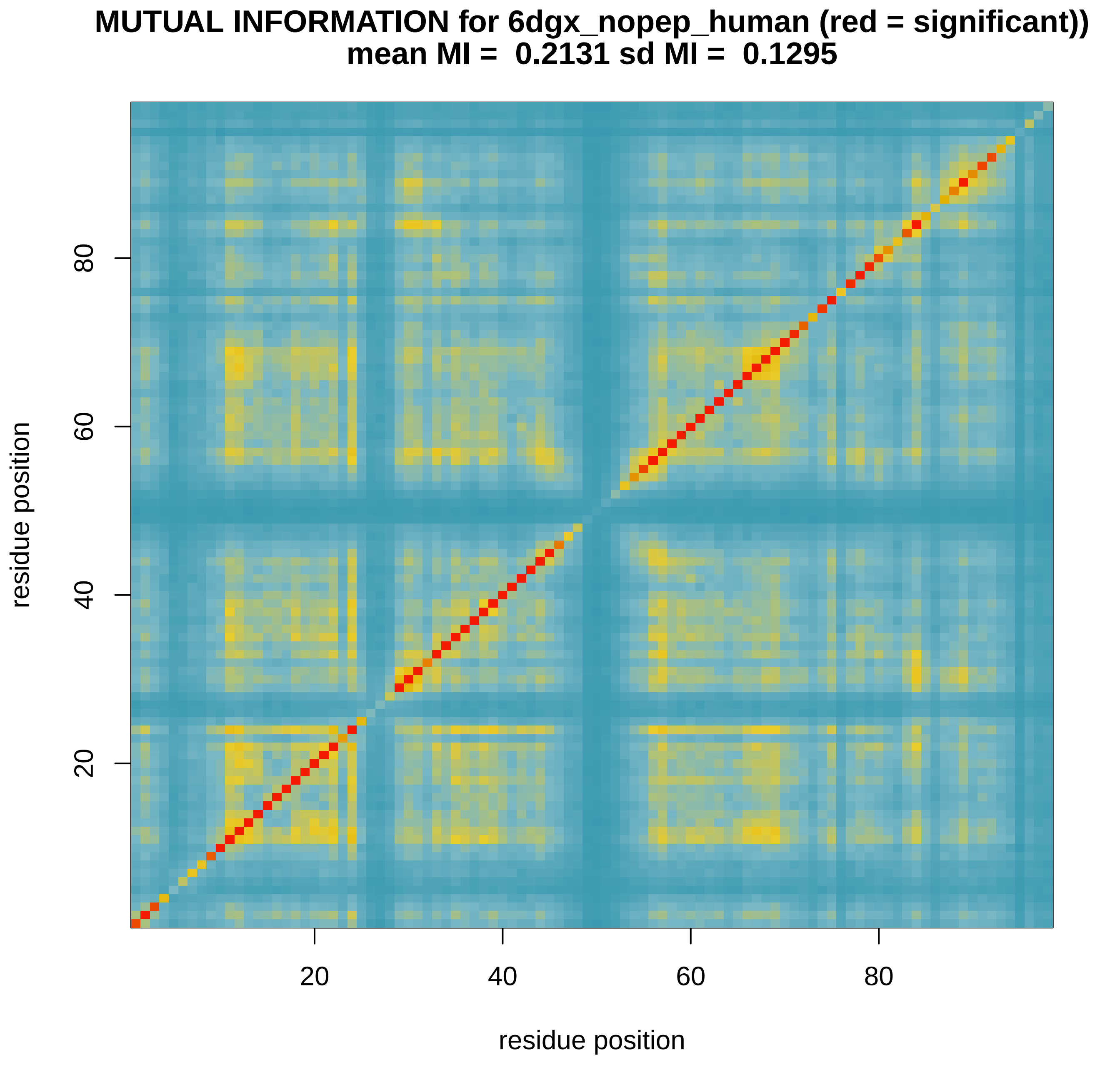

### MImatrix_RFOR_6dgx_nopep_human_1.pdf

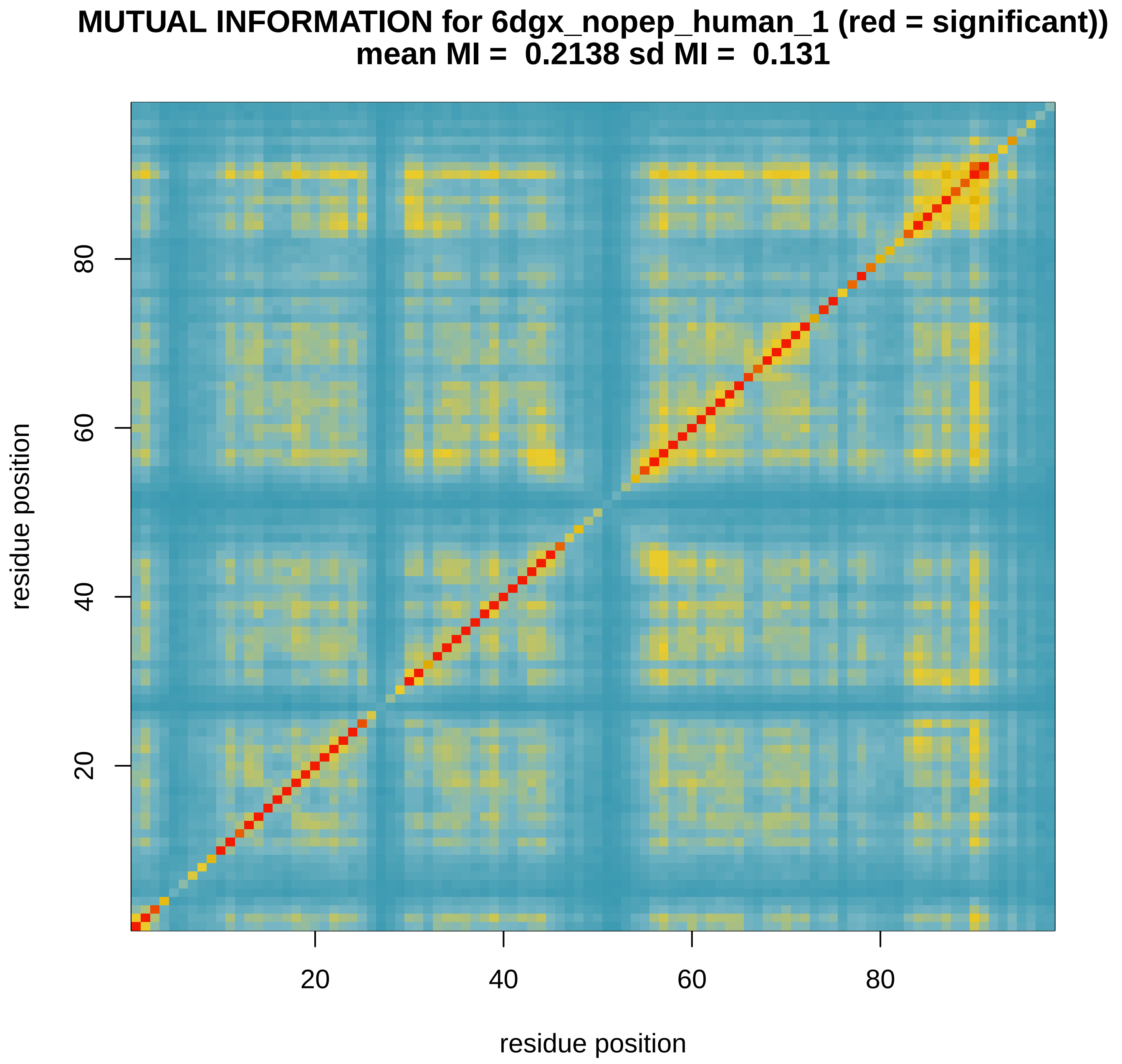

### MImatrix_RFOR_6dgx_nopep_human_2.pdf

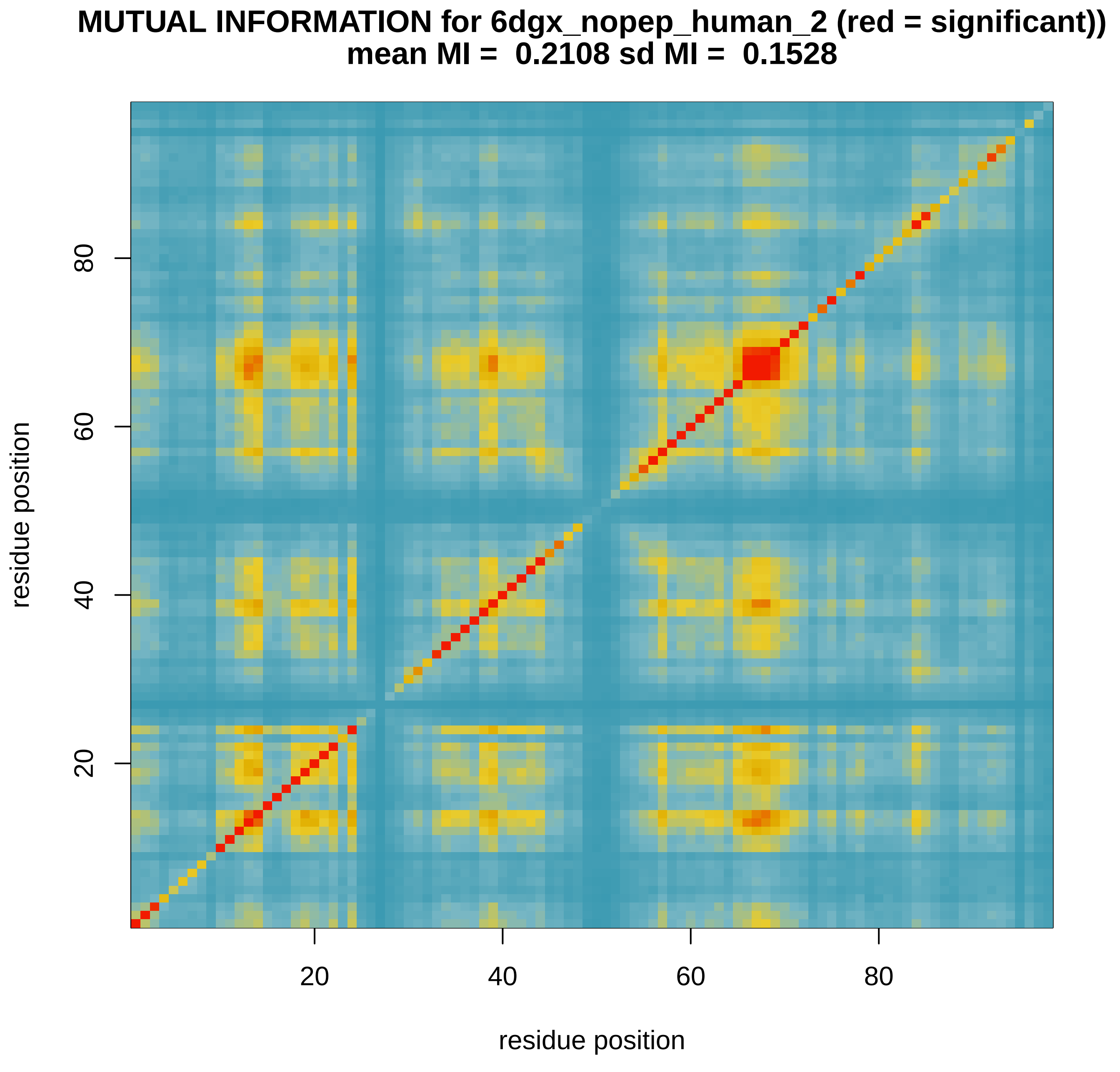

### MImatrix_SVM_1b11_monomer_feline.pdf

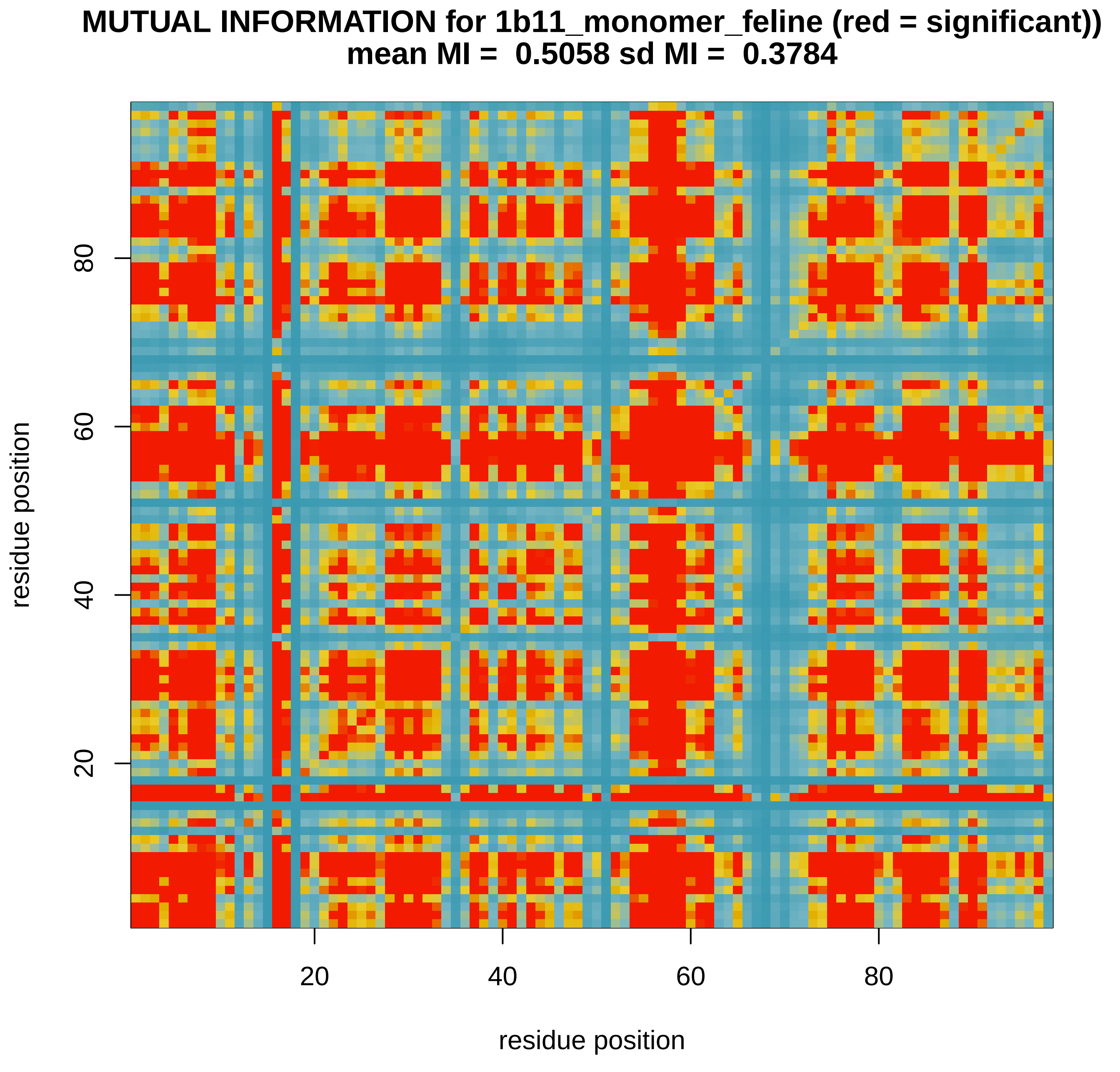

### MImatrix_SVM_1yti_monomer_simian.pdf

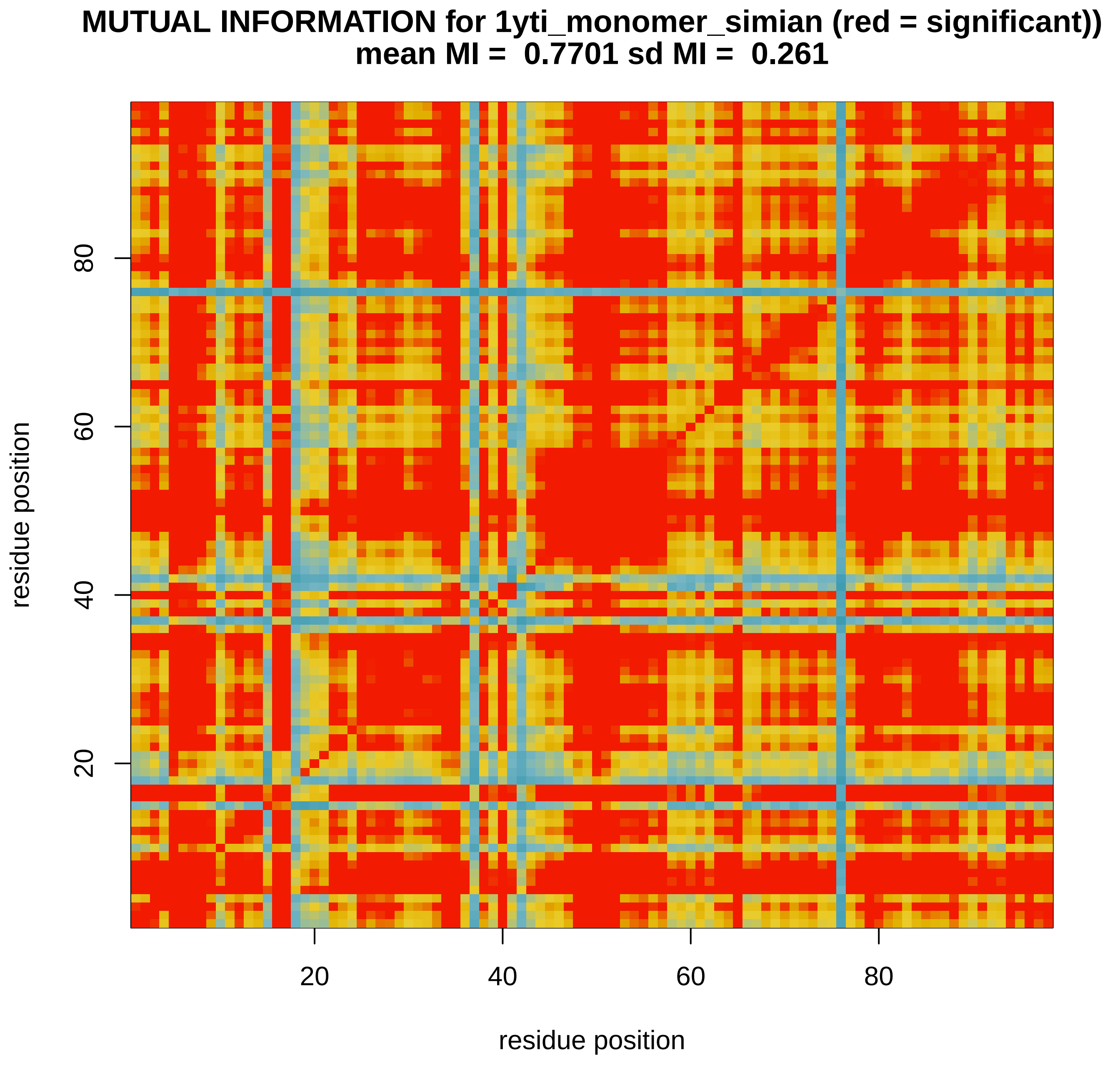

### MImatrix_SVM_1ytj_monomer_simian.pdf

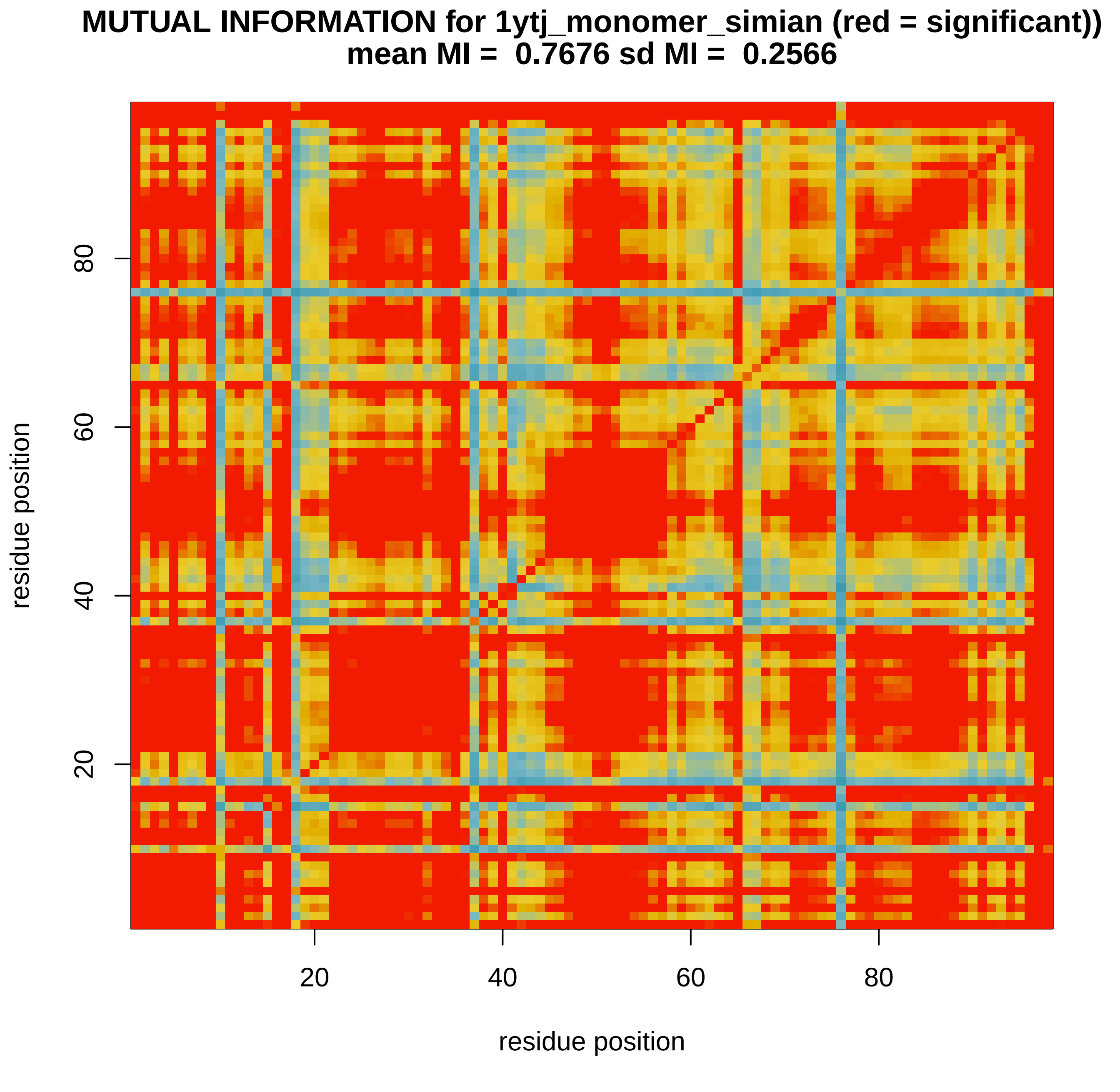

### MImatrix_SVM_6dgx_nopep_human.pdf

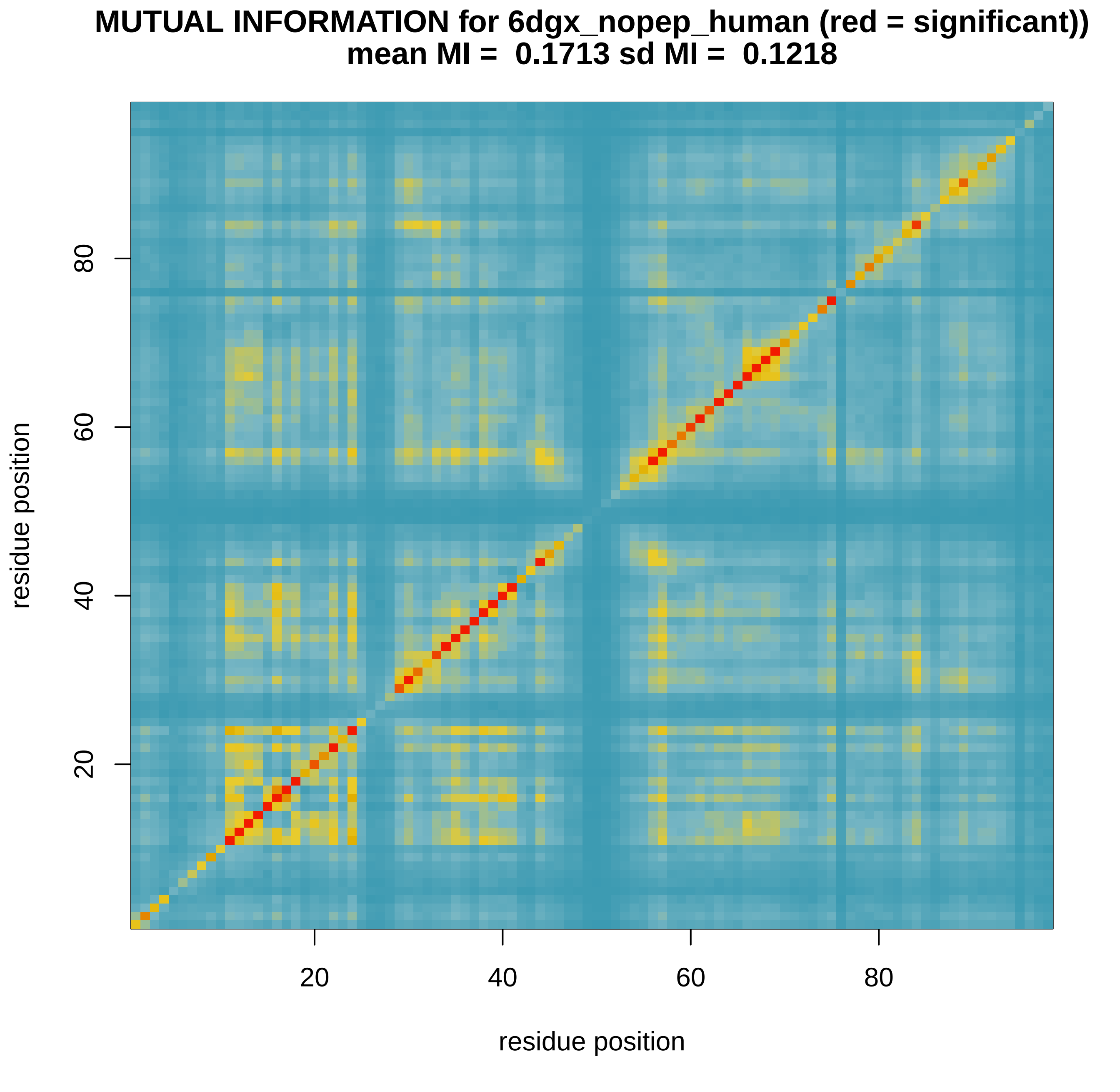

### MImatrix_SVM_6dgx_nopep_human_1.pdf

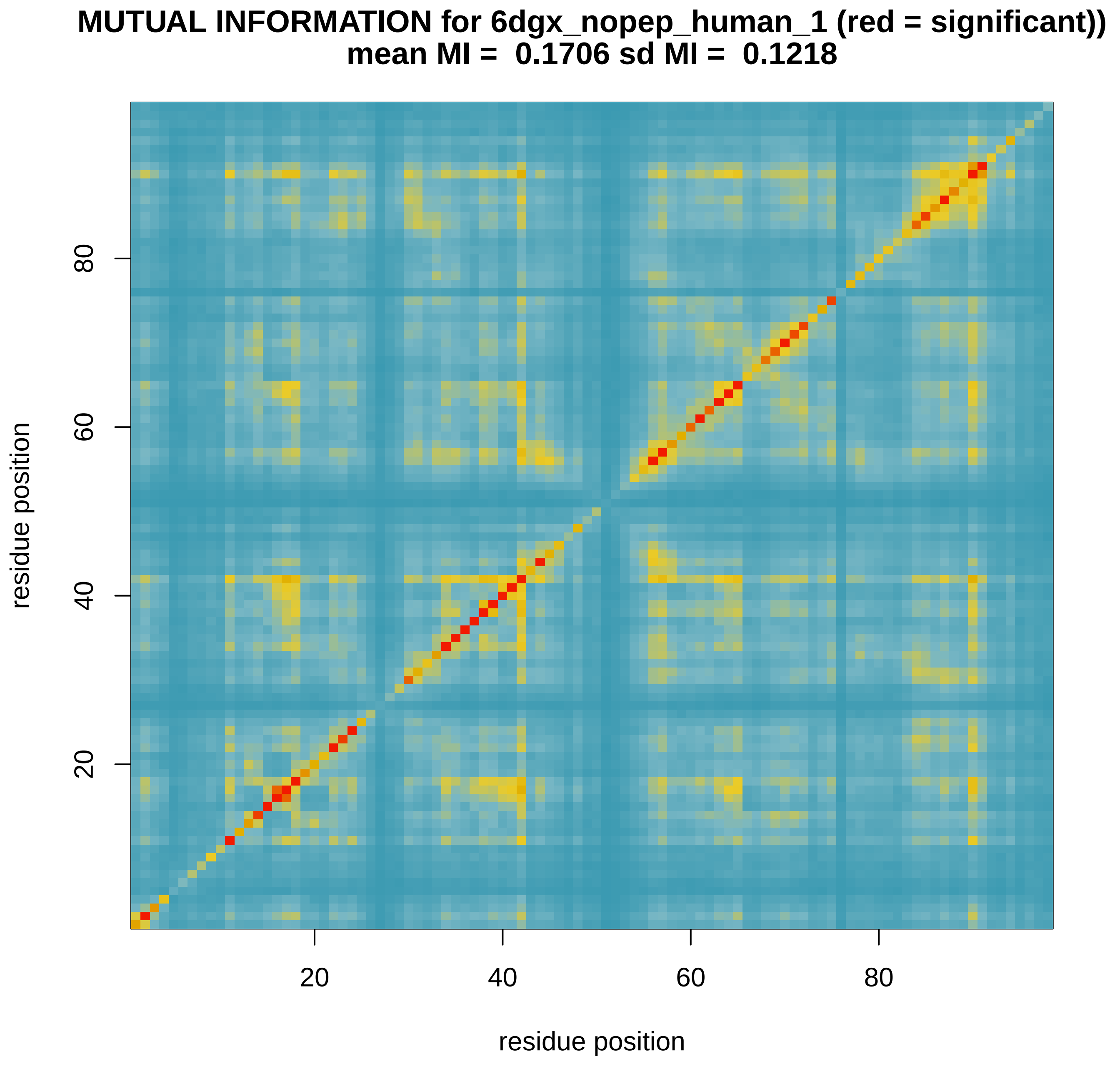

### MImatrix_SVM_6dgx_nopep_human_2.pdf

#### MUTUAL INFORMATION for 6dgx_nopep_human_2 (red = significant))
##### mean MI =  0.1744 sd MI =  0.1431

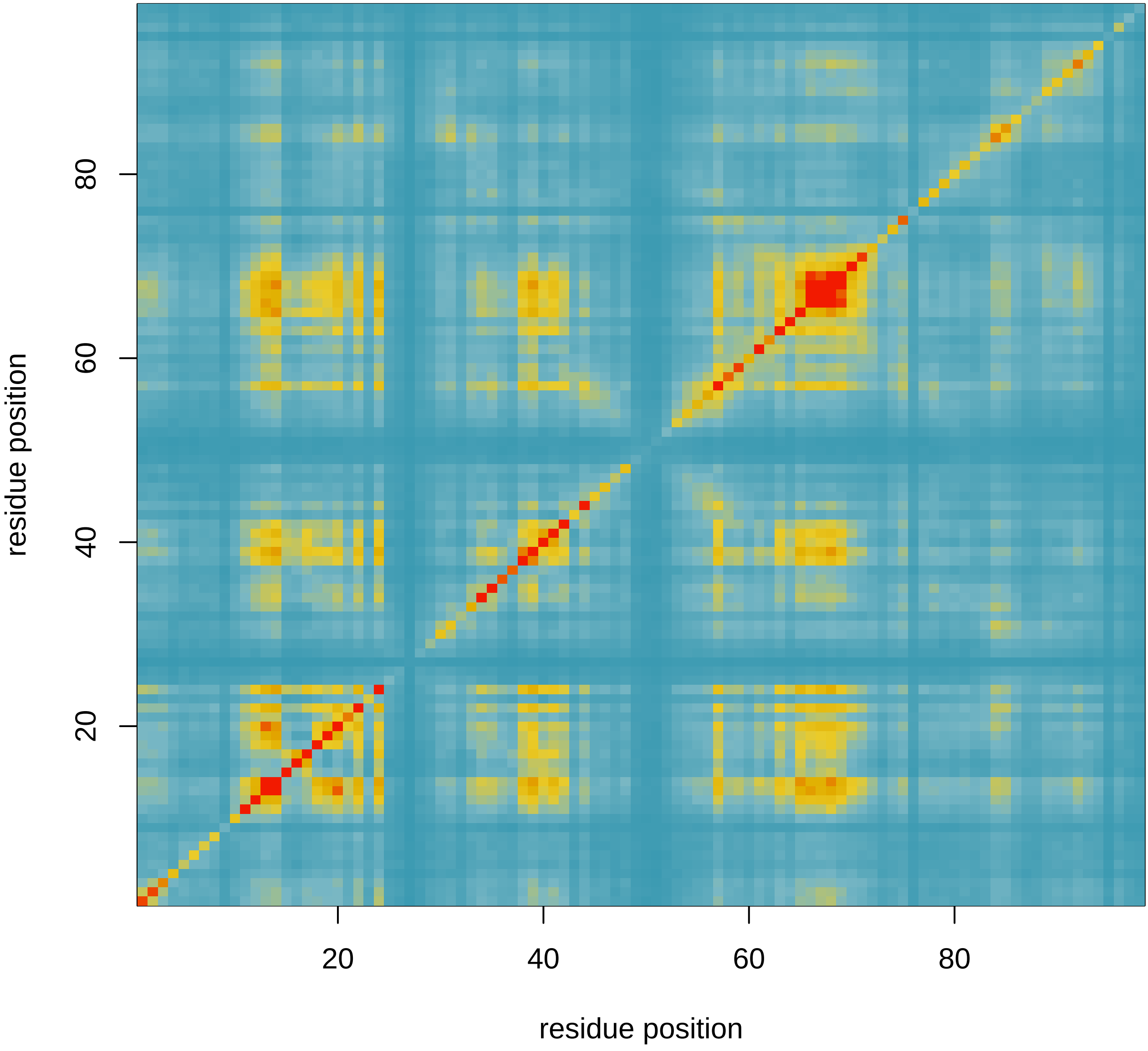
